# Distinct 3D contacts and phenotypic consequences of adjacent non-coding loci in the epigenetically quiescent regions

**DOI:** 10.1101/2023.09.11.557110

**Authors:** Peiyao Wu, Wei Wang

## Abstract

Non-coding regions of the human genome are important for functional regulations, but their mechanisms remain elusive. We used machine learning to guide a CRISPR screening on hubs (i.e. non-coding loci forming many 3D contacts) and significantly increased the discovery rate of hubs essential for cell growth. We found no clear genetic or epigenetic differences between essential and nonessential hubs, but we observed that some neighboring hubs in the linear genome have distinct spatial contacts and opposite effects on cell growth. One such pair in an epigenetically quiescent region showed different impacts on gene expression, chromatin accessibility and chromatin organization. We also found that deleting the essential hub altered the genetic network activity and increased the entropy of chromatin accessibility, more severe than that caused by deletion of the nonessential hub, suggesting that they are critical for maintaining an ordered chromatin structure. Our study reveals new insights into the system-level roles of non-coding regions in the human genome.

## Introduction

The human genome contains many non-coding and non-transcribed regions that have been largely overlooked for their functional importance. Most studies have focused on regions with direct regulatory roles, such as enhancers (Alexander et al., 2010; Blackwood & Kadonaga, 1998; Field & Adelman, 2020), or on genetic variants targeting these elements (Khurana et al., 2016; F. Zhang & Lupski, 2015). However, the annotated genomic regions including protein-coding regions, non-coding RNAs (ncRNAs) and regulatory elements only account for a small fraction of the entire genome and the majority of the genomic DNAs lack detectable transcriptional or epigenetic activity such as transcription factor (TF) binding, open chromatin and histone modifications. These “quiescent” regions cover a large portion of the genome in different cell types (Ernst & Kellis, 2017; Hoffman et al., 2013; van der Velde et al., 2021; Vu & Ernst, n.d.) and their roles in cellular functions remain understudied.

Recently, we analyzed the functional importance of these non-coding loci by focusing on the so-called hub loci, i.e. loci forming many spatial contacts with other loci in the Hi-C experiments (Ding et al., 2021; Dixon et al., 2012; Lieberman-Aiden et al., 2009; Lupiáñez et al., 2015). We performed a CRISPR-based deletion screening to investigate which loci are critical for cell viability (Bock et al., 2022; Jinek et al., 2012; Shalem et al., 2015). We found that 35 out of 960 hubs are essential for cell viability and this percentage (3.6%) of essentiality is comparable to that of ncRNA and protein-coding genes in CRISPR screening (Ding et al., 2021). Interestingly, 27 of these 35 (77%) essential hubs are located in the “quiescent” state, without TF binding, open chromatin, transcription, or histone marks. We further showed that deletion of a quiescent hub could cause global change in chromatin structure, impacting gene expression located millions of base pairs away from the deleted locus in the linear genome. These observations suggest pivotal roles of the “quiescent” regions in maintaining proper 3D genome organization and cellular functions.

This previous study also raised many interesting questions and challenges. For example, the initial study only found 35 essential hubs with a 3.6% hit rate. Because the number of non-coding loci is huge, an efficient strategy to find essential ones particularly those in the quiescent state is desired but remains a challenge. Furthermore, all the 960 hubs analyzed in the CRISPR screening form many 3D contacts, indicating that a large number of spatial contacts is not the only reason for essentiality (Figure S1B). The underlying mechanisms of why only some of these loci are essential remain elusive.

In the current study, we aimed to tackle these challenges. We first implemented a convolutional neural network-autoencoder logistic regression (CNN-AE LR) model (Y. Zhang et al., 2021) based on DNA sequences to select candidates for CRISPR-Cas9 screening. This model captured features associated with essential hubs. Assisted by this model, we selected 340 hubs that can be targeted by CRISPR (∼ 70% in the quiescent region), which significantly improved the discovery rate of essential hubs to 20.6% in the new CRISPR screening. We observed that essential and nonessential hubs exhibited similar sequence and epigenetic features, making it challenging to distinguish them. By assuming that the spatial interactions might be responsible for hubs’ essentiality difference, we analyzed one essential and one nonessential hub that are adjacent to each other in the linear genome. Surprisingly, these hubs formed distinct spatial communities. Subsequent single cell RNA-seq and ATAC-seq experiments on the hub-deleted cells showed that cell death could not be attributed to a specific set of genes or pathways but rather to the enhanced activity of the genetic network in the essential hub-deleted cells. Additionally, Hi-C experiments on the hub-deleted cells demonstrated that hub deletions caused the loss of many contacts in the wild type (WT). Interestingly, the deletion of the essential hub induced a higher number of new weaker contacts compared to the nonessential hub. Notably, many of these newly formed contacts overlapped with the H3K27ac marked regions in WT and were associated with the highly expressed genes upon essential hub deletion. Based on the evidence suggesting that hubs are critical for maintaining the order of chromatin organization, we proposed to quantify the impact of hub deletion using Shannon entropy on open chromatin signals. We found that hub deletion resulted in increased entropy compared to the WT, and a larger increase was induced by deleting the essential hub than the nonessential one.

## Results

### Rational selection of hub candidates for screening by a convolutional neural network-autoencoder logistic regression model (CNN-AE LR)

In our previous study (Ding et al., 2021), we discovered a correlation between the appearance of genomic variations (GVs) in hubs and significant change in 3D contact numbers, highlighting the importance of DNA sequences in hub formation. To further investigate this relationship, we examined whether hubs share any sequence patterns. Using 5-kb sequences of all 26,148 hubs including the 960 analyzed in Ding et al. study in the K562 cell line, we developed a convolutional neural network-autoencoder (CNN-AE) model (Y. Zhang et al., 2021) (Figure 1A) with two main components: (1) an encoder consisting of three convolution-max pooling blocks, and (2) a decoder with three transposed convolution blocks. The input hub sequence was one-hot encoded, and transformed into a latent representation vector by the encoder, and then reconstructed to the input sequence by the decoder. We ensured the convergence and no overfitting of the model in 10 independent runs (Figure S2A and Methods and Materials for additional details).

**Figure 1.**
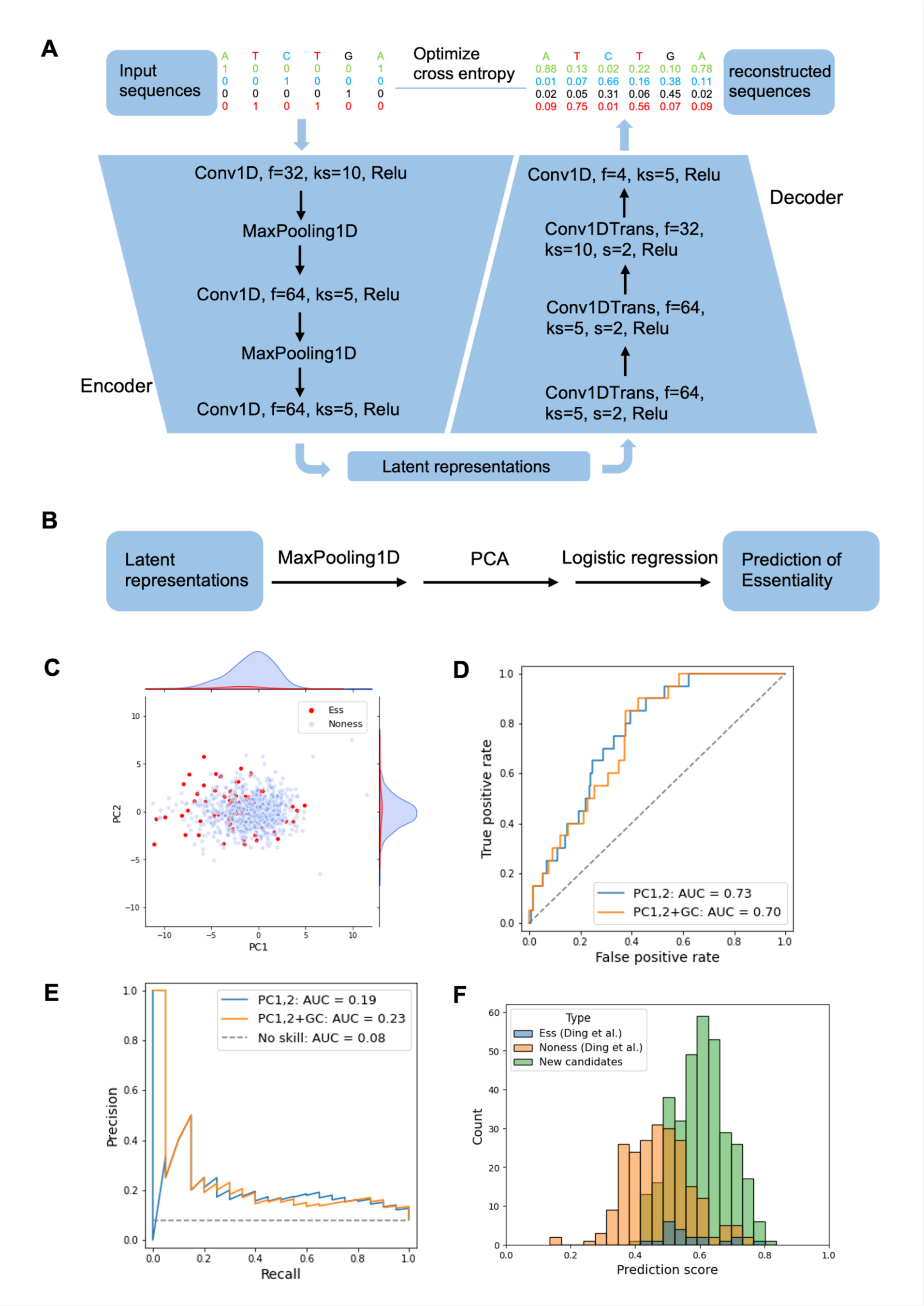
The CNN-AE model. **(A)** Architecture of the convolutional neural network autoencoder (CNN-AE) designed to learn the sequence grammar of all the hubs defined in Ding et al. **(B)** Architecture of the logistic regression (LR) model used to classify hub essentiality based on the sequence latent representations obtained from the CNN-AE. **(C)** PC plot showing the latent representations of the sequences of all the hubs tested in Ding et al. Essential hubs tend to have smaller PC1 values, which serves as the basis for the LR model. **(D)** Receiver operating characteristic (ROC) curve illustrating the performance of the LR models using the top2 PCs and GC content as features. **(E)** Precision-recall curve representing the performance of the LR models using the top2 PCs and GC content as features. **(F)** Visualization of the new screening candidates with a prediction probability score cutoff >= 0.4.

After performing max pooling (size = 25) on the sequence latent representation, we conducted a principal component analysis (PCA) on all the hub sequences (Figure 1B) and observed that the essential hubs identified from the Ding et al. study (Ding et al., 2021) were mainly located in the area with smaller PC1 (Figure 1C and S2C). We next utilized the top 2 PCs which explained 7.1% of the sequence variance to build a logistic regression (LR) model (Figure 1B) and it achieved an area under the ROC curve (ROC-AUC) of 0.73 (Figure 1D). Moreover, the area under the precision-recall curve (PR-AUC) was 0.19 (Figure 1E), which outperformed a no-skilled random guess by a factor of 2.38.

Despite the limited size of our dataset, particularly for the essential loci, this model provided guidance for selecting candidates for the second round of screening based on the following criteria (Figure 2A). We began with the hubs in K562 cells defined by (Ding et al., 2021), among which 25,188 were not included in the previous screening. We mapped these hub sequences to the PCs of the pooled latent representations and predicted their essentiality using the LR model. We removed the sites with low predicted scores using a threshold of 0.4 (Figure 1F). We also removed loci with GC content < 0.4 since we found that the essential loci have higher GC content than the nonessential loci in the first round of screening (see Figure S1). To focus on the quiescent state, we next removed the loci overlapping with any ChIP-seq peak of 7 active histone marks (H3K4me1, H3K4me2, H3K4me3, H3K9ac, H3K27ac, H3K36me3, and H4K20me1) or 3 transcription factors (TFs) important for chromatin looping (CTCF, RAD21, and SMC3) in K562. We further removed sites that cannot be targeted by at least 6 pgRNAs composed of stringent sgRNAs with high specificity and low off-target rate (see Methods and Materials for details). Finally, to accommodate the size limit of the oligo synthesis, we randomly selected 340 hubs for screening in this study: 70% are quiescent regions without any annotations and the remaining are marked by either the repressive H3K9me3 or H3K27me3 modification for assessing the importance of these marks on hub essentiality.

**Figure 2.**
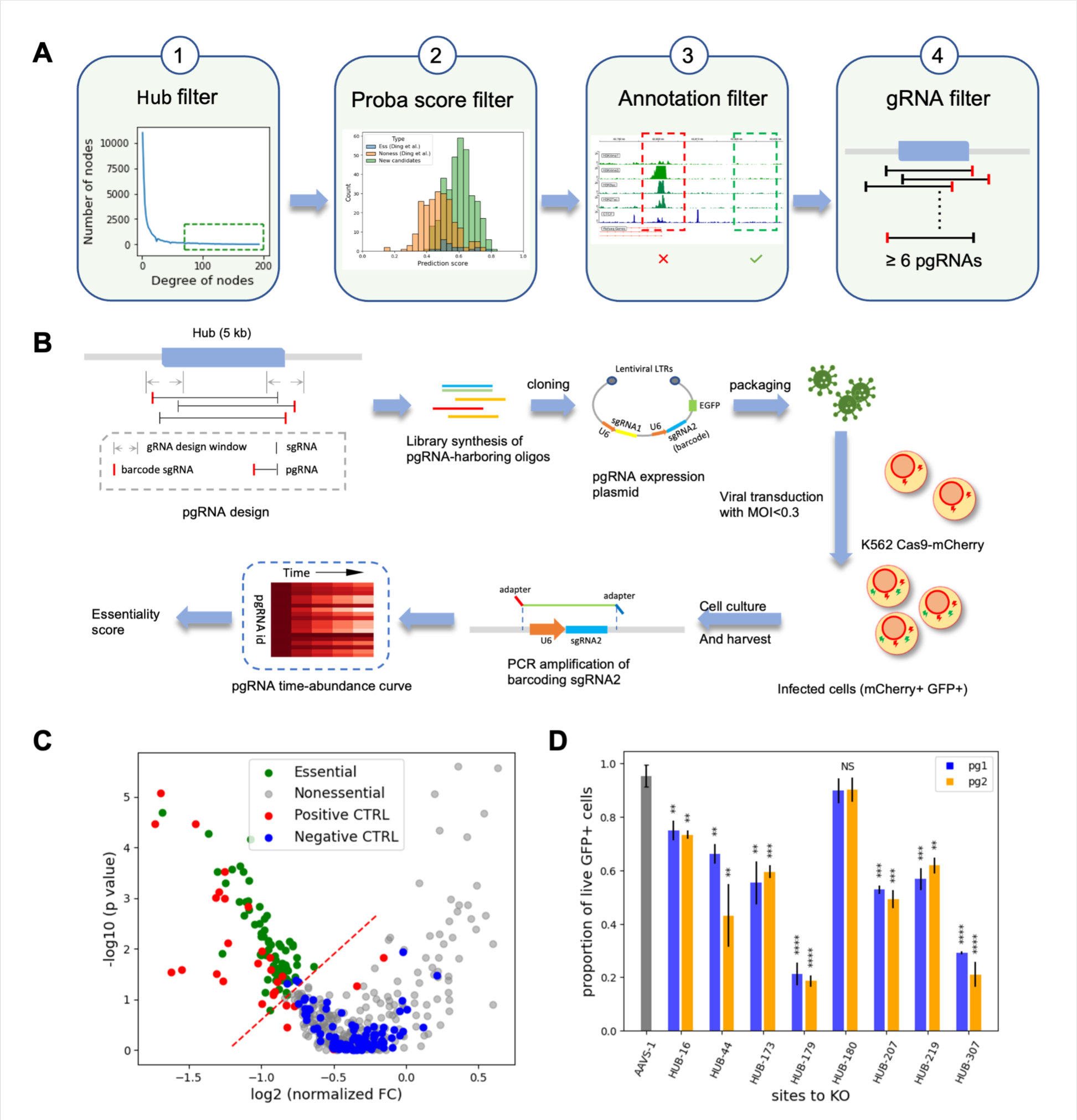
CRISPR-Cas9 screening experiments. **(A)** Schematic of rational selection process for hub candidates. Step 1: We chose hubs that form many 3D contacts in the Hi-C experiments (*Z*-score > 2, indicated by the green box). Step 2: We filtered the hubs based on the prediction probability scores (>= 0.4) obtained from the CNN-AE LR model and the GC content (>= 0.4). Step 3: We removed hubs overlapping with ChIP-seq peaks of major histone modifications, CTCF, RAD21, SMC3,or RefSeq signals (e.g. the hubs in the red box were removed and the ones in the green box remained). Step 4: We kept hubs that could be targeted by at least 6 eligible pgRNAs. **(B)** Schematic of the pgRNA design, library construction, and the screening process to assess cellular viability of the selected hub candidates. **(C)** Volcano plot of log2(normalized *FC*) versus -log10(*p*-value). The dashed red line represents the boundary for an *I* score = -1. Hub candidates above the line with *FC* < 1 were identified as essential for cell viability (green). **(D)** Barplots of the cellular viability upon hub deletion compared with AAVS-1 site deletion. All tested hub sites except HUB-180 caused significant decreases in GFP%, which validated the screening results. Data are presented as mean ± standard deviation (n = 2). * *p* < 0.05, ** *p* < 0.01, *** *p* < 0.001, **** *p* < 0.0001, and NS means not significant. All *p*-values were calculated by comparing the proportion of live GFP+ cells in the hub deletion samples with that of AAVS-1 site deletion samples, using a student’s t test. Each site was targeted by two individual pgRNAs.

### A significantly improved discovery rate of essential hubs achieved in CRISPR-Cas9 screening

We performed CRISPR-Cas9 screening in K562 cells following the protocols in Ding et al. study (Ding et al., 2021). In short, we designed a library of 6,449 pgRNAs targeting 340 hub loci (20 pgRNAs for most of the loci), 473 targeting the first exon regions of 29 essential ribosomal genes as positive controls, 100 targeting the *AAVS1* locus and 100 that do not target human genome as negative controls. We constructed the pgRNA library into a lentiviral-expression system and transduced it into K562 cells stably expressing Cas9-mCherry, with a controlled multiplicity of infection (MOI) of 0.3. Cells taking in the lentivirus were sorted on the third day post infection (marked as day 0 of the screening), and cultured for another 25 days. We harvested the cells every 5 days and determined the abundance of the pgRNAs remaining in the cells by genotyping the barcoding-gRNA region and sequencing (Figure 2B).

To achieve a reliable screening result, we selected pgRNAs with high specificity scores to minimize off-target effects (Morgens et al., 2017; Tycko et al., 2019) (Figure S3A), and achieved a high coverage of 200-fold and a uniform representation of all pgRNAs for effective and unbiased cleavage (Figure S3B, and see Methods for additional details). The read counts of the pgRNA were highly correlated between two biological replicates, with Pearson correlation coefficients of 0.893 and 0.882 for day 0 and day 25, respectively (Figure S3C and D). To determine hub essentiality, we first calculated the fold change for each pgRNA (*FC_guide_*) between day 25 and day 0. We next log-linearly normalized the *FC_guide_* values in this round of screening to ensure comparability with the first round screening in the Ding et al. study (Ding et al., 2021) (see Methods and Materials, Figure S4). The distributions of the normalized *FC_guide_* values for the control sites were indistinguishable in the two rounds (*p* = 0.816 for positive controls, and *p* = 0.979 for negative controls using Mann-Whitney U test), validating the normalization. We then averaged the *FC_guide_* for each locus to obtain *FC*_site_. To derive a *p*-value for each site, we used Mann-Whitney U test to compare the distributions of *FC_guide_* targeting each site (with 6 ∼ 20 pgRNAs per site) to that of the negative controls.

Smaller *FC*_site_ and *p*-values indicate that a hub is essential for cell viability. To combine the two scores into one metric, we derived *I* scores as previously defined (Ding et al., 2021): *I_site_* = *sign*(*Z_F,site_*) * ||*Z_F,site_*| + *Z_P,site_* |, where *Z_F,site_* is the *Z* scale of log_2_(*FC_site_*) of a given site, and *Z_P,site_* is the *Z* scale of its -log_10_(*p*-value). As a result, choosing the same cutoff of Isite<= -1 as the first round, we identified 70 hubs that are essential for the K562 cell line, achieving a much higher hit rate of 20.6% compared to 3.6% in the first round of screening in Ding et al. study (Table S4 and Figure 2C). Among the essential hubs, 50 are in the epigenetically quiescent regions (71.4%). These results suggest that the hub selection process successfully captured some features associated with essentiality. Furthermore, we found no significant difference in the percentage of essential hubs between those in the quiescent state and those marked by the repressive mark of H3K9me3 or H3K27me3.

We validated the screening results by individually testing 8 selected loci (see Methods), including 4 “strongly essential” hubs with extremely negative *I_site_* scores (< -4.0) and 4 “weakly essential” hubs with *I_site_* scores closer to the cutoff. For each hub, we selected two pgRNAs with highest specificity scores from the library and cloned them into individual lentiviral-expression plasmids. We performed proliferation assays to assess the effects of hub deletion on cell growth in comparison to AAVS1 deletion (Figure 2D and S5). We confirmed that all the hubs, except one “weakly essential” hub (HUB-180), were critical for cell survival and growth. These individual validations demonstrated that the screening results and the selected cutoffs were effective in detecting essential hubs.

### Indistinguishable sequence and epigenetic features but different 3D interacting communities between essential and nonessential hubs

We next attempted to investigate whether hub essentiality is associated with any sequence or epigenetic features. We began with GC contents and found that essential and nonessential hubs had comparable GC contents in the current screening (Figure S6A, *p* = 0.430 by Mann-Whitney U test). We also performed motif analysis using Homer (Heinz et al., 2010) and found no enrichment of known or *de novo* motifs in the essential hubs (see Methods). We then examined the genomic distribution of the hubs and observed no clustering or separation of essential and nonessential hubs in the linear genome (Figure S6C).

To further explore the potential epigenetic features associated with hub essentiality, we built a logistic regression classifier using sequence information and various epigenetic features (Figure 3A). The sequence information was derived from the top principal components (PCs) of the latent representations obtained from the trained CNN-AE. For the epigenetic features, we took the fragment contact network (FCN) constructed in Ding et al. study (Ding et al., 2021) and calculated the personalized Pagerank for each hub on each feature, which reflects its global importance in the FCN (see Methods). In short, we defined the 5-kb genomic fragments in Hi-C data (Rao et al., 2014) as nodes, and 3D contacts with *p*-values <= e^-20^ as edges. For each epigenetic feature, we set the node weight as 1 if the node overlaps with the peak of the epigenetic feature such as a H3K27ac peak and 0 otherwise. We computed the *Z* scale of the PageRank score in the FCN for each node on each epigenetic feature using the node weights as a personalization (see Methods for more details about “FCN construction” and “feature preprocessing”). We considered the following features: GC content, top 30 sequence principal components (PCs), as well as the PageRank scores of (i) nine histone modifications (H3K4me1, H3K4me2, H3K4me3, H3K9ac, H3K9me3, H3K27ac, H3K23me3, H3K36me3, H4K20me1), (ii) three TFs (CTCF, SMC3, RAD21), (iii) lncRNAs, and (iv) essential genes. The t-SNE analysis of these features (Figure 3B) did not show a clear separation between essential and nonessential hubs. We also tried different combinations of features for the logistic regression classifier, but none of them achieved an accuracy better than 0.6 (Figure S6B), indicating that hub essentiality is not easily predictable from sequence or epigenetic features alone.

**Figure 3.**
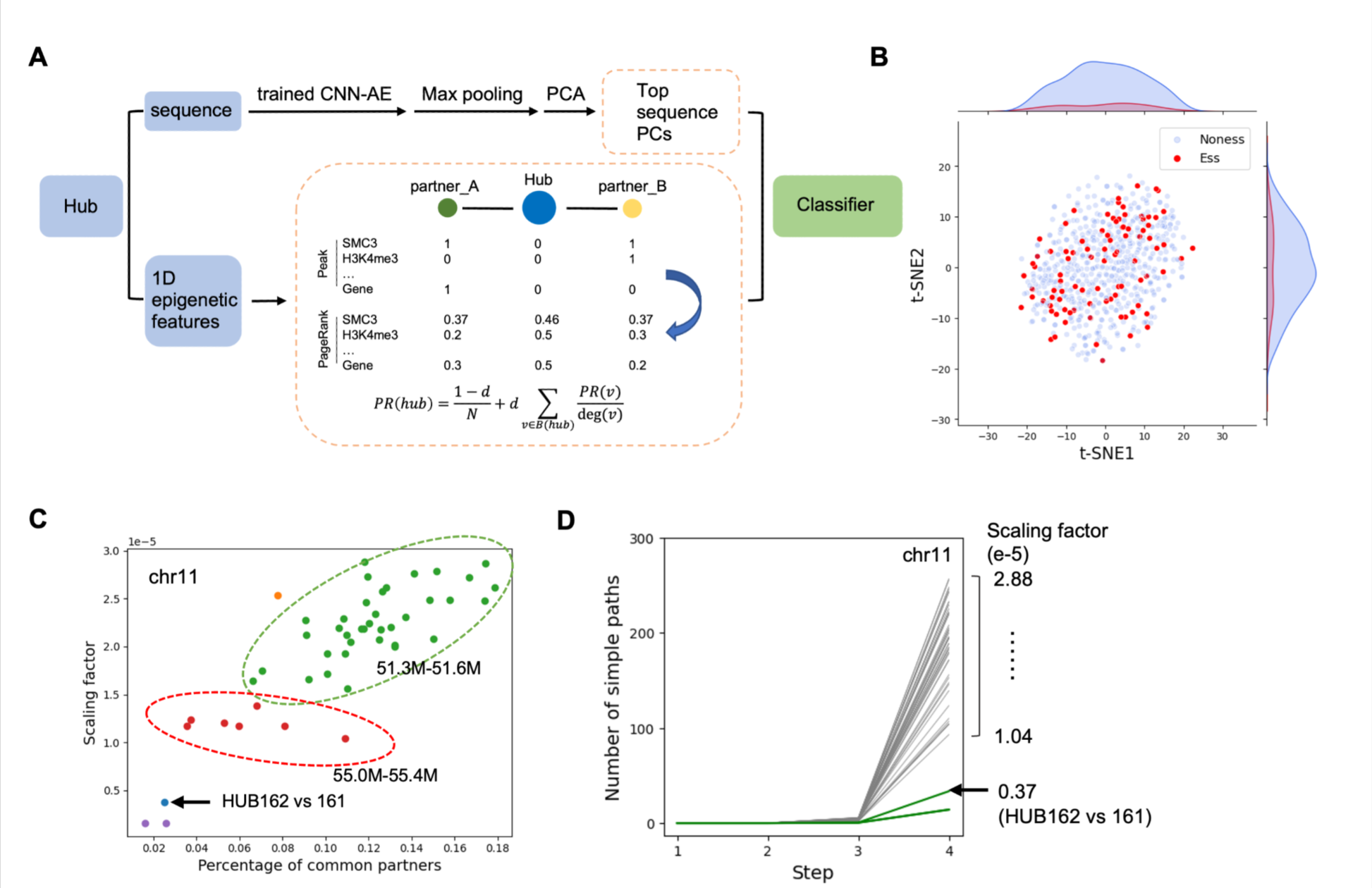
Essential and nonessential hubs are spatially different and cannot be distinguished by simple 1D epigenetic features. **(A)** Schematic of the construction of a classifier for hub essentiality. Sequence features: Top Sequence PCs were derived from hub sequences with a CNN-AE, max-pooling, and PCA. 1D epigenetic features: FCN was constructed as described in Methods (Ding et al., 2021); ChIP-seq peaks of histone modifications, TF bindings, and gene count reads were mapped to nodes; for each feature channel, the personalized PageRank was performed, and centrality was calculated as the *Z*-scale of its PageRank score. **(B)** t-SNE plot showing no clear difference between essential and nonessential hubs. Features including Centrality scores of (H3K4me1, H3K4me2, H3K4me3, H3K9ac, H3K9me3, H3K27ac, H3K23me3, H3K36me3, and H4K20me1, CTCF, SMC3, RAD21, lncRNAs, and essential genes), top 30 sequence PCs, and GC content (Methods). **(C)** Path analysis with step >= 2, i.e. common partners and scaling factors. The plot shows hub pairs on chr11. Pairs located in different regions tend to have varying percentages of common partners and scaling factors. More examples can be found in Figure S7C. **(D)** Path analysis with up to 4 steps, i.e. dense connectivities. The number of simple paths is exponentially correlated with the number of steps, and the scaling factor indicates how densely the pair is connected. The plot shows hub pairs on chr11. Most pairs have scaling factors between 1.04 and 2.88e-5, while certain hub pairs, including HUB162-HUB161 (0.37e-5), have smaller scaling factors (refer to scaling factors for all hub pairs in Figure S7D).

As we did not observe distinct sequence or epigenetic features when comparing all the essential and nonessential hubs, we hypothesized that the difference in essentiality might originate from their spatial interactions. We therefore searched for hub pairs located within <= 50 kb in the linear genome and found 4 Essential-Essential pairs, 396 Nonessential-Nonessential pairs, and 135 Essential-Nonessential pairs, i.e., neighboring hub pairs that have opposite essentialities (Table S7).

We examined how closely these hub pairs were connected in the FCN by analyzing the number of simple paths (i.e. paths without repeating nodes) connecting them. As one-step path indicates direct connection, it is not surprising that the closer the hub pairs are located in the linear genome, the more likely they form direct connections (Figure S7A and B). If two hubs are connected by a two-step path, they share “common neighbors” and thus the number of two-step paths reflects the density of 3D contacts between the hub pairs. We normalized these values against the geometric mean of the hubs’ degree numbers and found that they varied in a broad range from 0 to > 0.7. For example, hub pairs located within chr11:51.3M-51.6M (hg19) had values of 0.066 ∼ 0.179, suggesting relatively denser regional connectivity, compared to 0.035 ∼ 0.109 of those located within chr11:55.0M-55.4M (Figure 3C and S7C). As the step number *n* increases, the number of simple paths *D* increases exponentially (i.e. fitting to *D = Ae^Bn^ + C*, where *A*, *B* and *C* are fitting parameters, see Methods) and the scaling factor *A* suggests how densely the two hubs are connected. When the step number is 4, the scaling factors of hub pairs drastically diverge (Figure 3D and S7C). When analyzing the scaling factors of all possible pairs among all 26,148 hubs, we found some genomic regions having very low scaling factors, suggesting local segregation of chromatin structure (Figure S7D). Interestingly, some Essential-Nonessential pairs are located in these regions. This observation is striking particularly because some of these hubs are adjacent regions in the linear genome. Distinct 3D contacts with opposite essentialities made these hub pairs interesting candidates for the follow up studies.

### Differential spatial interactions of adjacent hubs with opposite essentialities

We selected one of these pairs located in the quiescent region (Figure 4A and Methods) and conducted in-depth analysis. The two loci are HUB-162 (chr11:42,878,450-42,883,450, hg38), which is an essential hub with a negative *I* score of -1.01, and its immediate genomic neighbor, HUB-161 (chr11:42,873,450-42,878,450, hg38), which is a nonessential hub with a positive *I* score of 2.16, and only has minimal effects on cell growth when deleted. We first performed individual validation assays using 3 different pgRNAs targeting each locus to confirm their difference on essentiality (Figure 4B, C, and Table S6). We further demonstrated that this difference is not resulting from different pgRNA cutting effectiveness, as the deletion efficiencies for both hubs measured by qPCR were comparable at every time point (Figure S9).

**Figure 4.**
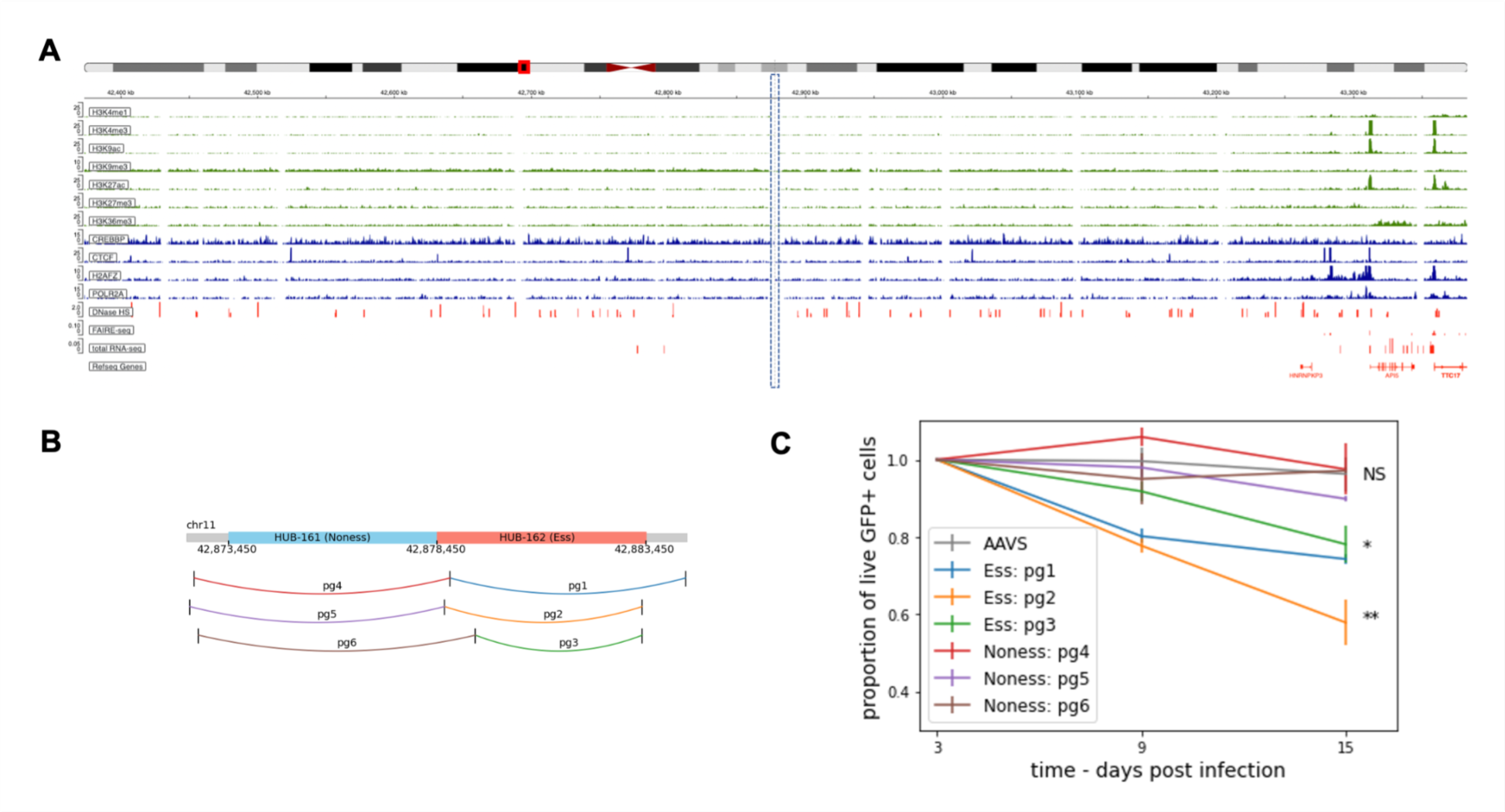
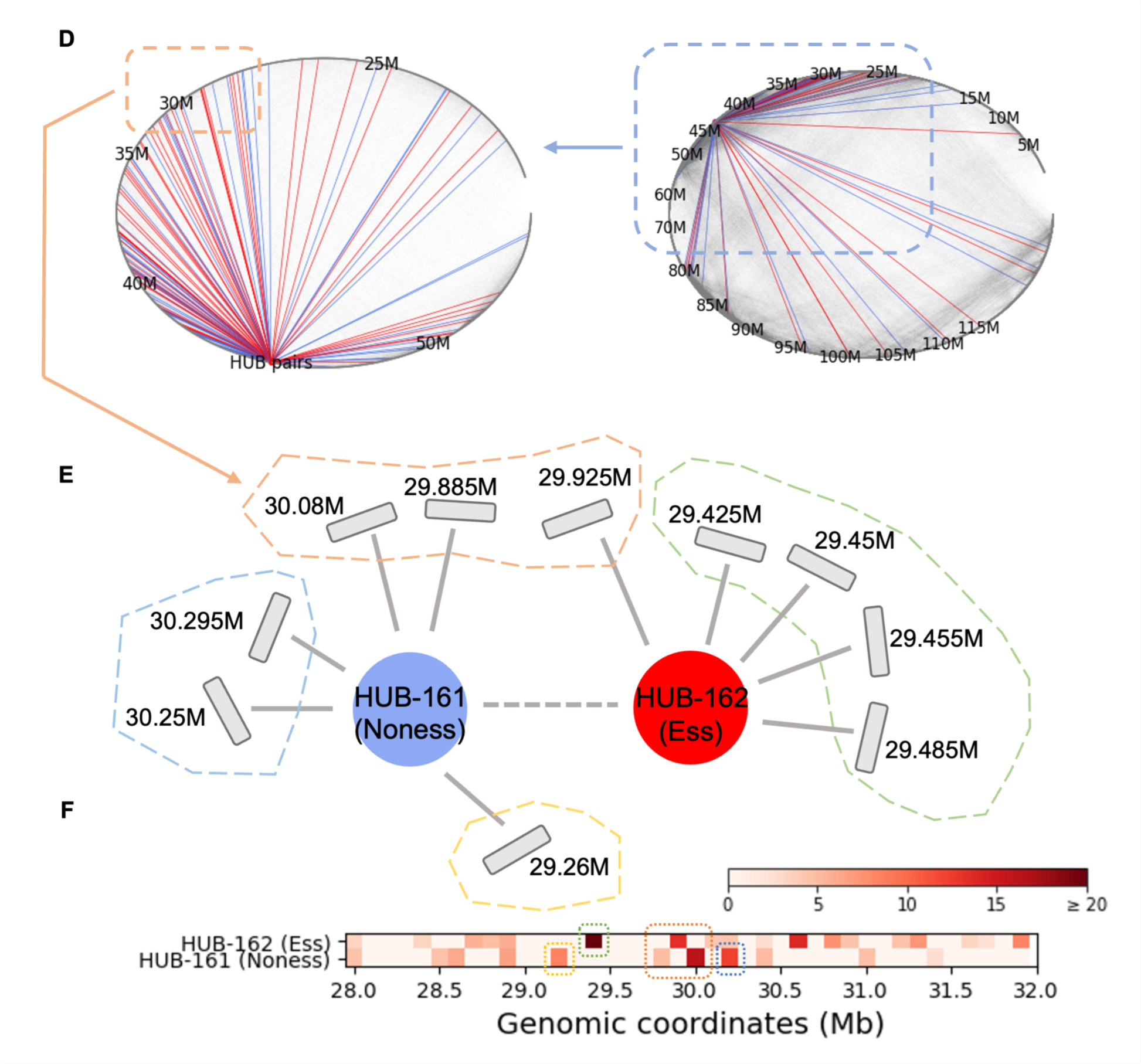
A neighboring pair of essential HUB-162 and nonessential HUB-161. **(A)** Histone marks, TF binding, open chromatin, and total RNA-seq signals surrounding the hub pair (chr11:42,373,450-43,383,450, hg38). The central 10-kb regions within the dashed box correspond to HUB-161 (Nonessential, left, 5-kb) and HUB-162 (Essential, right, 5-kb). More marks are shown in Figure S8. **(B)** Locations of the pgRNAs targeting HUB-161 (Nonessential, left, 5-kb) and HUB-162 (Essential, right, 5-kb). Coordinates are in hg38. **(C)** Survival rate of the hub-KO cells (GFP+). Data are presented as mean ± standard deviation (n = 2). Asterisks represent the *p*-values obtained by performing a two-tailed student’s t-test by comparing the *FC* of hub-KO cells with that of AAVS-1 site-KO cells on day 15. * *p* < 0.05, ** *p* < 0.01, and NS means not significant. **(D)** Cartoon plots show the Hi-C interactions associated with the 5-kb genomic nodes within 3 steps (inclusively) to either HUB-161 or HUB-162 on chr11. The nodes are sorted and aligned counter-clockwise. Red lines represent the edges associated with HUB-162 (Essential), and blue lines represent those of HUB-161 (Nonessential). Coordinates are in hg19. The blue and orange boxes highlight the nodes located in chr11:20M-70M and chr11:28M-32M. **(E)** Cartoon illustration of the 3D contacts between either of the hub pair and chr11:28M-32M. The hubs’ interacting partners are grouped by genomic coordinates in the dashed boxes. Refer to Figure S12 for more examples. **(F)** VC normalized Hi-C read counts in the 100-kb bins paired with HUB-162 (upper, Essential) or HUB-161 (lower, Nonessential).

It is not surprising that the two neighboring hubs have indistinguishable epigenetic features. They are both located in the quiescent state in K562 (Figure 4A and S8), with negligible levels of RNA transcripts measured by total RNA-seq experiments. Neither hub has any ChIP-seq peak of histone modification or transcription factor (TF) binding, nor any functional annotation in the 2.5-kb proximities at both ends. We analyzed their general properties in the FCN, and found that HUB-162 (Essential) interacts with 79 nodes while HUB-161 (Nonessential) interacts with 81. Their interacting nodes largely lack histone modifications (except the heterochromatin mark H3K9me3 indicating condensed chromatin) and TF binding sites, indicating no obvious differences between them (Figure S10). Additionally, the two hubs have similar PageRank scores of the epigenetic marks and TF binding (Figure S11), which is not surprising as the two hubs are adjacent.

However, the two adjacent hubs actually form distinct 3D contact communities. First, the two hubs are not densely connected in the FCN, as shown in Figure 3D, which displays the number of simple paths between all the node pairs in chr11. The two hubs have a scaling factor of 0.37e-5, which is much smaller than those for the other pairs. Second, they only share 2 out of 79 and 81 interacting partners respectively, indicating that they form distinct spatial communities.

We plotted the intrachromosomal Hi-C contacts with either of the two hubs (hg19) (Figure 4D). While their interacting nodes appear clustered around regions such as 42Mb (close to the hub loci), 30Mb and 50Mb regions, the two hubs do not interact with exactly the same nodes. For example, upon zooming in around 30Mb, it is evident that the two hubs form interactions with different loci (Figure 4D ∼ F, and more examples in Figure S12). In addition, there are regions such as 100Mb that specifically interact with only one hub.

Understanding why the deletion of the two hubs caused different impacts on cell viability may provide insight into how the different 3D contacts formed by the two hubs can lead to distinct phenotypic consequences, ultimately helping to reveal the mechanisms of hub essentiality.

### Single cell analyses reveal differential regulatory activities caused by hub deletion

To investigate the impact of hub deletion on cell growth and explore the heterogeneity of the cell population, we performed single-cell RNA-seq and ATAC-seq experiments using the 10x Genomics Chromium platform (see Methods) on the HUB-162 deletion (Essential) and HUB-161 deletion (Nonessential) cells. We jointly clustered the single cells from the RNA-seq and ATAC-seq experiments and then pooled the gene counts and ATAC fragments from each cluster to generate a pseudobulk data (Figure 5A, and S13 ∼ 14), which was then analyzed by a systems biology pipeline called Taiji (K. Zhang et al., 2019). Taiji predicts transcription factor (TF) binding sites using all the known motifs documented in the CIS-BP database (Weirauch et al., 2014) in the open chromatin peaks and then links the TFs to the target genes: if a TF’s binding site is a promoter, the associated gene is considered as the target; if a TF’s binding sites is an enhancer, the TF is linked to the genes predicted as the targets of the enhancer by a non-supervised learning method EpiTensor (Y. Zhu et al., 2016). For each pseudobulk sample, we considered the top 10% of the predicted 3D contacts by EpiTensor that overlap with the ATAC-seq peaks (i.e. active promoters and enhancers), which provided sample-specific TF-target pairs. Taiji then assembled all the TF-target regulations into a genetic network. The global importance of each node (i.e. gene) in the network was assessed by the personalized PageRank algorithm, in which each node was weighted by its expression level and each edge (i.e. the regulatory interaction between a TF and its target gene) weighted by the open chromatin intensity, the binding motif strength and the TF expression level.

**Figure 5.**
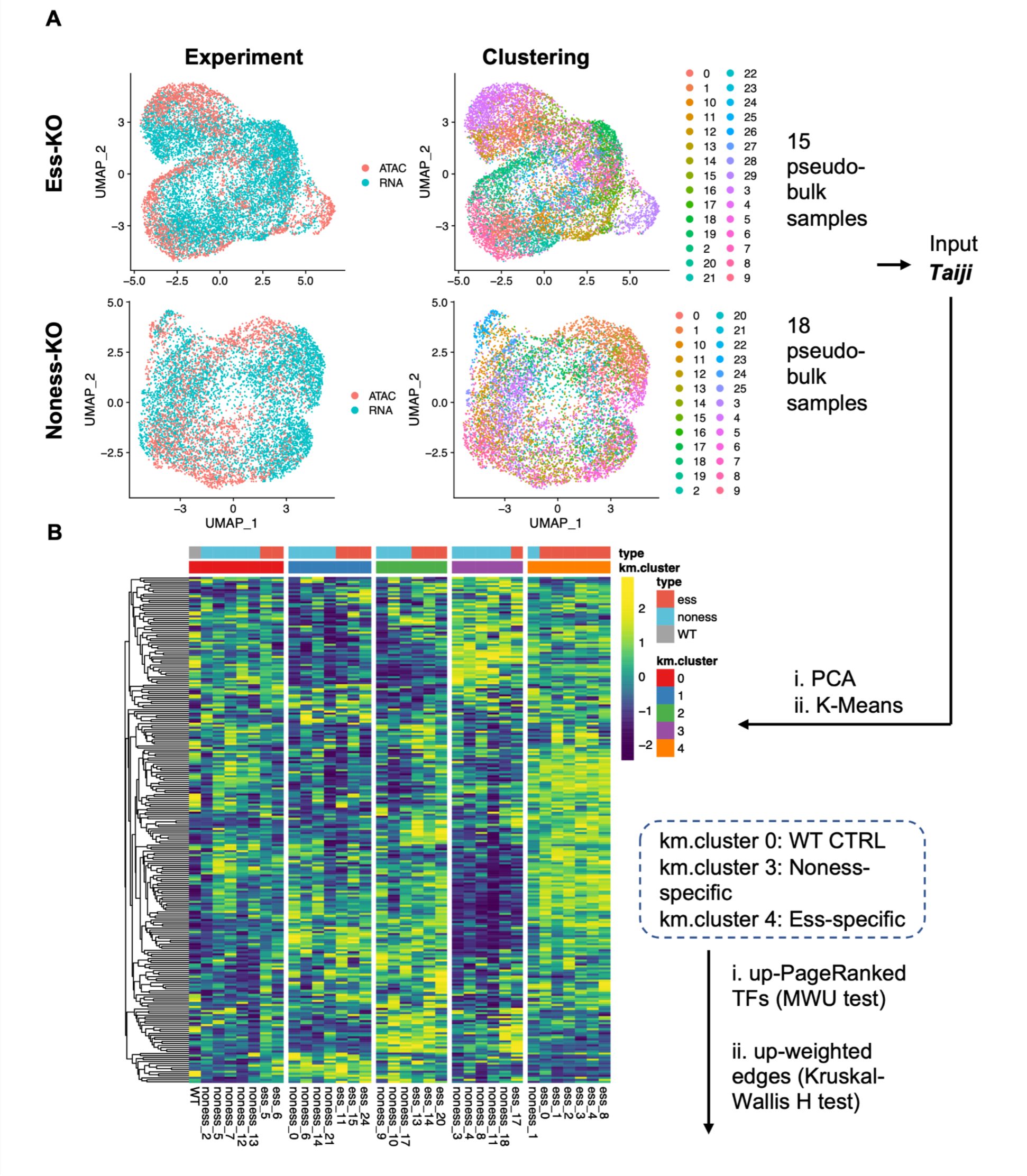
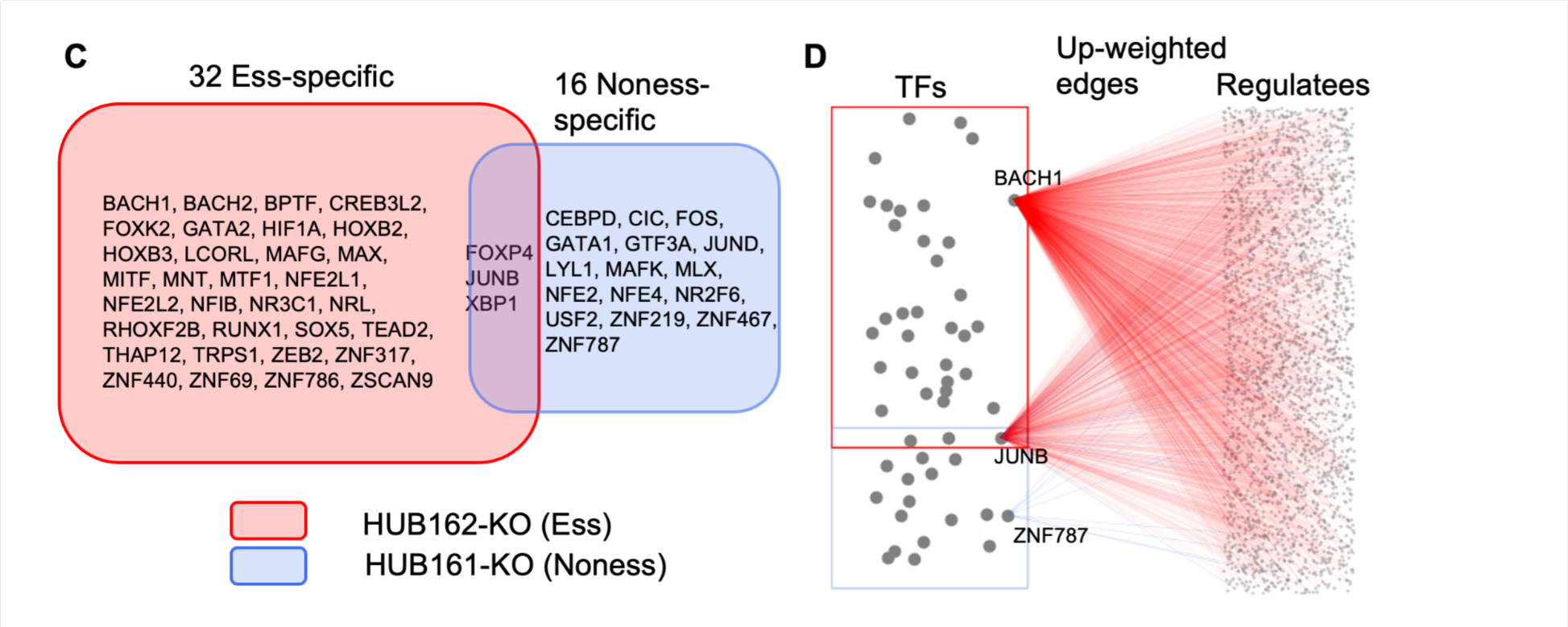
Single cell RNA-seq and ATAC-seq analysis on the hub deletions. **(A)** UMAP visualization plots by co-embedding the single cells in the scRNA-seq and scATAC-seq experiments. 30 clusters were identified for the Essential (red) sample and 26 for the Nonessential (blue). UMAP visualization plots for individual experiments were shown in Figure S13 and S14. Cells in each cluster were pooled into pseudobulks, and further filters were applied. As a result, 15 (Essential) and 18 (Nonessential) pseudobulk samples were used as inputs for the Taiji software (see Methods). **(B)** Heatmap of the Z-scales of the PageRank scores for the TFs, arranged by sample type and K-Means clusters. **(C)** Venn diagram shows the transcription factors (TFs) with significantly increased PageRank scores in the Essential (red) or Nonessential (blue) cluster. The TFs were selected using adjusted *p* <= 0.05 and fold change >= 2 by comparing them with the WT cluster using a Mann-Whitney U test (see Methods). **(D)** Each gray dot in the TF box (red, essential hub deletion; blue, nonessential hub deletion) represents a TF with a higher PageRank score in the hub-deleted cells listed in Figure 5C. Three TFs, BACH1 (essential-specific), JUNB (common) and ZNF787 (nonessential-specific) are representative TFs, and the outgoing edges represent regulatory interactions with their target genes with significantly increased weight (adjusted *p*-values <= 0.05 and percentile rank difference >= 0.5 by comparing with the WT and the cluster of the opposite essentiality type, see Methods). It is evident that essential-specific TFs regulate a significantly larger number of genes than common or nonessential-specific TFs.

We clustered all the pseudobulk data using their TF PageRank score profiles. We first performed PCA on the PageRank matrix and then generated clusters using top four PCs and K-Means with an optimal *K* = 5 selected by the elbow method (Figure S15A ∼ D). Figure 5B shows a heatmap of the clustering results.

We then analyzed the composition of each cluster (Figure S15E, F and TableS8). Cluster 0 contained two HUB-162 deletion (Essential) and five HUB-161 deletion (Nonessential) pseudobulk samples as well as the wildtype (WT), indicating that these cells either did not get sufficient CRISPR deletion or were at the early stages of deletion and still similar to the WT. Cluster 0 contained 24.1% of all the HUB-161 deletion (Nonessential) and 9.1% of all the HUB-162 deletion (Essential) cells, showing a 2.64-fold enrichment of the HUB-161 deletion (Nonessential) cells compared to the HUB-162 deletion (Essential) cells. This observation is consistent with the fact that HUB-161 deletion (Nonessential) is expected to cause less perturbation to the WT and be less lethal. Cluster 1 and 2 were a mixture of HUB-162 deletion (Essential) and HUB-161 deletion (Nonessential) pseudobulk samples without significant enrichment of either one, indicating that these cells were in a common state after hub deletion. Cluster 3 was a HUB-161 deletion (Nonessential)-specific cluster, having 23% of all the HUB-161 deletion (Nonessential) cells from five HUB-161 deletion (Nonessential) pseudobulk samples, but only 3.2% of all the HUB-162 deletion (Essential) cells from one pseudobulk sample, representing a 7.2-fold enrichment of the HUB-161 deletion (Nonessential) cells. Cluster 4 was a HUB-162 deletion (Essential)-specific cluster containing 30.7% of all the HUB-162 deletion (Essential) cells and 6.0% of all the HUB-161 deletion (Nonessential) cells, i.e. 5.12-fold enrichment of the HUB-162 deletion (Essential) cells.

Next, we identified TFs in each cluster with significantly higher (adjusted *p*-value <= 0.05 and *FC* >= 2) or lower (adjusted *p*-value <= 0.05 and *FC* <= 0.5) PageRank scores than the WT cluster (cluster 0), using Mann-Whitney U test (see details in Methods). In cluster 4 (Essential), 35 TFs showed higher PageRank scores, while only 6 showed lower scores than the WT. In contrast, cluster 3 (Nonessential) had 19 TFs with higher and 37 with lower PageRank scores than the WT, respectively (Figure 5C and Table S9).

We then compared the weights of the outgoing edges (reflecting the regulatory strength that considers TF expression, open chromatin strength and TF binding motif affinity) from the TFs with higher PageRank scores in the cluster 3/4 than in cluster 0 (the WT cluster). In cluster 4 (Essential), 14,948 outgoing edges from the 35 important TFs showed considerably higher weights than in cluster 0 (WT) and cluster 3 (Nonessential) (Figure 5D and Table S10, Methods), contributing to the increased PageRank scores of the TFs. In contrast, only 106 outgoing edges from the 19 important TFs in cluster 3 (Nonessential) showed higher weights than in cluster 0 (WT) and cluster 4 (Essential). These observations suggested that the genetic regulatory network became more active in the HUB-162 deletion (Essential) cells than in the WT cells, while the impact was minimal for HUB-161 deletion (Nonessential).

### Deletion of essential hub alters chromatin structure and activates gene regulatory network

We investigated how the deletion of these two adjacent hubs affects the chromatin structure by performing high-resolution Hi-C experiments (Rao et al., 2014). We found that removal of either hub resulted in loss or formation of many spatial contacts. At 5-kb resolution, HUB-162 (Essential) and HUB-161 (Nonessential) deletion led to the loss of 971,917 and 987,107 contacts were lost (ln*p* <= -10 in WT but > -3 after deletion, see Methods), respectively, including (832,685) common ones (Figure 6A). This collapse of the chromatin organization is not surprising as both hubs have many 3D contacts in the WT. As a result, network modularity and effective diameter in the FCN of either hub deletion are increased compared to the WT (Figure S17).

**Figure 6.**
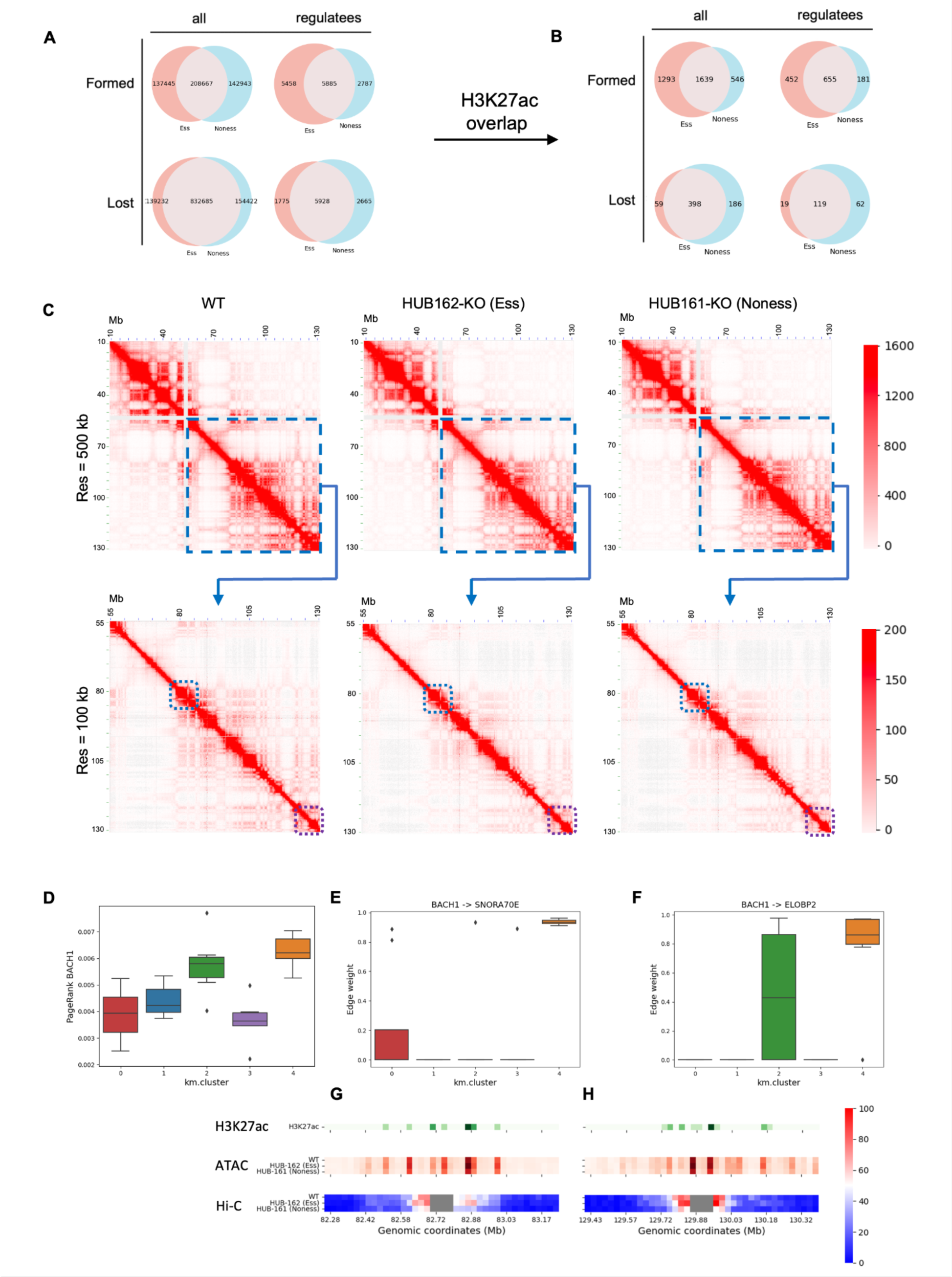
The impact of hub deletion on the chromatin structure. **(A)** The numbers of newly formed and lost Hi-C contacts (5-kb resolution and ln(*p*) <= -10) upon hub deletion compared to the WT. Red: HUB-162 deletion (Essential). Blue: HUB-161 deletion (Nonessential). All, all Hi-C contacts in the genome. Regulatees, Hi-C contacts associated with the weight-altered edges in the Taiji analysis (see Figure 5C and 5D). Results with different ln(*p*) cutoffs are shown in Figure S18A. **(B)** The numbers of formed or lost contacts with both interacting loci overlapping with H3K27ac peaks in WT K562 cells. **(C)** Hi-C contact maps showing VC-normalized reads on chr11, at a resolution of 500 kb and 100 kb, respectively. **(D)** PageRank scores for the transcription factor BACH1 in the K-Means clusters. The importance of BACH1 is significantly increased in cluster 4 (Essential). **(E, F)** The weights for the edge **(E)** BACH1 -> SNORA70E, and **(F)** BACH1 -> ELOBP2 in the K-Means clusters. Each edge has the highest weight in cluster 4 (Essential), and low weights in both cluster 0 (WT) and 3 (Nonessential). **(G, H)** The genome browser tracks of the H3K27ac peaks in K562 WT, ATAC peaks, and Hi-C normalized reads in WT, HUB-162 deletion (Essential), and HUB-161 deletion (Nonessential) samples, in the 1-Mb proximity of **(G)** SNORA70E (blue boxes in the 100-kb Hi-C maps in panel **(C)**), and **(H)** ELOBP2 (purple boxes). The resolution is 25 kb, and the coordinates are in hg19.

HUB-162 (Essential) and HUB-161 (Nonessential) deletion also led to the formation (ln*p* > -3 in WT but <= -10 after deletion, see Methods) of 346,112 and 351,610 contacts, respectively, including 208,667 common ones (Figure 6A). These results emphasized the hubs’ important functions in stabilizing the global chromatin structure: when a hub is deleted, the chromatin organization is altered. Since the two hubs are adjacent, their Hi-C contact maps in Figure 6C largely resemble each other at the global scale but differ in specific loci.

We next examined how these structural changes were associated with the significant TFs and their regulatee genes identified by Taiji. We found that HUB-162 deletion (Essential) induced formation of 5458 new contacts in the promoters of the regulatee genes, much more than the 2787 new contacts induced by HUB-161 deletion (Nonessential) (Figure 6A, and Figure S18). When overlapping these contacts with H3K27ac peaks in the WT cells, we discovered that 271 more (32% higher) contacts were formed upon HUB-162 deletion (Essential) while HUB-161 deletion (Nonessential) led to disruption of 43 more (24% higher) contacts (Figure 6B). This observation is unexpected but consistent with the above analysis that HUB-162 deletion (Essential) caused the gene regulatory network to be more active (i.e. many edges with increased weights) than HUB-161 deletion as the regulatee genes formed more interactions with active enhancers marked by H3K27ac. Figure 6D ∼ H show an example of BACH1, a TF specifically important in the essential cluster 4. It is clear that BACH1 gained new interactions with two of its regulatee genes, SNORA70E or ELOBP2, only upon HUB-162 deletion (Figure 6G and 6H). More examples can be found in Figure S16.

We tested whether the regulatee genes of the important TFs in the essential cluster 4 associated with the new Hi-C contacts are enriched for specific functions or pathways, but no enrichment was found by GREAT (McLean et al., 2010). This observation suggests that the essentiality of HUB-162 was not due to activating specific sets of genes or pathways but rather to a broad alteration to the chromatin structure and enhanced activity of the gene regulatory network.

### Many weak Hi-C contacts formed upon hub deletion lead to increased disorder of chromatin activity

To complement our analysis of Hi-C contact formation or loss based on a *p*-value cutoff, we examined the global trends of all the Hi-C contacts regardless of their significance. We analyzed the strength of the Hi-C contacts in the WT, HUB-162 deletion (Essential) and HUB-161 deletion (Nonessential) cells by plotting the number of contacts against the ln(*p*) of the contacts (Figure 7A ∼ C). As expected from Figure 6A ∼ B, deletion of either hub reduced many 3D contacts, as indicated by fewer contacts in the smaller ln(*p*) area (e.g., ln(*p*) <= -10 in Figure 7A and other cutoffs in Figure S18A). However, deletion of the essential HUB-162 increased the activity of the regulatory network and genome accessibility, especially on the set of the regulatee genes, as shown by more contacts that are associated with H3K27ac peaks and regulatee genes in Figure 7B ∼ C, and S18B.

**Figure 7.**
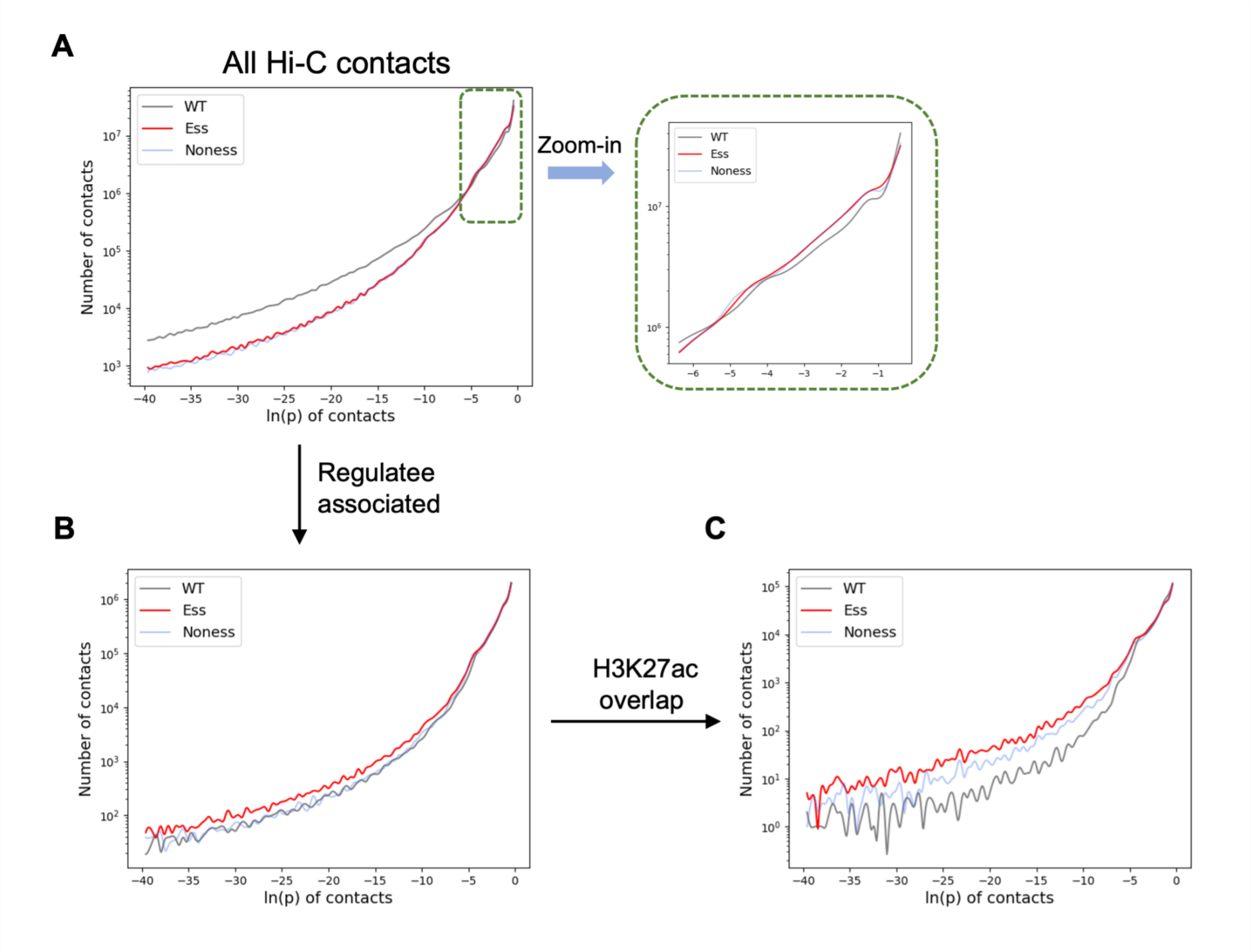
Hi-C contacts statistics. (A) Number of all Hi-C contacts at different ln(*p-*values). The Y-axis is in the logarithmic scale. Contacts with ln(*p-*values) between -6 and 0 are emphasized. (B) Number of the regulatee-associated Hi-C contacts at different ln(*p-*values). (C) Number of the regulatee-associated Hi-C contacts overlapping H3K27ac peaks at different ln(*p-*values).

Noticeably, hub deletion formed many weak contacts (i.e., ln(*p*) between -3 and -10) that did not exist in the WT (ln(*p*)>-3). This effect was more pronounced in HUB-162 deletion (Essential) than in HUB-161 deletion (Nonessential) (Figure S18B). We hypothesized that creating many weak contacts could increase the disorder of chromatin organization and consequently regulatory activity. We tested this hypothesis by measuring the disorder of chromatin activity using scATAC-seq that better characterizes the heterogeneous cell population upon hub deletion. We chose not to use single-cell Hi-C (Nagano et al., 2013) for the analysis because these experiments are difficult to achieve high quality. We calculated Shannon entropy for each pseudobulk sample in the WT (cluster 0), nonessential (cluster 3) and essential (cluster 4) clusters in Figure 5B. We divided the genome into 5-kb bins and computed the percentage of reads falling into each bin, *P_i_*. The Shannon entropy was then computed as *H = −ΣP_i_ln*(*P_i_*) (Figure 8A), which reflects the distribution of the ATAC signals in the genome: a cell state with dispersed open chromatin signals would have higher entropy values than that with spiky and concentrated signals.

**Figure 8.**
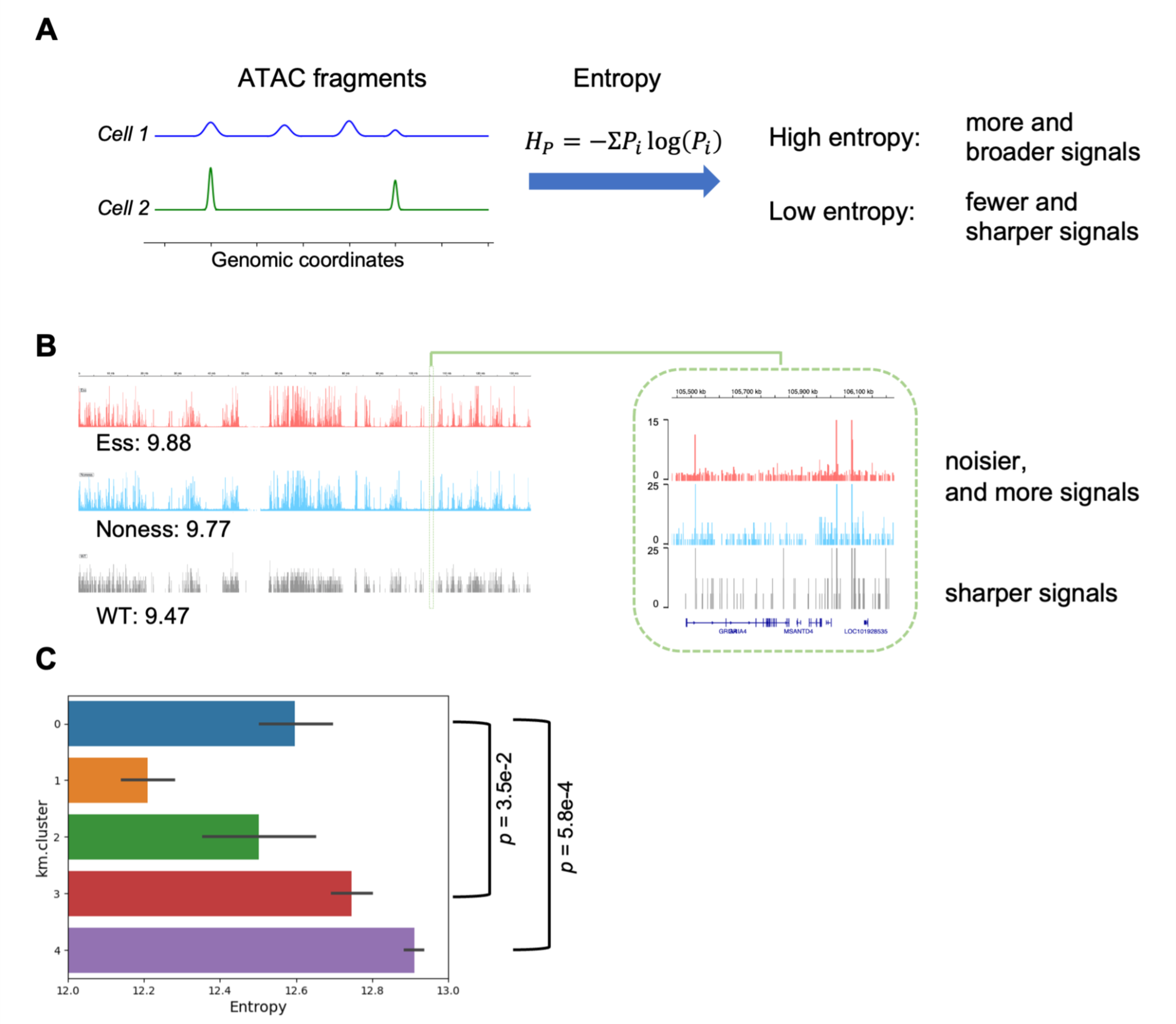
Disorder of chromatin activity. (A) Schematic illustrating the derivation of chromatin accessibility (ATAC) values entropy (refer to Methods). (B) ATAC-seq signal intensities normalized to the same total reads on chr11, for the WT (gray), as well as pseudobulk samples in the Ess-cluster (cls-4, red), and pseudobulk samples in the Noness-cluster (cls-3, blue). Entropy values for the ATAC-seq signals on chr11 are displayed. Region chr11:105.5M-106.5M is shown as an example that deletion of hubs led to noisier ATAC-seq signals. (C) Barplot shows the ATAC entropy values of the five K-Means clusters. *p*-values were calculated by comparing the entropy values of cluster 3 (Nonessential) with those of cluster 0 (WT), or cluster 4 (Essential) with those of cluster 0 (WT), using the Mann-Whitney U test.

We observed that deletion of either hub increased the genome-wide open chromatin entropy compared to WT (cluster 0), with *p*-values of 5.8e-4 and 3.5e-2 for the essential HUB-162 (cluster 4) or the nonessential HUB-161 (cluster 3), respectively (tested with Mann–Whitney U test) (Figure 8C). Moreover, HUB-162 deletion resulted in higher open chromatin entropy than HUB-161 deletion, with a *p*-value of 2.3e-3 tested with Mann–Whitney U test. Unlike the essential one, deletion of the nonessential hub did not always significantly increase entropy on all the chromosomes. (Figure S19). The essential cluster also exhibited universally higher open chromatin entropy values than its nonessential counterpart, with *p*-values ranging from 1.1e-3 to 4.6e-3 (tested with Mann–Whitney U test on different chromosomes respectively). These results suggest that hub deletion increases the disorder of the chromatin activity, and deleting essential hubs causes more significant impact on the order of chromatin organization than the nonessential hubs, thus leading to cell death.

## Conclusions and Discussion

### Neural network model improves the discovery of essential hubs

Our previous research demonstrated that hubs, which are non-coding loci forming many 3D contacts, even in the epigenetically quiescent regions, can be essential for maintaining cell viability and stabilizing chromatin organization. In this study, we report an additional round of CRISPR screening to validate this observation. We developed a neural network model trained on all the hubs in K562 cells, which significantly improved the discovery rate of essential loci by a factor of six, increasing it from 3.5% in the previous screening to 20.6% in the current screening.

These findings suggest that the model captured certain features associated with essentiality. However, we could not identify any distinctive histone modifications, open chromatin, or TF binding profiles specifically associated with the essential or nonessential loci themselves or their 3D contacting loci. One potential explanation is the limited number of loci tested, which consisted of only 1,300 out of 26,148 hubs, with 105 (70 this round + 35 in the previous round) of them being essential hubs. This constrained dataset hindered the training of a more sophisticated model required for accurate prediction of hub essentiality, underscoring the importance of conducting additional screening studies in the future to expand the dataset and improve the predictive capabilities of the model.

### Distinct 3D contacts and phenotypic consequences of neighboring hubs

We further investigated pairs of hubs located close to each other in the linear genome but having opposite essentiality upon deletion. Surprisingly, we discovered that some hub pairs formed distinct 3D contacts despite being adjacent in the 1D genome. This observation is remarkable since neighboring loci are typically expected to have similar 3D contacting partners. The presence of discrete 3D communities and the divergent phenotypic consequences on cell survival indicates that the significance of non-coding loci, particularly those in the epigenetically quiescent state, is determined by their impact on chromatin organization rather than their sequence or epigenetic characteristics.

We performed an in-depth analysis of one such pair, the essential HUB-162 and nonessential HUB-161, using Hi-C data. Our findings revealed that the deletion of either hub led to a significant reduction in spatial contacts compared to the WT. However, the deletion of the essential hub also created a greater number of weak contacts compared to the deletion of the nonessential hub. Furthermore, through single-cell RNA-seq and ATAC-seq analyses, we identified distinct cell states resulting from the deletion of the two hubs, although no specific pathways or biological processes exhibited enrichment. This observation suggests that hub deletions induce a global perturbation in the cell rather than targeting specific genes, which aligns the findings from the Hi-C analysis.

### Enhanced genetic network activity and increased disorder of chromatin activity upon essential hub deletion

We uncovered two major impacts of deleting the essential hub on the cell through the integration of Hi-C and single cell analyses.

Firstly, essential hub deletion induced the enhanced activity of the gene regulatory network compared to nonessential-hub deletion. We observed a significant increase in the formation of new Hi-C contacts, overlapping with H3K27ac, between the uniquely important TFs and their regulatee genes, specifically in the essential hub-KO cell state.

Secondly, essential hub deletion resulted in the creation of many weak contacts, contributing to an increased disorder of chromatin activity. We quantified chromatin disorder using the Shannon entropy of ATAC-seq signals. We observed that essential hub deletion, but not nonessential hub deletion, significantly elevated the open chromatin entropy compared to the WT. It is worth noting that this difference between the essential and non-essential hubs indicates this increase of chromatin disorder is not a result from the introduction of CRISPR into the cell but rather from the functionality of the hubs. This observation uncovers a novel mechanism through which epigenetically quiescent non-coding loci can be important for cellular functions by maintaining the order of chromatin activity and, thus, regulatory functions of the cell.

In summary, our study provides new insights into the role of non-coding loci, particularly those in epigenetically quiescent regions, in regulating cell function and genome organization. We demonstrate that these loci, despite their proximity in the linear genome, can exhibit opposite essentiality for cell viability and form distinct 3D contacts. Moreover, deleting the essential hub elevates genetic network activity and disorder of chromatin activity. Additionally, we provide a powerful computational tool to accelerate the identification of essential loci. Our findings have implications for comprehending the molecular mechanisms of non-coding loci marked by low epigenetic signals, as well as for developing new therapeutic strategies to address diseases involving genomic instability and dysregulation.

## Acknowledgement

We acknowledge the Sanford Consortium (UC San Diego) for providing all FACS and cell sorting services. We acknowledge the staff of the Institute of Genomic Medicine (IGM, UC San Diego) and BioPharma (Admera Health, NJ) for the next-generation sequencing (NGS) services. We acknowledge Dr. Q. Yang and B. Ren (UC San Diego) for instructions on the 10x single-cell chromium experiments. We acknowledge Dr V. Ngo for conceiving the idea of training CNN models to predict hub essentiality. We acknowledge Dr. Z. Chen, Y. Zhao (UC San Diego), and Dr. Y. Liu (Peking University, China) for their guidance and instructions on cell experiments. We acknowledge M. Yao for testing the codes.

## Funding

This project was supported by funds from CIRM (RB5-07012) and the NIH (R01HG009626).

## Competing interests

The authors declare that they have no competing interests.

## Data and code availability

All codes for running the CNN-AE-LR, PR-LR, FCN and entropy analysis are available from the Github repository (https://github.com/yyaoisgood2021/HUB-screening). The sequencing data are accessible through GEO Accession Series GSE231384 and NCBI Sequence Read Archive (SRA) under BioProject ID PRJNA963076.

## Methods and Materials

### Constructing fragment contact network (FCN) from Hi-C

Hi-C data in WT K562 at 5-kb resolution were downloaded from the Rao et al. study (Rao et al., 2014). We constructed the K562-specific genome data by translocating the ABL1 site (chr9:133,589,268-133,763,062, hg19) on chr9 and the BCR site (chr22:23,522,552-23,660,224, hg19) on chr22 to build der9 and phil22. We then aligned the sequencing files onto the translocated K562 genome with Juicer.sh (v1.6) (Durand et al., 2016) with the default settings to generate an inter_30.hic file. We then obtained the expected and the vanilla coverage (VC) normalized read counts for all the interaction pairs at a resolution of 5 kb, and calculated the *p*-values by fitting a Poisson distribution (Lieberman-Aiden et al., 2009).

We constructed a fragment contact network (FCN) (Ding et al., 2021) in which the nodes are 5-kb genomic segments and edges represent Hi-C contacts with log(*p*-values) <= -20 utilizing Stanford Network Analysis Platform (SNAP v6.0) (Leskovec & Sosič, 2016) for each chromosome.

Specifically, any nodes recorded in the blacklist regions (Amemiya et al., 2019) were removed and their related edges were deleted.

### Definition of the hubs

Hubs are defined as the nodes in the FCN with *Z* scores of degree >= 2 in their individual chromosomes. In the K562 cell line, we identified a total of 26,148 hubs.

### Constructing and training of convolutional neural network-autoencoder (CNN-AE)

The architecture of the CNN-AE model is shown in Figure 1A. Training was based on all the hubs defined above (26,148 in the K562 cell line). The CNN-AE model takes the one-hot encoded hub sequences (5kb-long) as inputs, converts it into a latent representation of 40,000 features with an encoder module, and then reconstructs it with a decoder. The model minimizes the categorical cross entropy loss between the input and the reconstructed sequences, using an Adam optimizer with the default parameters (Keras, v2.9.0). Randomly selected 19,611 hubs (75%) were used as training data, while the remaining 6,537 hubs (25%) were held out for validation.

The training was performed with a batch size of 128, an initial learning rate of 0.01, and the default ReduceLROnPlateau and EarlyStopping callbacks. The model was trained for a maximum of 200 epochs. We independently performed 10 runs with random partitions of the input sequences. The learning curve, as shown in Figure S2A, demonstrates that the model reached convergence without overfitting, proving that the architecture and training policies are reasonable. The model with the smallest validation loss was selected for subsequent analyses.

### Constructing and training of logistic regression to facilitate candidate selection

Note that in this part, we did not use the stringent specificity cutoff to remove possible off-target sgRNAs in the Ding et al. study (Ding et al., 2021). As a result, we included 77 essential hubs and 883 nonessential hubs. While some false positives might be included, we argue this approach is reasonable because our primary goal is to select candidate hubs for the second round of screening rather than build an accurate predictor.

The main features we included to train the logistic regression model are the top 30 principal components (PCs) of the sequence representations. Firstly, We applied the CNN-AE model on the hubs tested in the Ding et al. study (Ding et al., 2021) to yield a 40,000-dimensional latent representation vector for each hub. We then performed max pooling (kernel size = 25) to reduce its dimension to 1,600, upon which principal component analysis (PCA) was conducted. Figure 1C and S2C show the top PCs, and essential hubs tend to be located in the area of smaller PC1.

Other features include GC content and node connection degree. Among the 960 hubs screened in the first round by Ding et al. (Ding et al., 2021), we tested whether the essential hubs had higher GC contents or node degrees by Mann-Whitney U test. As a result, essential hubs tend to have higher GC contents (*p* = 3.06e-9, Figure S1A) but the difference in node degrees was not statistically significant (*p* = 0.381, Figure S1B). Since all the hubs in Ding’s screening have high degrees, this finding suggests that the number of Hi-C contacts is not the determining factor for hub essentiality.

Various combinations of these features were tested and the model performances are presented in Figure S2D and E. Logistic regression was trained with the Scikit-learn package (v0.20), with 58 essential and 662 nonessential hubs as training data and 19 essential and 221 nonessential hubs held-out for validation (validation proportion is 0.25). The weight of the cross entropy loss was balanced to 0.92 for essential hubs (1-58/(58+662)=0.92), and 0.08 for nonessential hubs. Each model was trained 20 times with independent dataset splitting and random initialization, and each training session lasted a maximum of 1000 iterations. Figure S2D and E show the ROC-AUC scores and PR-AUC scores. The model with PC1 and 2 achieved the highest ROC-AUC score of 0.73 and was subsequently used in the next section for candidate selection.

### Selecting candidate hubs for CRISPR-Cas9 screening

We applied the following filters to select around 300 5-kb genomic loci in the K562 cell line for the CRISPR screening (Figure 2A). Firstly, the candidate loci must be hubs (nodes with the *Z*-score of degree > 2 in their respective chromosomes). This left 25,188 eligible hub loci untested in the screening conducted by Ding et al. (Ding et al., 2021)

Secondly, we derived the prediction probability for all the hub candidates and removed any loci with probability < 0.4 or with GC content < 0.4. After this step, 4,675 loci remained.

Thirdly, we downloaded peak files for histone modifications (narrow peaks for: H3K4me1, H3K4me2, H3K4me3, H3K9ac, H3K27ac, and H4K20me1; and broad peaks for H3K36me3) and 3 transcription factors (TFs) (CTCF, RAD21, and SMC3). We utilized bedtools (v2.30) (Quinlan & Hall, 2010) to find peaks intersecting with the 5-kb segment of the hub itself or the 2.5-kb flanking regions at each end (i.e. total of 10-kb regions), and retained only the non-overlapped hubs. Broad peaks of H3K9me3 and H3K27me3 were also downloaded but we allowed their intersections. We further confirmed that the remaining loci (10 bk) did not overlap any annotated blacklist regions (Amemiya et al., 2019) and were not listed in the copy number amplification regions (Olshen et al., 2004). The reference datasets can be found in Table S12. After this step, 3,753 loci remained. All the aforementioned steps were performed on hg19, then the remaining candidates were lifted-over to hg38 for sgRNA design using GuideScan (Perez et al., 2017).

Fourthly, we performed gRNA design using GuideScan. Any loci with fewer than 6 pgRNAs composed of eligible sgRNAs were excluded. (see “**Design and construction of pgRNA library**” section for more details). After this step, 543 loci remained. Note that most of the candidates were removed at this step, which is understandable since hub regions were sometimes enriched in heterochromosome regions and were very repetitive, making it difficult to design highly specific gRNAs.

Finally, to accommodate the oligo synthesis size limit, we randomly selected 340 promising hub loci for the screening in this study. (Table S2)

### Design and construction of pgRNA library

For each hub candidate, the sgRNA-design windows are the two 2kb-long flanking regions centered at its two boundaries. We used the GuideScan v1 (the latest version in Apr. 2019 when this design was performed) software (Perez et al., 2017) to search for and score all the possible sgRNAs within the design windows. We then filtered the sgRNAs using the following criteria (Shalem et al., 2014; Wang et al., 2014; S. Zhu et al., 2016, 2017): (1) the sgRNA should have a unique sequence for the intended target locus and at least two base pair mismatches to any other loci in the human genome, (2) its GuideScan specificity score must be above 0.25, (3) the GC content should vary between 0.2 and 0.8, and (4) it should not have “AAAAAA” or “TTTTTT”. Subsequently, the sgRNAs on both sides of the same hub that passed the above filtering criteria were paired together, and all possible pairs were considered. Candidate loci with fewer than 6 possible pairs were discarded, resulting in 6,449 pgRNAs remaining to target 340 candidate hubs. We also included 473 pgRNAs targeting the first exon regions of 29 essential ribosomal genes as positive controls, 100 targeting the *AAVS1* locus, and 100 that do not target the human genome as negative controls (Ding et al., 2021).

The library of pgRNA-harboring lentiviral plasmids was prepared following the methodology described in Ding et al. study (Ding et al., 2021). In short, 128-nt oligos with pgRNA-coding sequences were synthesized with Agilent oligo library synthesis service (Agilent OLS, product No. G7261A) and then cloned into pCG-EGFP backbone using Gibson assembly. The middle sequence was then replaced in a T4 ligation step, forming the complete pgRNA lentiviral plasmid. All steps, including cloning, transformation, PCR, etc., were performed to achieve a minimum of 1000-fold coverage. The primer sequences are listed in Table S1.

### Cell culture

K562 cells stably expressing spCas9 were generously shared with us by Dr. Wensheng Wei (Peking University, China). K562-Cas9 cells were maintained in RPMI-1640 medium (Invitrogen). HEK293T cells were maintained in Dulbecco’s modified Eagle’s medium (DMEM, Invitrogen). Both media were supplemented with 10% fetal bovine serum (FBS, Nucleus Biologics, LLC) and 1% penicillin/streptomycin (from VWR, LLC). All cells were cultured in an incubator with 5% CO_2_ at 37°C.

### CRISPR-Cas9 screening

Lentiviruses that express pgRNA-EGFP were produced in HEK293T cells by transfecting a mixture of pgRNA-EGFP plasmid library, psPAX2 plasmid, and pMD2.G (3:2:1, by mass), and X-tremeGENE™ HP DNA Transfection Reagent (Sigma), following the protocol described by Addgene. To ensure that each cell takes in only one lentivirus particle and that copies of each lentivirus infect at least 200 cells (providing a 200-fold coverage for pgRNA), we infected 30 million K562-Cas9-mCherry cells at an MOI of 0.3. At 72 hours post infection, EGFP and mCherry dual positive cells were sorted at Sanford Consortium (termed as day 0 in the analysis), and were subsequently maintained routinely, and harvested every 5 days until day 25. At least 1.5 million cells were harvested at each time point to ensure a 1000-fold coverage. The collected cells were subjected to (1) genomic DNA extraction by ethanol precipitation, (2) PCR amplification with Caslib-1 primers to specifically amplify the surrounding regions of sgRNA2, (3) PCR amplification with Caslib-2 primers to assemble the library with the Illumina Nextera adapters, (4) construction of Nextera library with a dual index kit (cat. TG-131-2001) and (5) next-generation sequencing (NGS) of single-end reads (SR100) on the Illumina HiSeq 4000 platform. Sequencing was performed at the Institute of Genomic Medicine (IGM), UCSD. Two biological replicates were performed.

### Screening data processing and identification of essential hubs

Since this work is an extension to Ding et al. study (Ding et al. 2021), we hoped to cross-compare our results, thus we did not utilize CRISPR analysis tools such as MAGeCK in this work. We followed the data analysis protocol described in Ding et al. study (Ding et al. 2021) with minor modifications. Firstly, we removed sequencing reads with mutations within ± 4bp of the barcoding sgRNA, and mapped the remaining reads to each sgRNA2. The final successful mapping rates ranged from 65% to 92% with an average of 80.4% (Table S5). Read counts were normalized to hundred reads per million (HRPM), and sgRNAs with HRPM < 0.05 at day 0 were excluded from subsequent analysis, as these gRNAs might not have been amplified to a sufficient copy number in the original library. At this step, 84 gRNAs were discarded.

Next, We performed quantile normalization between the two replicates and calculated the fold change (*FC*) values for each sgRNA by dividing HRPM-day25 by HRPM-day0. Any sgRNA with inconsistent *FC* values in the two replicates was removed: if its *FC* was smaller than the mean *FC* of the positive control set by 0.2 in one replicate, but larger than the mean *FC* of the negative control set by 0.2 in another. At this step, 489 gRNAs (7%) were discarded.

We further ensured that all the sites must have at least 5 remaining pgRNAs for reliable statistical tests. No sites were discarded at this step. After applying these stringent filtering criteria, we obtained 5,876 pgRNAs targeting 340 hubs for further analyses.

Next, we computed the mean of *FC* values between the two replicates as the *FC* for each pgRNA (*FC_guide_*) (Table S3). For hub candidates and ribosomal genes, we calculated the *FC* scores (*FC_site_*) by averaging (*FC_guide_*) over all the pgRNAs targeting that site. Since (*FC_guide_*) values rely not only on intrinsic properties but also on the composition of the entire pool of the sequencing library, we could not directly compare these values with those derived from the first round of screening in Ding et al. study (Ding et al., 2021). We thereby log-linearly normalized all the *FC* values using the following formula:

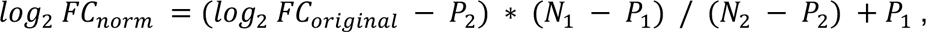

where *P*_1_ and *N*_1_ are the mean *log*_2_ *FC* for all the gRNAs belonging to the positive and negative controls in the first round screening, and *P*_2_ and *N*_2_ are the corresponding values in the second round.

This transformation ensured that the *FC* values of the control gRNAs were comparable and formed the same distribution (*p* = 0.816 for positive controls, and *p* = 0.979 for negative controls according to Mann-Whitney U test), making them directly comparable (Figure S4).

To determine the score cutoff for calling essential hubs, we performed a two-sided Mann-Whitney U test, by comparing the (*FC_guide_*) scores of the candidate sites against those of all the pgRNAs in the negative controls (AAVS-1-targeting and non-targeting pgRNAs), and thus derived a *p*-value for each site. We plotted log_2_(*FC_site_*) vs. -log_10_(*p*-value), as a V-plot (Figure 3B). We also created 100 virtual negative sites by sampling 20 pgRNAs from the negative controls with replacement per virtual site to indicate their locations on the plot. We observed a clear gap between the positive and negative control sites, which aided in determining the decision boundary for the essential hubs. We computed the *Z* scales for log_2_(*FC_site_*) and -log_10_(*p*-value), and the final essentiality scores were derived as follows:

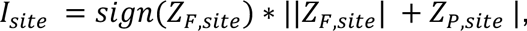

where *Z_F,site_* is the *Z* scale of log_2_(FC) of a given site, and *Z_P,site_* is the *Z* scale of its - log_10_(*p*-value)

To make the two rounds of screening comparable, we selected the same *I* score cutoff of -1.0 as the in Ding et al. study (Ding et al., 2021) to call essential hubs in the second round of screening. As a result, 70 hubs were identified as essential, achieving a hit rate of 20.6% (70/340) among the total loci. This hit rate is significantly higher than the 3.6% hit rate observed in the first round screening (Table S4). These findings suggested that our selection criteria of candidate loci captured informative features of the essential hubs.

### Individual validation for hub essentiality by proliferation assay

The primers required for cloning the hub-targeting pgRNA sequences into the pCG lentiviral-expression plasmids are listed in Table S1. We cloned the pgRNAs into individual pCG lentiviral expression plasmids through a Golden Gate ligation, and packaged the individual pgRNA-EGFP expressing lentivirus according to the Addgene manual (https://www.addgene.org/guides/lentivirus/#resources, accessed in June, 2019). K562-Cas9-mCherry cells were then infected with controlled MOI of <= 1. The impact of hub knockout (hub-KO) on cell proliferation was measured by the reduction in the percentage of GFP/mCherry (Cas9) dual-positive cell populations. The initial GFP/mCherry% was assessed 72 hours post infection, and then monitored every 6 days using fluorescence-activated cell sorting (FACS) until day 15. We conducted at least two biological replicates and performed a two-tailed t-test to compare the *FC* values of all hub-targeting pgRNAs against the AAVS-1-targeting pgRNA, assessing whether hub-KO led to a significant decrease. Detailed information can be found in Table S6 and Figure 2D and S5.

## Classifying hub essentiality with sequence and epigenetic features

We constructed the FCNs as described in the **“Constructing fragment contact network (FCN) from Hi-C”** section. Multiple features were incorporated as node metadata, and a PageRank algorithm was employed to derive the centrality score for each feature. Subsequently, logistic regression models were utilized to classify the hub essentiality. The detailed steps are outlined below.

### Preparation of features for ChIP-seq data and gene profiling

We downloaded the following datasets from ENCODE: (a) ChIP-seq peaks of core histone modifications: H3K4me1, H3K4me3, H3K9me3 (broad), H3K27ac, H3K27me3 (broad), H3K36me3 (broad); (b) additional histone modifications: H3K4me2, H3K9ac, H4K20me1; and (c) peaks of CTCF, RAD21, and SMC3. Accession codes are listed in Table S12. Gene annotations were downloaded from Gencode (Harrow et al., 2012). Information regarding gene (Morgens et al., 2016; Wang et al., 2015) or long-noncoding RNA (lncRNA) (Liu et al., 2018) essentiality for cell survival in K562 cell lines was obtained from CRISPR-KO assays. We used bedtools (Quinlan & Hall, 2010) to intersect all the 5-kb nodes with ChIP-seq peaks and gene coordinates, generating a binary feature map for each node, where a value of 1 indicated the presence of a ChIP-seq peak or gene, while 0 indicated no overlap. This feature map was then employed as the personalization dictionary in the PageRank algorithm.

### PageRank for each feature

Personalized PageRank was executed with the Networkx package (v 2.5.1) ((Hagberg, Aric A., Schult, Daniel A. and Swart, Pieter J., n.d.)), with a default damping factor of 0.85 and a convergence error tolerance of 1e-12. The edge weights were set to 1. PageRank scores for all nodes corresponding to each ChIP-seq signal were calculated with five repetitions and then averaged. Additionally, a default PageRank score with no personalization was derived, considering every node to have equal importance, also with five repetitions. Subsequently, we normalized the PageRank scores across different chromosomes by computing their *Z*-scales as centralities within their respective chromosomes.

### Logistic regression models

We combined the following factors: (a) Top principal components (PCs) of the sequence representations derived from the CNN-AE model, (b) centralities of the ChIP-seq and gene profiling features, and (c) GC contents, as the metadata of each hub. To mitigate any experimental errors and GC bias present in the Ding et al study (Ding et al., 2021), we excluded (1) the low-specificity pgRNAs with specificity harmonic scores <= 0.1, and (2) hubs of GC% < 0.4. After applying these filters, there are 91 essential and 548 nonessential hubs left from both rounds of screening.

The logistic regression part was implemented with the scikit-learn package. We split the dataset into a training and a test set in a ratio of 0.75:0.25. In both datasets, we maintained the same ratio (91/(91+548)=0.142) for essential hubs. Class weights were adjusted (Ess: 548/(91+548), and Noness: 91/(91+548) to address the issue of data imbalance. Default data preprocessing was performed using the scikit-learn package. Twenty independent repeats were performed. Finally, the evaluation metric was derived as the area under the ROC curve (ROC-AUC) for the essential hubs in the test set. Figure S6D showed the balanced ROC-AUC performance.

### Motif search

Motifs were searched using the findmotifs.pl (Homer, v4.11) with the default settings. The foreground consisted of the 91 essential hubs (refer to the “**Logistic regression models**” section above), while the background included the 548 nonessential hubs.

### Simple path analysis on the hub pairs

For all the chromosomes (chr1-8, 10-21, X, der9, and phil22), we extracted all possible pairs of hubs located within a 50-kb genomic distance. Among these pairs, there were 4 Essential-Essential pairs, 135 Essential-Nonessential pairs, and 396 Nonessential-Nonessential pairs (see Table S7).

To assess the denseness of each pair, we applied the concept of simple paths (paths linking two nodes without repeating intermediate nodes). The number of simple paths from node **i** to node **j** at a step of *n* (> 1) can be derived using the following equation. 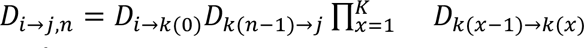, where *K* represents the set of intermediate nodes.

According to the equation, the number of simple paths is dependent on the degrees of the pair. So we then normalized *D* using the following equation:

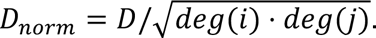

To search for simple paths, we used the all_simple_paths function from the Networkx package, with a step cutoff ranging from 1 to 4. The normalized number of simple paths with step n (*D_n_*) scales exponentially with the step *n*. therefore, we ran a curve fit with the equation:

*D = Ae^Bn^*+ *C* where *A*, *B*, and *C* are coefficients to optimize. The exponential factor *B* depends on the degrees of the intermediate nodes and these values are various on different chromosomes, so we constrained *B* to have the same value for each individual chromosome, and this helped to avoid overfitting. The scaling factor *A* was further used to evaluate the density of the connections between hub pairs. The fitting process was implemented using scipy.optimize.curve_fit. (Table S7)

### Centrality contribution analysis

The centrality contribution from the interacting node *i* was computed using the equation: *C_i_ = PR_i_ / deg_i_ * a*, where *PR_i_ PR_i_* is the PageRank score of the node *i, deg_i_* is the degree of the node *i*, and *α* is the damping factor, which is set to 0.85.

### Selection of a pair of indistinguishable hubs for comparative analysis

We applied the following criteria to select hub pairs: (1) one hub must be essential (*I* score <= -1) and the other nonessential, with a minimum difference in their *I* scores of 2, (2) they should be located within 50 kb in the linear genome and no more than 3 nodes away from each other in the FCN, (3) both sites should be located in the quiescent regions without any overlap with gene exons, annotated regulatory elements, and ChIP-seq peaks of histone modifications, (4) both loci should be well mappable, (5) both loci should be at least 1 Mb away from the centromere, and (6) both sites should be indistinguishable in the PageRank logistic regression models, with the nonessential hub showed no significantly lower predicted values than its essential counterpart.

After applying these filters, we selected HUB-162 (chr11:42,878,450-42,883,450, hg38), an essential hub with a negative *I* score of -1.01, and its immediate genomic neighbor, HUB-161 (chr11:42,873,450-42,878,450, hg38), a nonessential one with a positive *I* score of 2.16, showing minimal effects on cell growth if deleted.

### Individual validation on the candidate pairs

Individual proliferation assays were performed according to the same protocol described above. The primer sequences can be found in Table S1, and detailed data can be found in Table S6.

### Measurement of hub deletion ratio

We chose Ess-pg2 and Noness-pg4 for all downstream analyses. To rule out the possibility of deletion efficiency difference, we performed qPCR. K562-Cas9 cells infected with either HUB-162 (Essential) or HUB-161 (Nonessential) lentivirus at an MOI of 1 were sorted on day 3 post infection. Cells were harvested on day 8, 10, and 16 post infection, and the genomic DNAs (gDNAs) were extracted with the Quick-DNA kit (Zymo, D4068). We also isolated the gDNAs from K562 WT cells as the control. qPCR was performed in a Biorad CFX Opus 96 machine under the default conditions. Two qPCR replicates were performed. The primer sequences can be found in Table S1.

### scRNA-seq experiments

K562-Cas9 cells were infected with either HUB-162 (Essential) or HUB-161 (Nonessential) lentivirus at an MOI of 1 and sorted on day 10 post infection. Libraries were prepared according to the 10x Genomics chromium GEM Single Cell 3ʹ RNA library manual (product numbers PN-1000075 and PN-1000153, and document number CG000183-Rev C). Approximately 10,000 hub-KO cells (with > 90% live rate at the experiment date) were used to prepare each library. Each library was sequenced to obtain 250M of pair-end 150nt (PE150) reads on the HiSeq 4000 platform, by Biopharma Services at Admera Health, NJ.

### scRNA-seq data processing and cell clustering

The sequencing data from both the HUB-162 deletion (Essential) and HUB-161 deletion (Nonessential) libraries was counted and aggregated with the 10x Genomics Cellranger (v7.0.0), and loaded with Seurat (v4.1.0) with the parameters min.cells = 200 and min.features = 200. Low-quality single cells were filtered out if they did not satisfy the criteria: (percent.mt < 5) & (nCount_RNA > 5000) & (nCount_RNA < 50000) (Figure S13A, B) (Hao et al., 2021). The remaining cells (9321 Essential and 5899 Nonessential) were visualized with a UMAP, showing minimal batch effects between the Essential and Nonessential libraries (Figure S13C). We then processed the Essential and Nonessential libraries separately. We extracted the expression of all non-mitochondrial genes, performed log-normalization and scaling, and reduced the dimensions by PCA (npcs = 30, 23.2% explained variance for the Essential library, and 28.4% for the Nonessential library). FindNeighbors and FindClusters were then run on the top 30 principal components (PCs) with a resolution of 3. Consequently, 30 clusters were identified for the Essential library, and 26 for the Nonessential library (Figure 5A).

### scATAC-seq experiments

K562-Cas9 cells were infected with either HUB-162 (Essential) or HUB-161 (Nonessential) lentivirus at an MOI of 1 and sorted on day 10 post infection. Libraries were prepared according to the 10x Genomics chromium Next GEM single cell ATAC manual (product numbers PN-1000175 and PN-1000162, and document number CG000209-Rev F). Approximately 10,000 hub-KO cells (with > 90% live rate at the experiment date) were used to prepare each library. The libraries were sequenced to obtain over 100M of PE100 reads on the Illumina Novaseq platform, at IGM, UCSD.

### scATAC-seq data processing and cell cluster prediction

We randomly downsampled the sequencing reads of the HUB-162 (Essential) library by a factor of 0.4 to ensure similar depth with that of HUB-161 (Nonessential). The sequencing data was counted by the Cellranger software (v7.0.0), and processed with Seurat (v4.1.0) and Signac (v1.8.0) (Hao et al., 2021; Stuart et al., 2021). The count data was loaded with parameters min.cells = 20 and min.features = 1000. We then filtered out low-quality single cells if they did not satisfy the criteria (peak_region_fragments > 3000) & (peak_region_fragments < 20000) & (pct_reads_in_peaks > 15) & (nucleosome_signal < 4) & (TSS.enrichment > 2), resulting in 3554 cells in the Essential library and 2559 in the Nonessential library (Figure S14). Gene activities were calculated using the top 2000 variable ATAC peaks, and finally anchor transfer was performed with the default parameters to assign each single cell in the ATAC experiment to a cluster in the scRNA experiment (Figure 5A).

### Data preparation for Taiji input

For the K562 WT sample, the raw sequencing Fastq files of the bulk mRNA or poly-A plus RNA-seq were downloaded from ENCODE (Table S12), and aligned with STAR (2.7.10a) (Dobin et al., 2013), and mapped to annotations (v41) (Harrow et al., 2012) to generate GeneQuant files without any normalizations. The GeneQuants files were then summed up as the input for Taiji. The IDR thresholded peak files of the bulk ATAC-seq were downloaded from ENCODE (ENCSR483RKN, Table S12) and used directly in the ATAC-seq “NarrowPeak” format. The top 10% of the promoter-enhancer loops identified from Epitensor (Y. Zhu et al., 2016) were used under the Hi-C “ChromosomeLoop” tag.

For the hub-KO pseudobulk samples (Essential or Nonessential), clusters containing fewer than 100 scRNA-seq cells or < 20 scATAC-seq cells were considered too sparse and were thus removed. Raw gene counts for the cells in each cluster were extracted and summed up to create the GeneQuant files, and the counts of the corresponding ATAC fragments were extracted and summed up and normalized to the same total read counts as the alignment files under the ATAC-seq “PairEnd” tag. Same as the WT control sample, the top 10% of the promoter-enhancer loops identified from Epitensor (Y. Zhu et al., 2016) were directly used under the Hi-C “ChromosomeLoop” tag.

### Identifying important TFs using Taiji

The Taiji program was run with the default settings to yield the following results: (1) ATAC-seq peaks were called for each hub-KO pseudobulk sample using MACS2 (Y. Zhang et al., 2008), (2) for each sample, a regulatory network was constructed, and (3) PageRank scores of the transcription factors (TFs) were generated. We calculated the *Z*-scales of the PageRank scores to identify the top 250 variable TFs across all the pseudobulk samples, including bulk WT, and then PCA was performed (Figure S15A, B). We then ran the within-cluster sum of square (elbow) method and Silhouette score method with the “factoextra” (Kassambara, 2016) and “NbClust” (Charrad et al., 2014) packages on the top four PCs to determine the optimal *K* value. The optimal *K*, derived from the elbow method and Silhouette method using Manhattan distance, was 5 (Figure S15C, D).

For both Essential and Nonessential libraries, we calculated their proportions of the single cells (in the scRNA-seq or scATAC-seq experiments) in each K-Means cluster (km.cls), and the ratio of Essential proportion to Nonessential proportion reflected the enrichment of the Essential library (Figure S15E, F, and Table S8). We notated cls-4 as Essential-specific, 3 as Nonessential-specific, 1 and 2 as mixed-type, and 0 as WT.

To identify TFs with significantly increased or decreased scores (importance) in each K-Means cluster (km.cls), we performed Mann-Whitney U test on the normalized PageRank scores against the WT cluster (cls-0). The *p-*values were adjusted with the Benjamini & Hochberg method. We then selected TFs with the (1) increased regulatory importance by applying the filters of adjusted *p-*value < 0.05 and FC > 2, or (2) decreased regulatory importance by applying the filters of adjusted *p-*value < 0.05 and FC < 0.5. The full list is in Table S9.

### Identifying cluster-specific regulatory edges (regulatees)

We extracted all the edges with significant TFs identified in the previous step as start nodes and determined their weight percentiles. Edges not appearing in certain pseudobulk samples may suggest either these edges are too weak to be detected or these edges are noisy and not reliable. Thus, we removed noisy edges that did not consistently exist in >= 5 pseudobulk samples in any K-Means cluster. For the remaining edges, if they were not detected in certain pseudobulk samples, it indicated their weakness, and their percentiles were assigned as 0.

Next, we analyzed the percentiles of the edges with a Kruskal-Wallis H-test, and selected the significantly changed edges with *q* values <= 0.05 (by adjusting the *p*-values with Benjamini & Hochberg method) and percentile differences (compared with the WT cluster and the cluster of the opposite essentiality type) >= 0.5 (increased weights) or <= -0.5 (reduced weights). The numbers of significantly changed edges were shown in Table S10.

The regulatees are defined as the end nodes of the corresponding edges.

### Hi-C experiments

K562-Cas9 cells were infected with either HUB-162 (Essential) or HUB-161 (Nonessential) lentivirus at an MOI of 1 and sorted on day 10 post infection. Hi-C samples were generated using the Arima Hi-C kit for mammalian cell lines, following the manufacturer’s manual (A160134 v01). For each sample, 1 million infected cells with a live rate of > 90% at the time of the experiment were used for Hi-C library preparation. Each library was sequenced for 1.5 billion of pair-end 100nt (PE100) reads on the Illumina Novaseq platform at IGM, UCSD.

### Hi-C data processing

The K562 genome was constructed as described in the **“Constructing fragment contact network (FCN) from Hi-C”** section. We processed the raw fastq files using the Juicer pipeline (v1.6) with the default settings (Durand et al., 2016) to generate inter_30.hic files. We then obtained the expected and vanilla coverage (VC) normalized read counts for all interaction pairs at a resolution of 5 kb or 25 kb, and calculated the *p-*values by fitting a Poisson distribution (Lieberman-Aiden et al., 2009).

### Effective diameter analysis

FCNs of the hub-KO samples were constructed for each chromosome according to the protocols described in the **“Constructing fragment contact network (FCN) from Hi-C”** section. Effective diameters were calculated using GetAnfEffDiam (SNAP v6.0) (Leskovec & Sosič, 2016). The algorithm was run ten times, and the average was taken to account for differences caused by random initialization.

### Modularity analysis

Communities were detected for each chromosome using Clauset Newman Moore method, and the corresponding modularity was computed with CommunityCNM (SNAP v6.0) (Leskovec & Sosič, 2016). Modularities of each chromosome in seven WT cell lines (GM12878, K562, HUVEC, IMR90, NHEK, KBM7, and HMEC) were tested as normal distributions in the Ding et al study, so we tested how deviated the modularities would be upon hub-KO using the cumulative distribution function (*cdf*) of the normal distribution. *p*-values were derived as 1 - *cdf*.

### Identifying TF/regulatee-associated Hi-C contact pairs

We extracted the transcription start site (TSS) coordinates of the TF/Regulatees from gencode (Harrow et al., 2012) and extended them to integer 5-kb bins (by up to 5 kb). These coordinates were then intersected with the 5-kb Hi-C maps.

### Identifying changes in Hi-C contact pairs

For the 5-kb or 25-kb contact maps (refer to **Hi-C data processing section** for the procedures to calculate the *p*-values) generated from hub-KO bulk Hi-C experiments or WT bulk Hi-C experiment (Rao et al., 2014), unreliable interaction pairs were removed if they were not detected in at least two experiments. We collected the remaining interaction pairs and compared their *p*-values. Missing values suggest those interaction pairs were too weak to be detected and were thus assigned *p*-values of 1.

We then defined interactions with *p*-values <=*e*^-10^, or *e*^-15^, or, *e*^-20^ as significantly strong, and *p*-values >= *e*^-3^ as negligibly weak. Formed interactions referred to the pairs with *p*-values >= *e^-3^* in the WT sample but below the significant thresholds in the hub-KO sample, while lost interactions vice versa (Table S11).

### Identifying H3K27ac-associated Hi-C contact pairs

The H3K27ac narrow peak file (ENCFF044JNJ) for the WT K562 cells was downloaded from ENCODE. For each contacting pair in Hi-C, we considered it to be associated with H3K27ac if both sites overlapped with H3K27ac peaks.

### Entropy analysis for chromatin accessibility

We extracted and pooled the scATAC-seq fragments within each pseudobulk sample (generated using the procedures described in the **scATAC-seq data processing and cell cluster prediction** section). To ensure consistent coverage across different sizes of K-Means clusters, we randomly sampled fragments from the pool. The fragment counts were then normalized as reads per million (RPM). We converted the RPM data into a density plot within each 5-kb genomic bin for each chromosome (or for the whole genome) and calculated the entropy using scipy.stats.entropy. We collected the entropies of all the pseudobulk samples in the same K-Means cluster (refer to **Identifying important TFs using Taiji** section), and performed Mann-Whitney U tests to compare the entropy of km.cls-4 (Essential) or cls-3 (Nonessential) with that of cls-0 (WT).

## Notes

https://github.com/yyaoisgood2021/HUB-screening

### Competing Interest Statement

The authors have declared no competing interest.

