## supplementary figures and tables for "Distinct 3D contacts and phenotypic consequences of adjacent non-coding loci in the epigenetically quiescent regions"

**Supplementary materials**

**This PDF file includes:**

**Figure S1 ~ S18**
**Table S5 Mapping rate of the CRISPR screening**
**Table S10 Number of significantly changed edges in Taiji analysis**

**Other supplementary materials for this manuscript includes excel files for:**

**Table S1 Primer sequence information**
**Table S2 Hub information**
**Table S3 gRNA fold change (FC) values**
**Table S4 Hub HTS results**
**Table S6 Individual validation for hub essentiality**
**Table S7 Hub pair analysis in the fragment contact network (FCN)**
**Table S8 Cluster information about the single cell experiments**
**Table S9 TFs with significant fold change (FC) in the Taiji analysis**
**Table S11 Hi-C contact formation or Loss**
**Table S12 Reference datasets from ENCODE**
**Table S13 Quality reports of the NGS data**

### Supplementary Figures and legends

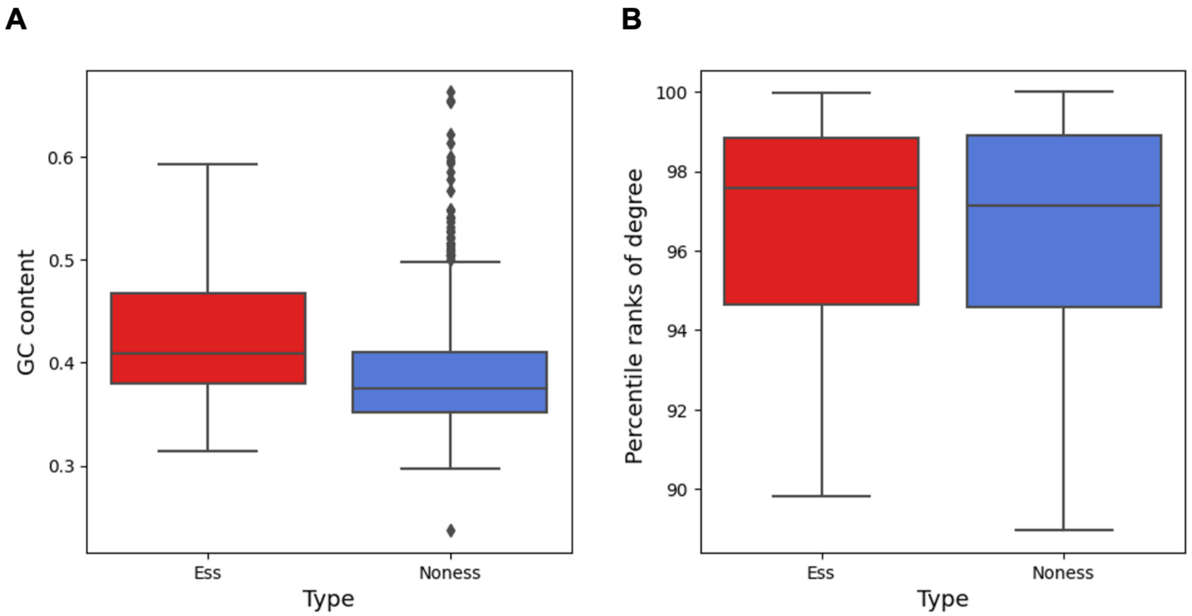


**Figure S1. Hub essentiality vs. GC content and node degree**

**(A)** Boxplot of the GC contents of essential (red) and nonessential (blue) hubs identified from the first round of screening in (Ding et al. 2021). Essential hubs tend to have higher GC contents ($p$ = 3.06e-9 with Mann-Whitney U test).

**(B)** Percentile rankings of the hub degree. Essential hubs do not have significantly different percentile rankings of node degree ($p$ = 0.381 with Mann-Whitney U test).

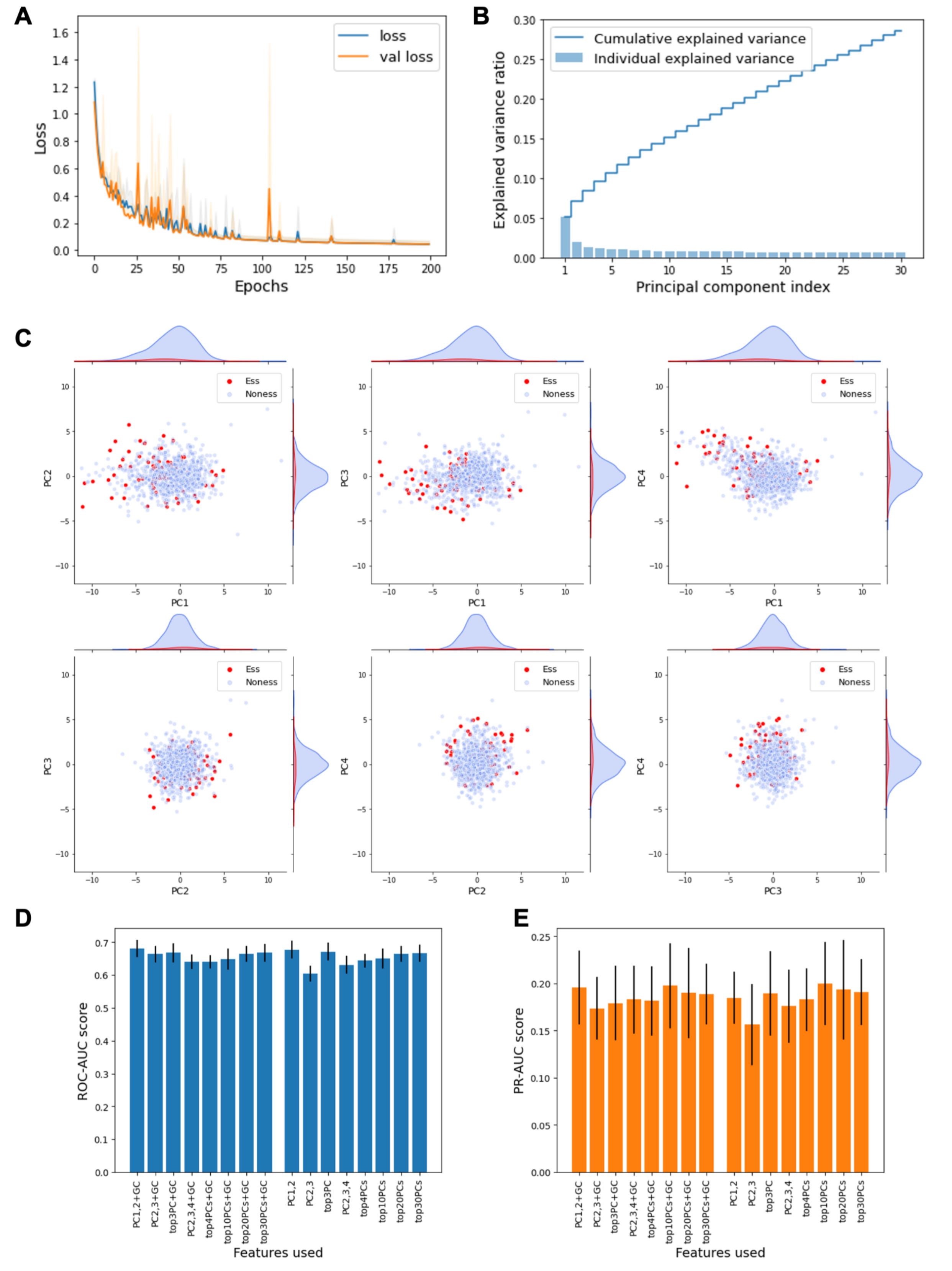


**Figure S2. CNN-AE LR model**

**(A)** Loss curve of the CNN-AE model. We performed 10 independent runs to generate this loss curve. Blue curve: averaged training loss vs epoch. Orange curve: average validation loss. Gray area: the 95% confidence intervals of the training loss over the 10 replicates. Orange area: the 95% confidence intervals of the validation loss, and it almost overlaps with the gray area. There is no significant overfitting and the model achieved convergence at around epoch 200.
**(B)** Variance explained by the top 30 principal components.
**(C)** PCA plots of the latent representations of the sequences of the hubs tested in Ding et al. study.
**(D)** ROC-AUC scores on the validation datasets for the models using different combinations of sequence features (PCs + GC content). The data is represented as mean ± standard deviation in 20 independent replicates. Using the top2 PCs without GC content can achieve the optimal performance.
**(E)** PR-AUC scores. Random guess is 0.08.

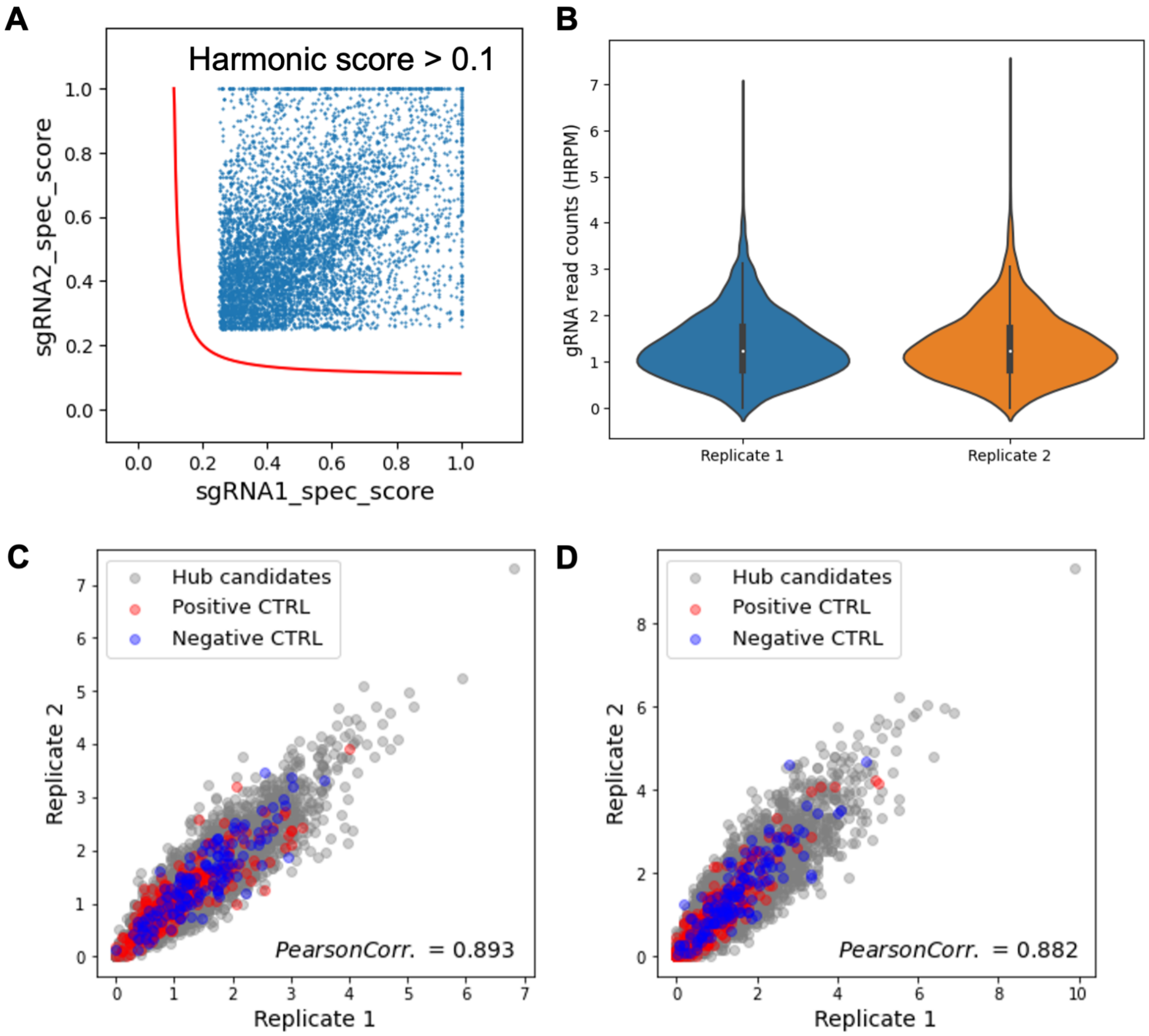


**Figure S3 Quality control for the CRISPR screening**

**(A)** Scatterplot shows the specificity scores of the gRNAs in the screening pool. The red line represents the boundary line of specificity harmonic score of 0.1. pgRNAs beyond the line suggest reliable cutting specificities (Ding et al. 2021; Perez et al. 2017; Tycko et al. 2019).

**(B)** Violinplots show the normalized barcoding gRNA read counts (HRPM) on day 0 in the two biological replicates. Day 0 data indicates the library quality. It is evident that the two replicates have the same average (1.342), and the counts are concentrated within small ranges (> 90% of data points are located in 0.2 ~ 2.3). This suggests that the experiment could be reproducible and the screening library had a uniform representation of each pgRNA in the pool.

**(C, D)** Scatterplots show correlations of the normalized barcoding gRNA read counts (HRPM) between the two biological replicates in the current screening. **(C)** on day 0. **(D)** on day 25.

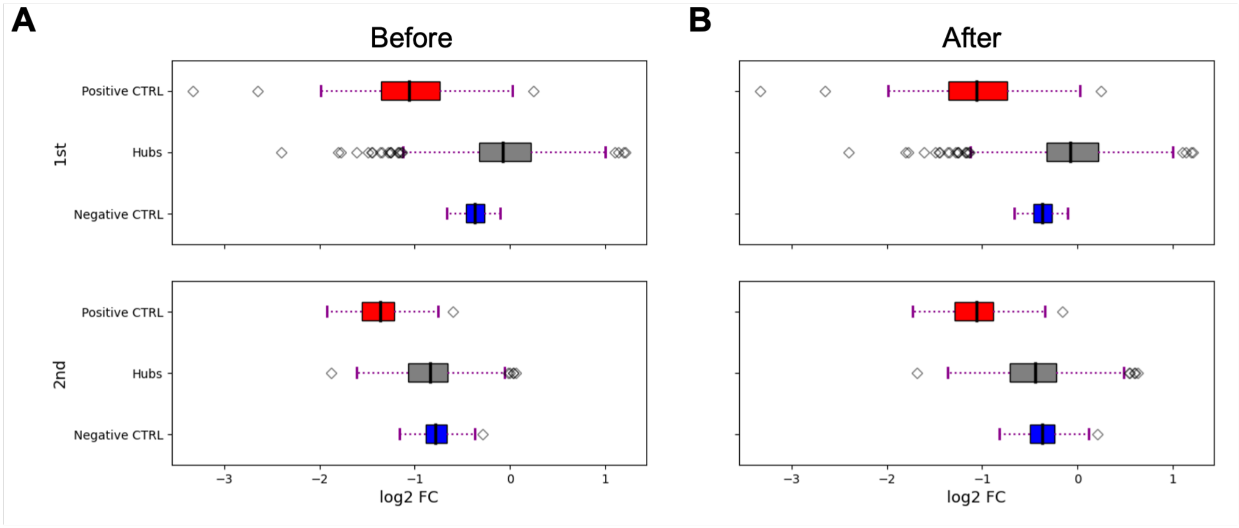


**Figure S4. Normalization of the log2FC values**

The boxplots show the distribution of the log2FC values of the gRNAs in the negative control, hub candidates, and positive control pools. The black bar in the box represents the mean value. 1st, the first round of screening in Ding et al. study. 2nd, the second round of screening in the current study. **(A)** before normalization, and **(B)** after performing linear normalization on the results of the second round in the log2 space, the mean of log2FC for positive (and negative) control pools from the both rounds of the experiment become equal.

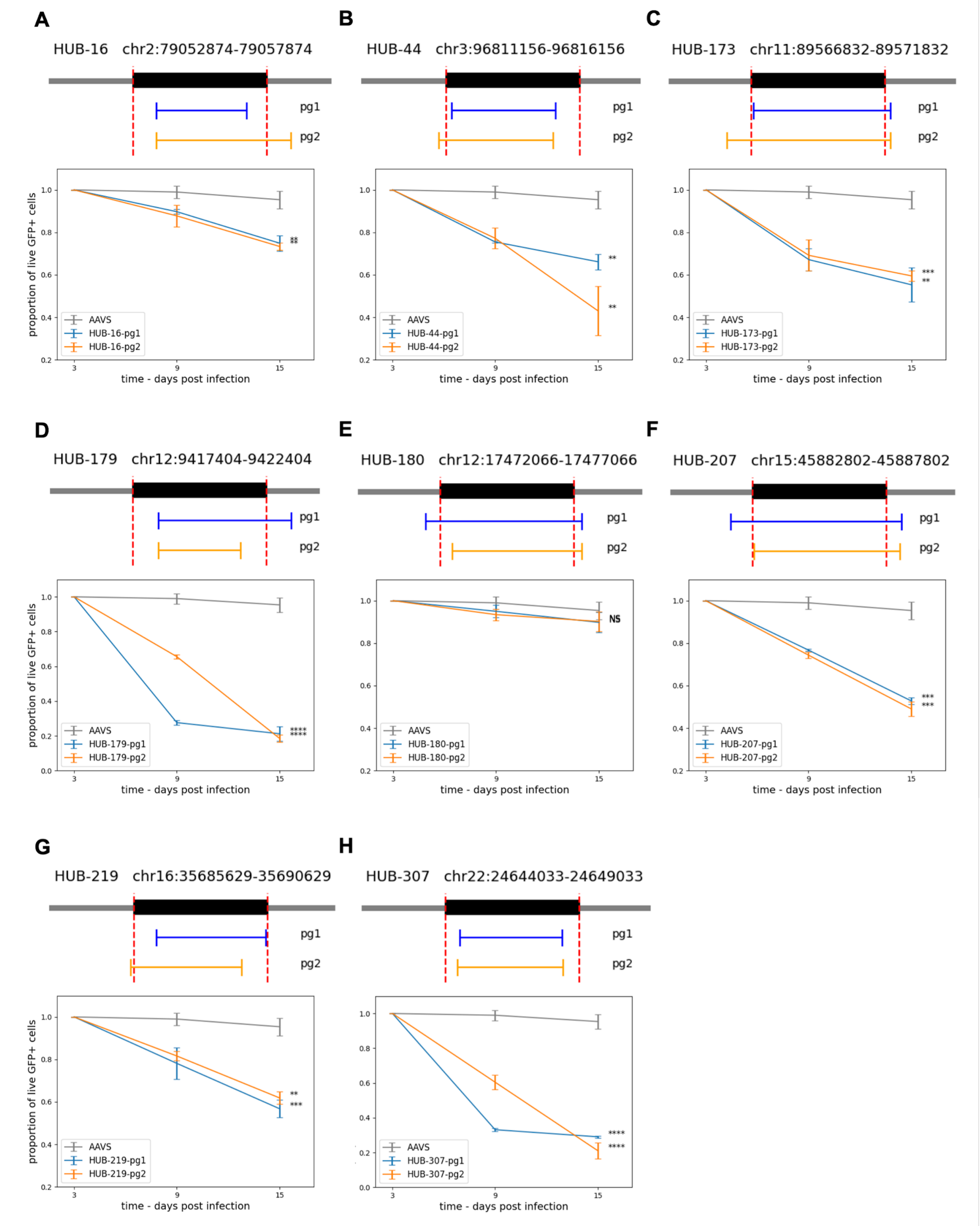


**Figure S5. Validation of the screening results by individual deletion experiments**

(**A** - **I**) Upper: Illustration of the locations of the pgRNAs targeting the selected hubs, with coordinates are in hg38. Below: time curve that indicates the survival rate of the hub-KO cells (GFP+). Data is presented as mean ± standard deviation (n = 2). Asterisks represent the *p*-values obtained from performing a two-tailed student's t-test by comparing the *FC* of hub-KO cells with that of AAVS-1 site-KO cells on the day 15. * $p < 0.05$, ** $p < 0.01$, *** $p < 0.001$, **** $p < 0.0001$, and NS means not significant. (**F**) only HUB-180 showed insignificant essentiality.

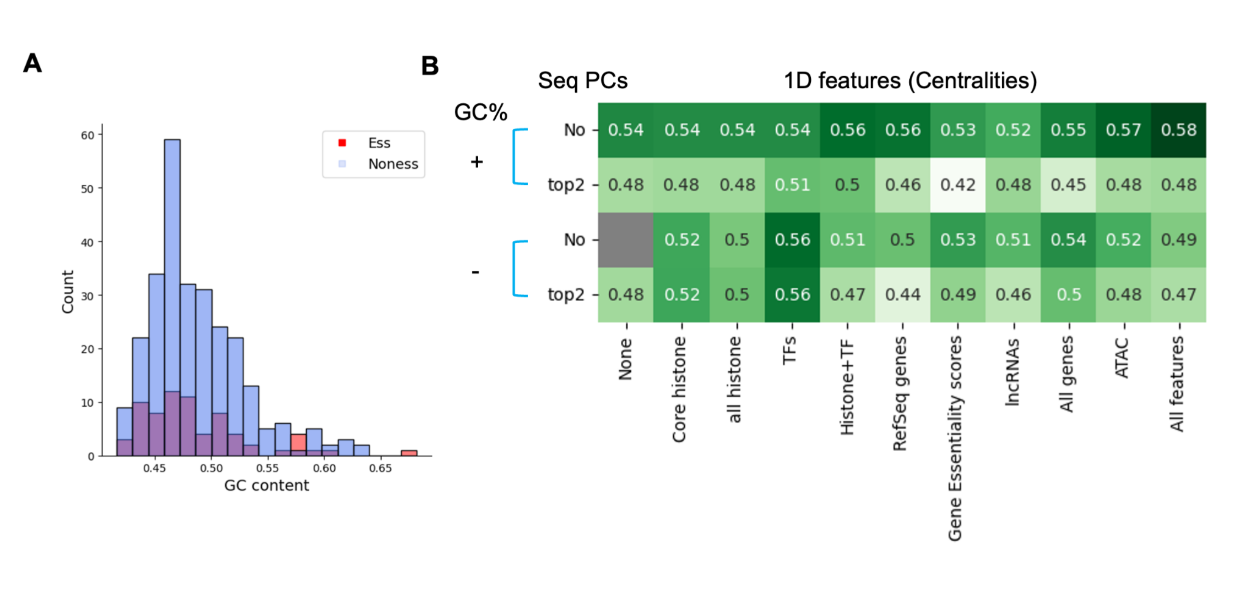


| GC% | Seq PCs | None | Core histone | all histone | TFs | Histone+TF | RefSeq genes | Gene Essentiality scores | lncRNAs | All genes | ATAC | All features |
|---|---|---|---|---|---|---|---|---|---|---|---|---|
| + | No | 0.54 | 0.54 | 0.54 | 0.54 | 0.56 | 0.56 | 0.53 | 0.52 | 0.55 | 0.57 | 0.58 |
| + | top2 | 0.48 | 0.48 | 0.48 | 0.51 | 0.5 | 0.46 | 0.42 | 0.48 | 0.45 | 0.48 | 0.48 |
| - | No | | 0.52 | 0.5 | 0.56 | 0.51 | 0.5 | 0.53 | 0.51 | 0.54 | 0.52 | 0.49 |
| - | top2 | 0.48 | 0.52 | 0.5 | 0.56 | 0.47 | 0.44 | 0.49 | 0.46 | 0.5 | 0.48 | 0.47 |

**C**

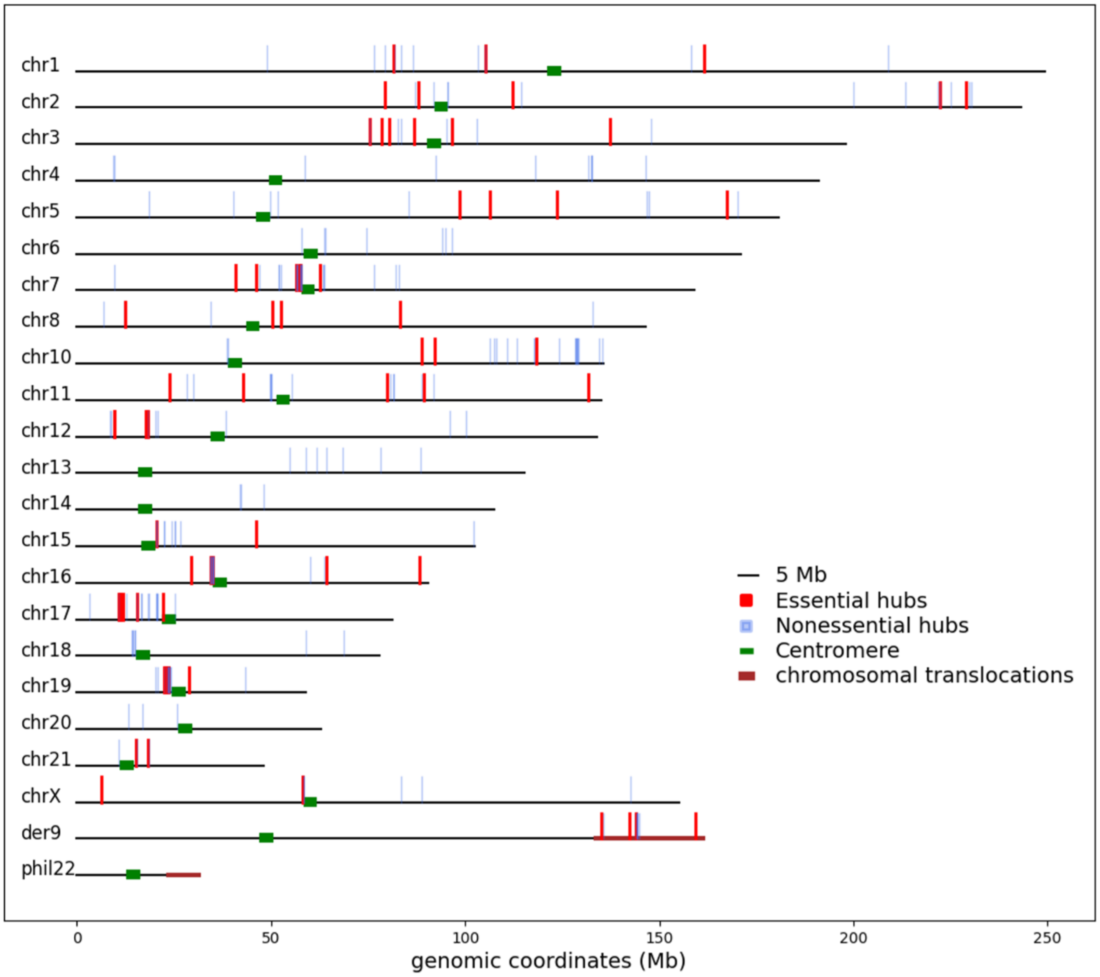


**Figure S6 Essential and nonessential hubs cannot be distinguished by sequence or epigenetic features**

**(A)** Distribution of GC contents for the essential and nonessential hubs identified in the current screening. The GC content of the essential hubs is not significantly different from that of the nonessential hubs ($p$ = 0.430 by Mann-Whitney U test).

(**B**) Performance of different feature combinations. The numbers show ROC scores. The figure suggests no prediction power since all values are below 0.6.

(**C**) Hub loci (hg19) on each chromosome. Vertical lines above the chromosomes represent the locations of the hubs. Among them, red lines represent essential hubs, and blue lines represent nonessential hubs. Green blocks on the chromosomes represent the locations of centromeres. In the K562 cell line, translocation occurs between chr9 and chr22, resulting in the elongated der9 and truncated phil22. Brown bars show the locations of reciprocal translocation. Green horizontal blocks on the chromosomes represent the locations of centromeres.

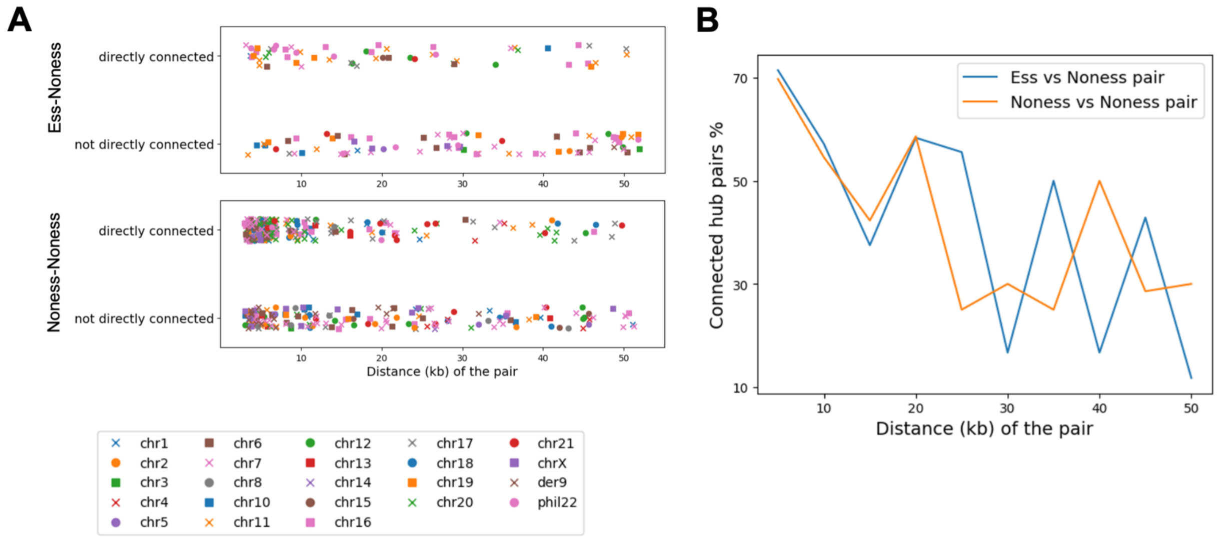
A
Ess-Noness
directly connected
not directly connected
Noness-Noness
directly connected
not directly connected
Distance (kb) of the pair
chr1
chr2
chr3
chr4
chr5
chr6
chr7
chr8
chr10
chr11
chr12
chr13
chr14
chr15
chr16
chr17
chr18
chr19
chr20
chr21
chrX
der9
phil22
B
Ess vs Noness pair
Noness vs Noness pair
Connected hub pairs %
Distance (kb) of the pair

C

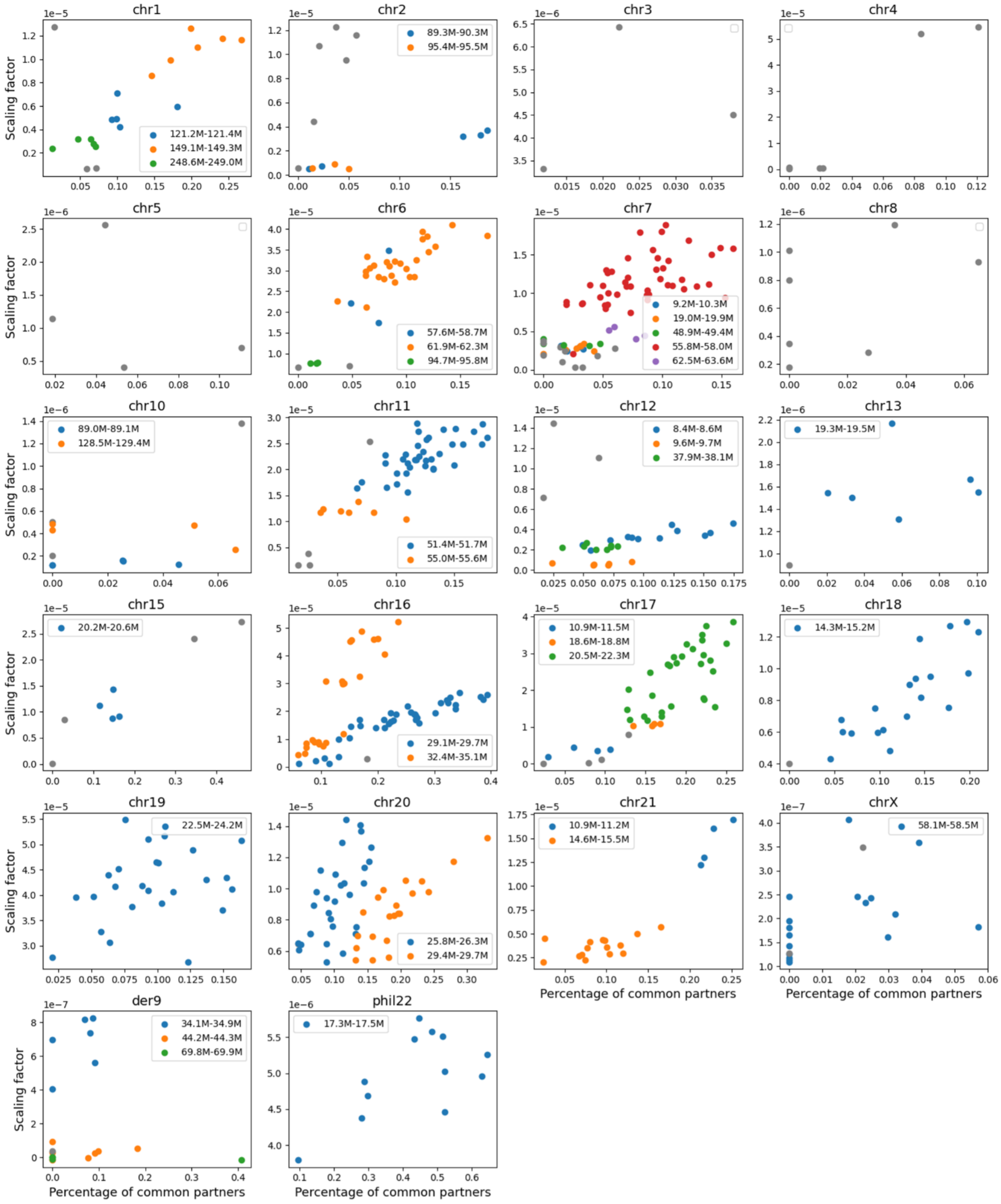

chr1
121.2M-121.4M
149.1M-149.3M
248.6M-249.0M
chr2
89.3M-90.3M
95.4M-95.5M
chr3
chr4
chr5
chr6
57.6M-58.7M
61.9M-62.3M
94.7M-95.8M
chr7
9.2M-10.3M
19.0M-19.9M
48.9M-49.4M
55.8M-58.0M
62.5M-63.6M
chr8
chr10
89.0M-89.1M
128.5M-129.4M
chr11
51.4M-51.7M
55.0M-55.6M
chr12
8.4M-8.6M
9.6M-9.7M
37.9M-38.1M
chr13
19.3M-19.5M
chr15
20.2M-20.6M
chr16
29.1M-29.7M
32.4M-35.1M
chr17
10.9M-11.5M
18.6M-18.8M
20.5M-22.3M
chr18
14.3M-15.2M
chr19
22.5M-24.2M
chr20
25.8M-26.3M
29.4M-29.7M
chr21
10.9M-11.2M
14.6M-15.5M
chrX
58.1M-58.5M
der9
34.1M-34.9M
44.2M-44.3M
69.8M-69.9M
phil22
17.3M-17.5M
Scaling factor
Percentage of common partners

**D**

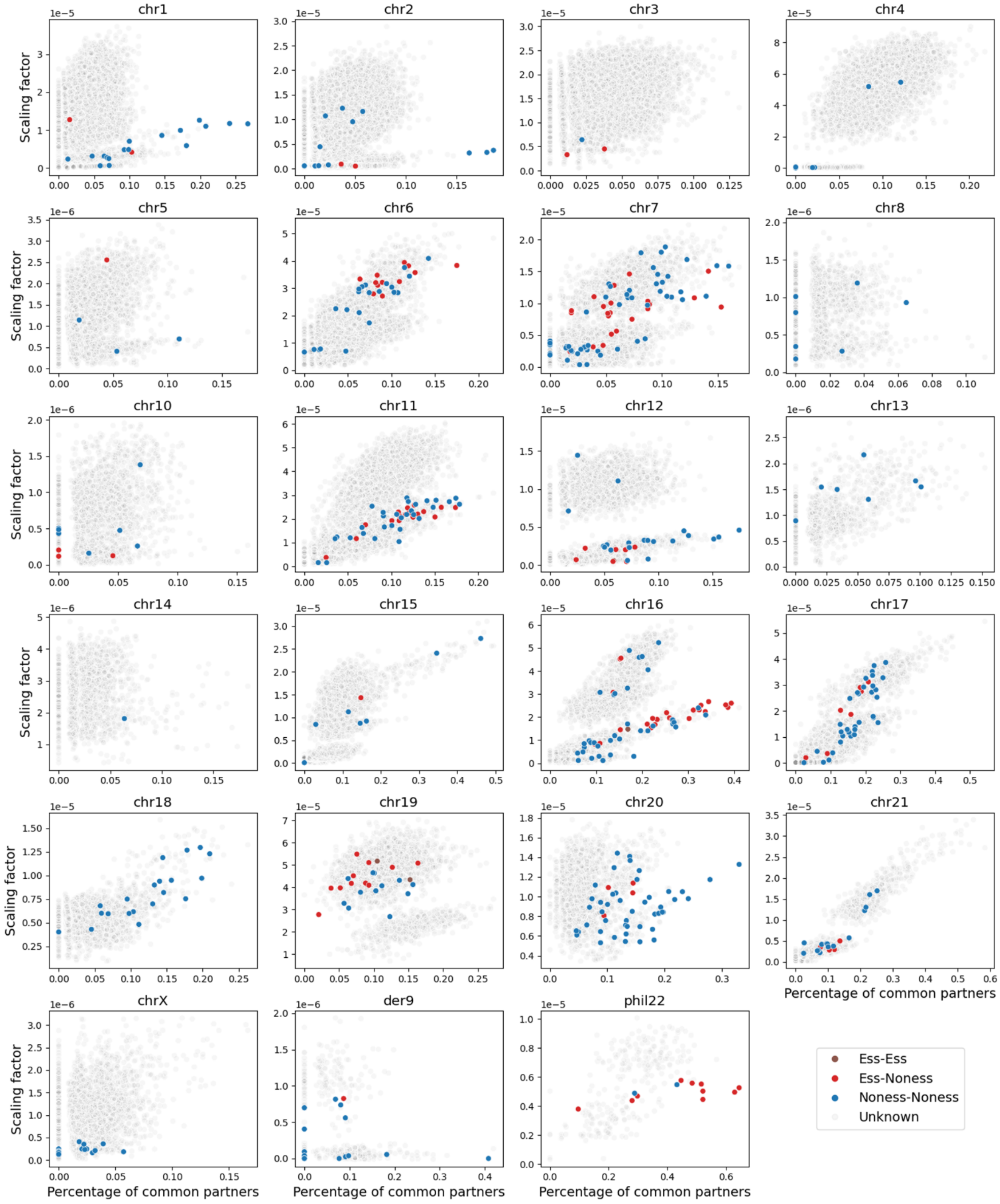
chr1
chr2
chr3
chr4
chr5
chr6
chr7
chr8
chr10
chr11
chr12
chr13
chr14
chr15
chr16
chr17
chr18
chr19
chr20
chr21
chrX
der9
phil22
Scaling factor
Percentage of common partners
Ess-Ess
Ess-Noness
Noness-Noness
Unknown

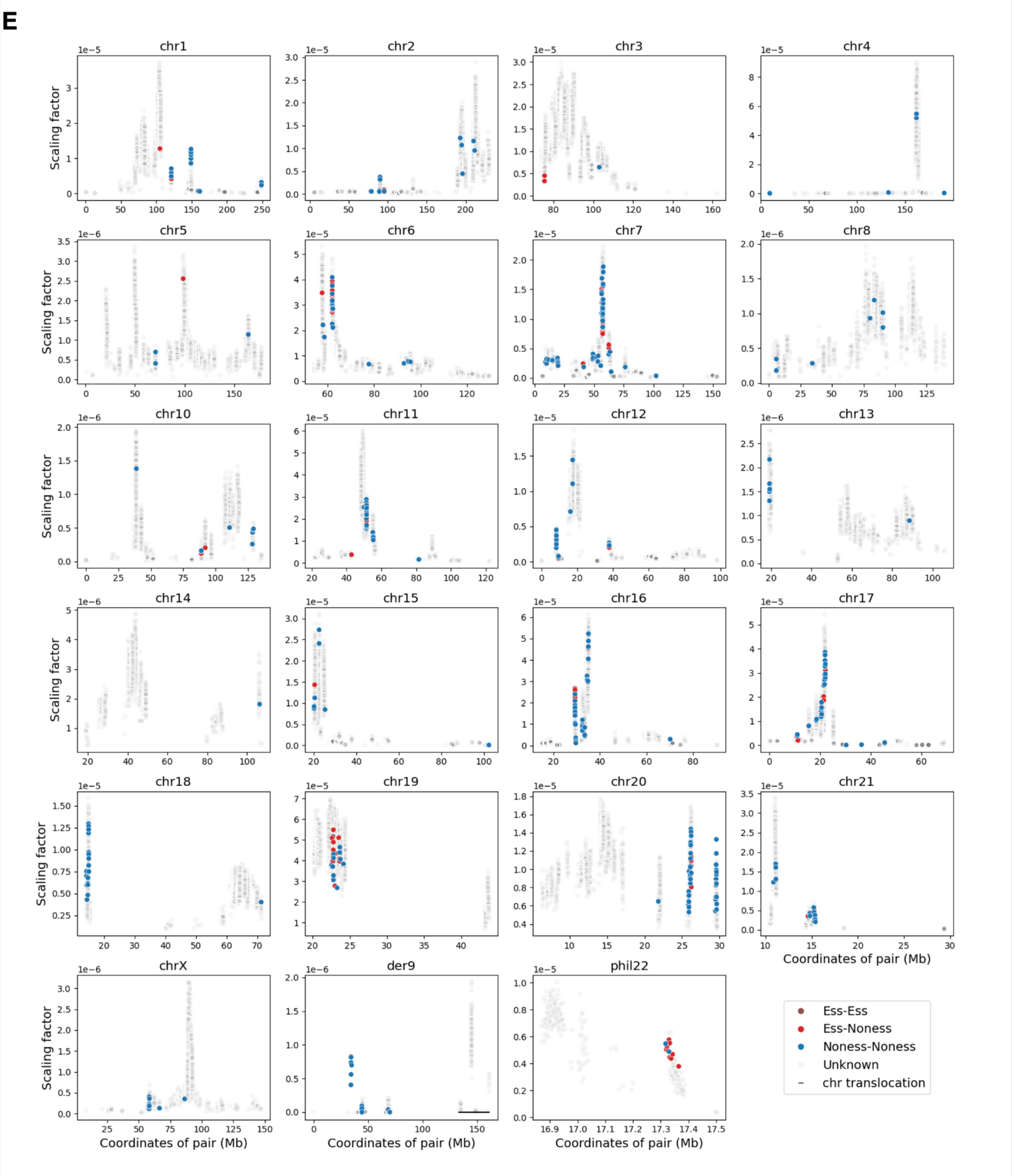


**Figure S7 Connectivity analysis in the FCN on the nearby hub pairs**

**(A)** Scatter plots show hub pairs at different distances on different chromosomes.

**(B)** Path analysis with step = 1, i.e. direct connections. The X-axis represents the distance (kb) between hub pairs in the linear genome, and the Y-axis represents the percentage of directly connected hub pairs among all the identified hub pairs. Hub pairs that are closer in the linear genome are more likely to form direct contacts than hub pairs that are distant in the linear genome. Pearson *r* = -0.716, for essential-nonessential hub pairs, and Pearson *r* = -0.684, for nonessential-nonessential hub pairs.

**(C)** Analysis of paths with step numbers of 2 to 4, i.e. percentage of common interacting partners and scaling factors (see Methods). Scatter plots show the pairs of hubs tested in Ding et al. and this study, on different chromosomes. Hub pairs are colored in groups based on their genomic coordinates.

**(D)** Scatter plots showing all possible pairs among 26,148 hubs in K562. X-axis: percentage of common interacting partners, and Y-axis: scaling factors. Each dot represents a hub pair. Unknown: gray dots in the background representing the hub pairs where at least one hub was not tested in either Ding et al. or this study.

**(E)** Scatter plots showing all possible pairs among 26,148 hubs in K562. X-axis: the genomic coordinates of the pair (mean of the hubs), and Y-axis: scaling factors.

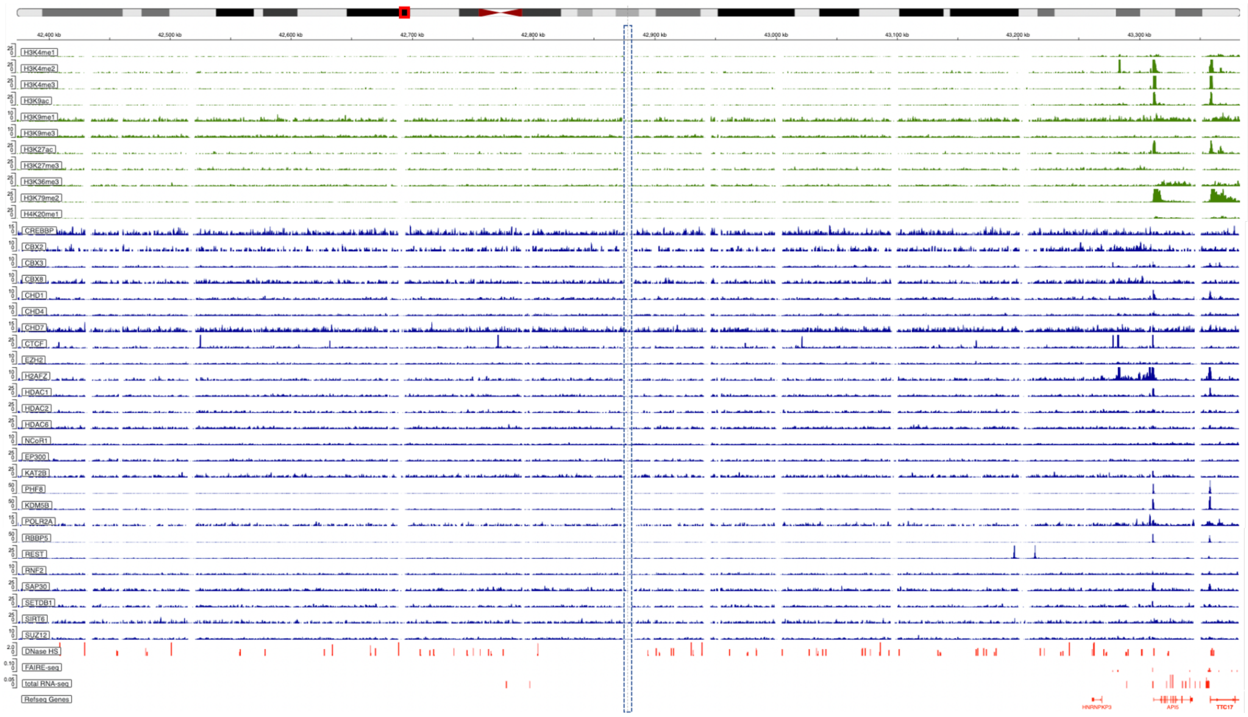


**Figure S8 Genome browser view of annotations and epigenetic signals around the nonessential HUB-161 (chr11:42,873,450-42,878,450, hg38) and essential HUB-162 (chr11:42,878,450-42,883,450, hg38)**

The central 10-kb region within the dashed box represents HUB-161 (Nonessential, left, 5-kb) and HUB-162 (Essential, right, 5-kb).

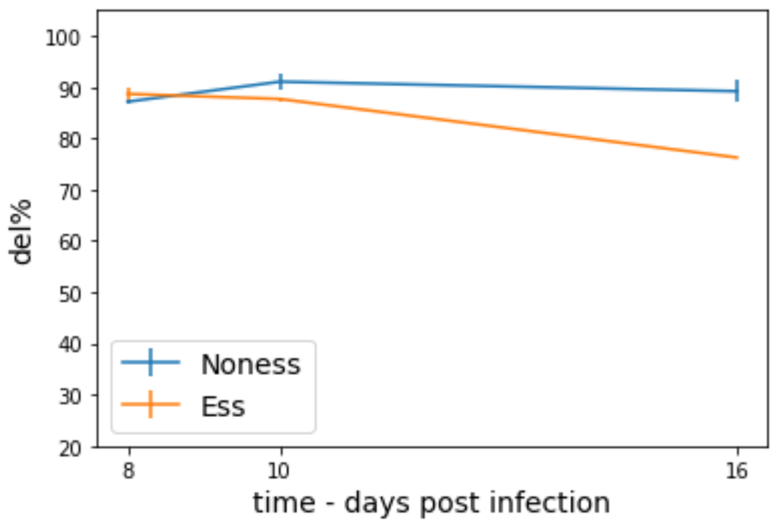


**Figure S9 Deletion efficiency measured by pPCR on day 8, 10, and 16 post infection.** The essential hub-targeting pgRNA (Ess-pg2) consistently showed comparable deletion to that of the nonessential hub-targeting one (Noness-pg4)

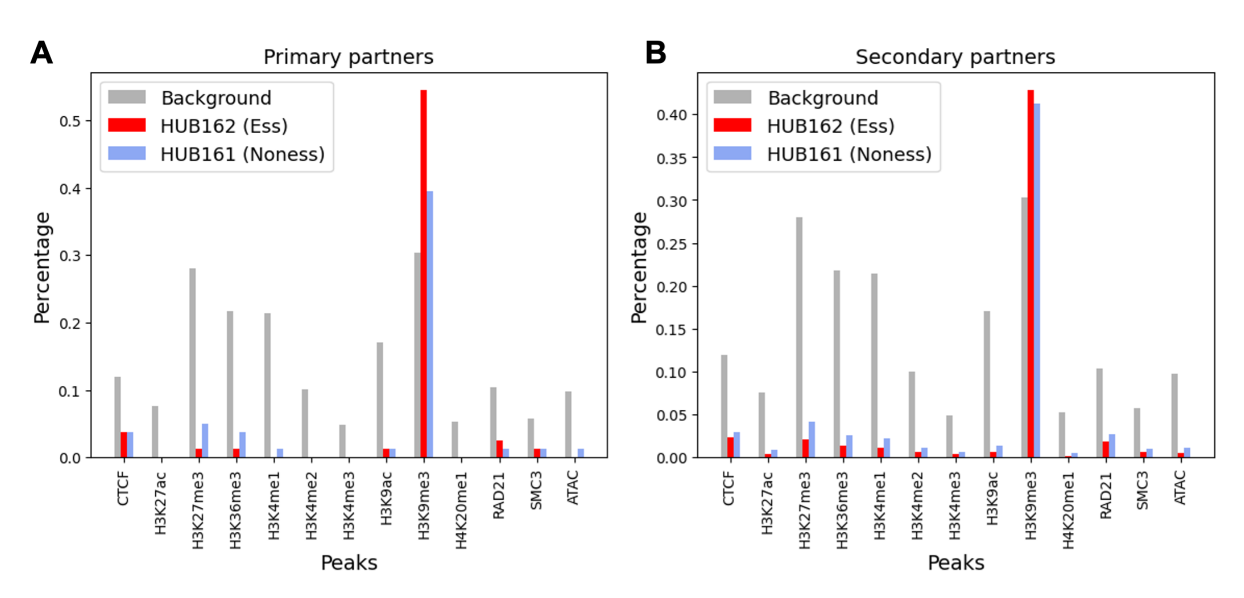


**Figure S10 Comparison of interacting nodes of HUB162 vs HUB161**

**(A)** Barplots show the percentages of nodes overlapping with the ChIP-seq peaks of histone modifications and TFs, as well as open chromatin peaks, for the 5-kb nodes directly interacting with HUB-162 (Essential, red) or HUB-161 (Nonessential, blue). Gray bars represent the background values for all nodes on chr11.

**(B)** Barplots show the values for the secondary partners linked to the hub through an intermediate node.

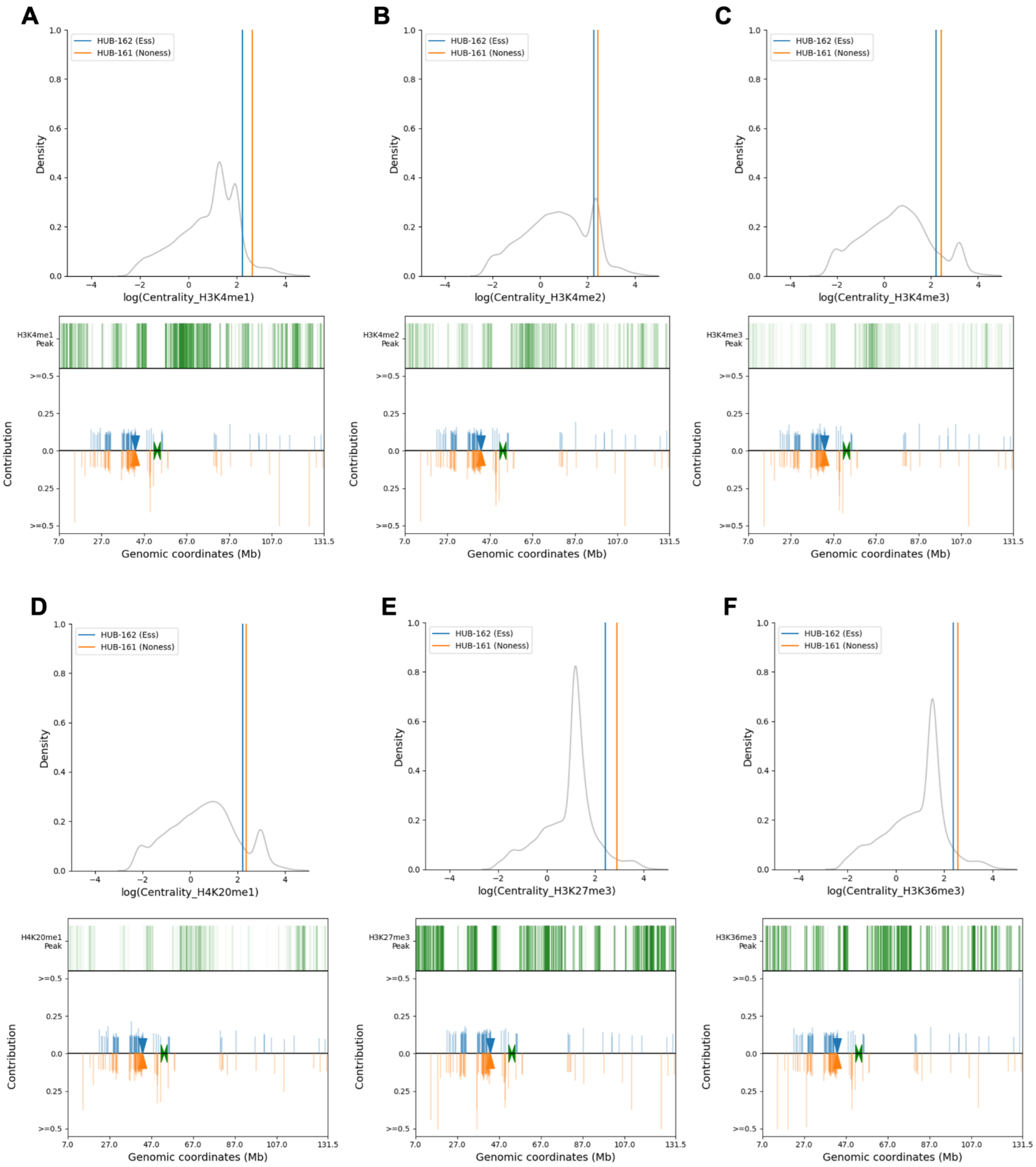
A
B
C
D
E
F
HUB-162 (Ess)
HUB-161 (Noness)
Density
log(Centrality_H3K4me1)
log(Centrality_H3K4me2)
log(Centrality_H3K4me3)
log(Centrality_H4K20me1)
log(Centrality_H3K27me3)
log(Centrality_H3K36me3)
H3K4me1 Peak
H3K4me2 Peak
H3K4me3 Peak
H4K20me1 Peak
H3K27me3 Peak
H3K36me3 Peak
Contribution
>=0.5
0.25
0.0
Genomic coordinates (Mb)
7.0
27.0
47.0
67.0
87.0
107.0
131.5

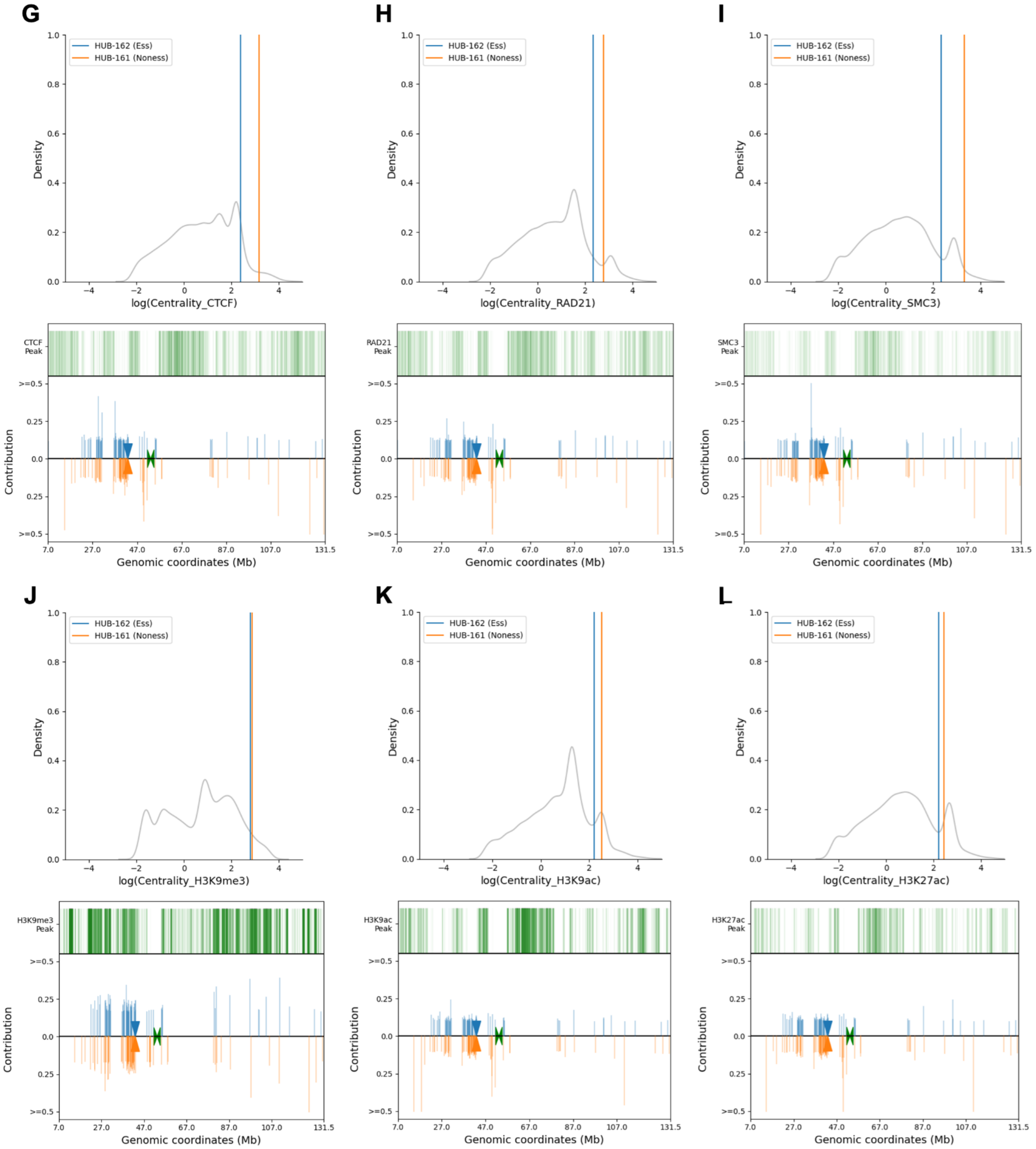
G
H
I
J
K
L
HUB-162 (Ess)
HUB-161 (Noness)
Density
log(Centrality_CTCF)
log(Centrality_RAD21)
log(Centrality_SMC3)
log(Centrality_H3K9me3)
log(Centrality_H3K9ac)
log(Centrality_H3K27ac)
CTCF Peak
RAD21 Peak
SMC3 Peak
H3K9me3 Peak
H3K9ac Peak
H3K27ac Peak
Contribution
Genomic coordinates (Mb)

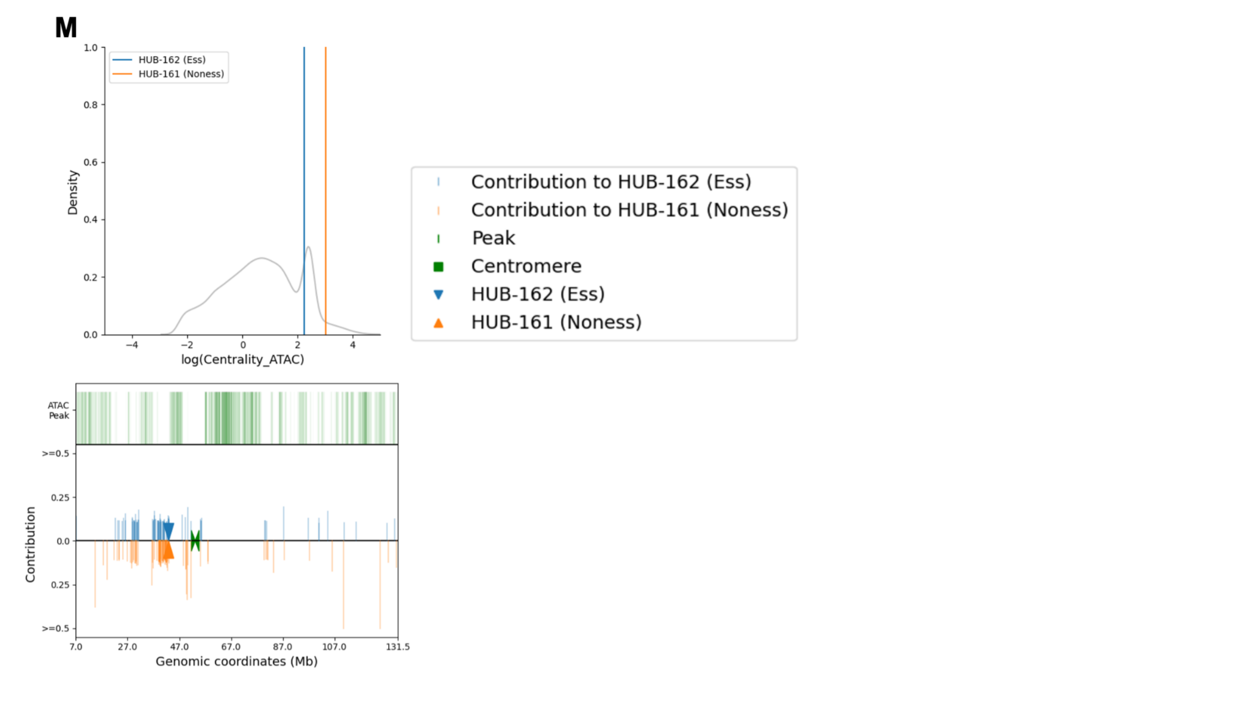


**Figure S11 PageRank scores of epigenetic features**

Each distribution figure displays the log(PageRank scores) of the epigenetic feature in K562 WT cells. **(A)** H3K4me1, **(B)** H3K4me2, **(C)** H3K4me3, **(D)** H4K20me1, **(E)** H3K27me3, **(F)** H3K36me3, **(G)** CTCF, **(H)** RAD21, **(I)** SMC3, **(J)** H3K9me3, **(K)** H3K9ac, **(L)** H3K27ac, and **(M)** ATAC. The gray background represents all 5-kb genomic loci on chr11. The blue line represents the value of HUB-162 (Essential) on the X-axis, and the orange line represents the value of HUB-161 (Nonessential) on the X-axis. Each genome browser track shows the contribution from each node to the PageRank scores of HUB-162 (blue, Essential) and HUB-161 (orange, Nonessential). Coordinates are in hg19.

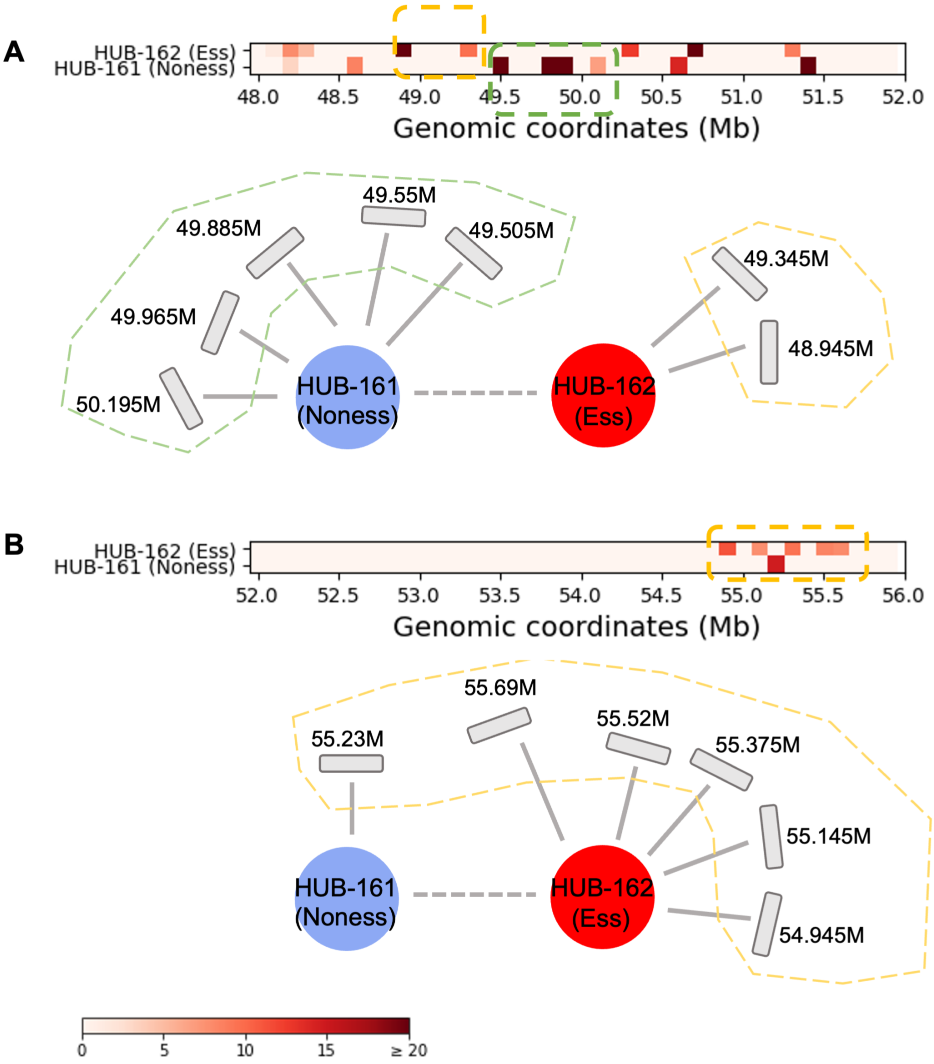


**Figure S12 Distinct 3D contact communities formed by the hub pair HUB-161 and HUB-162**

Each heatmap displays the VC normalized Hi-C read counts in the 100-kb bins that contact HUB-162 (upper, Essential) or HUB-161 (lower, Nonessential). Each cartoon illustrates the local connections to each hub in different communities (dashed boxes). **(A)** chr11:48M-52M, and **(B)** chr11:52M-56M. Coordinates are shown in hg19.

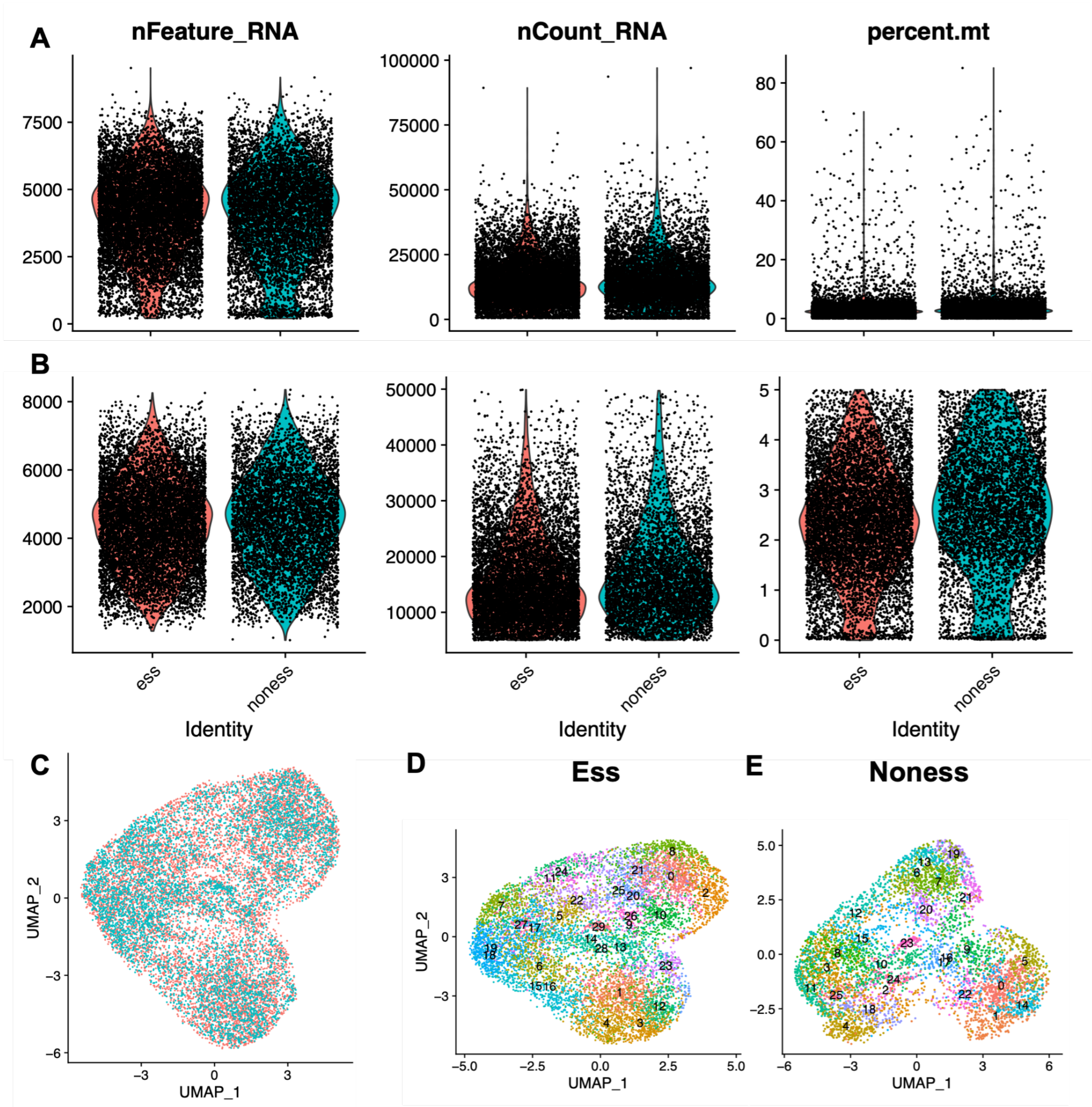


**Figure S13 scRNA-seq quality control results for the HUB-162 (Essential) and HUB-161 (Nonessential) deletion samples**

**(A, B)** Quality control (QC) plots for three metrics (i) the number of detected RNAs, (ii) the total counts of RNAs, and (iii) the percentage of mitochondrial RNAs. Each black dot represents a single cell (barcode). The red violin plots represent the distributions of the HUB-162 deletion sample, and the blue violin plots represent HUB-161 deletion. **(A)** before and **(B)** after filtering out the low-quality single cells.

(**C**) UMAP visualization of the filtered single cells in the scRNA-seq experiments. Single cells from the HUB-162 and HUB-161 deletion experiments are mixed with each other, showing no observable batch effects.

(**D, E**) UMAP visualization plots of the single cells in the scRNA-seq experiments. Colors and numbers indicate the clusters identified by SNN modularity optimization (Seurat v4.1.0). (**D**) HUB-162 (Essential) and (**E**) HUB-161 (Nonessential).

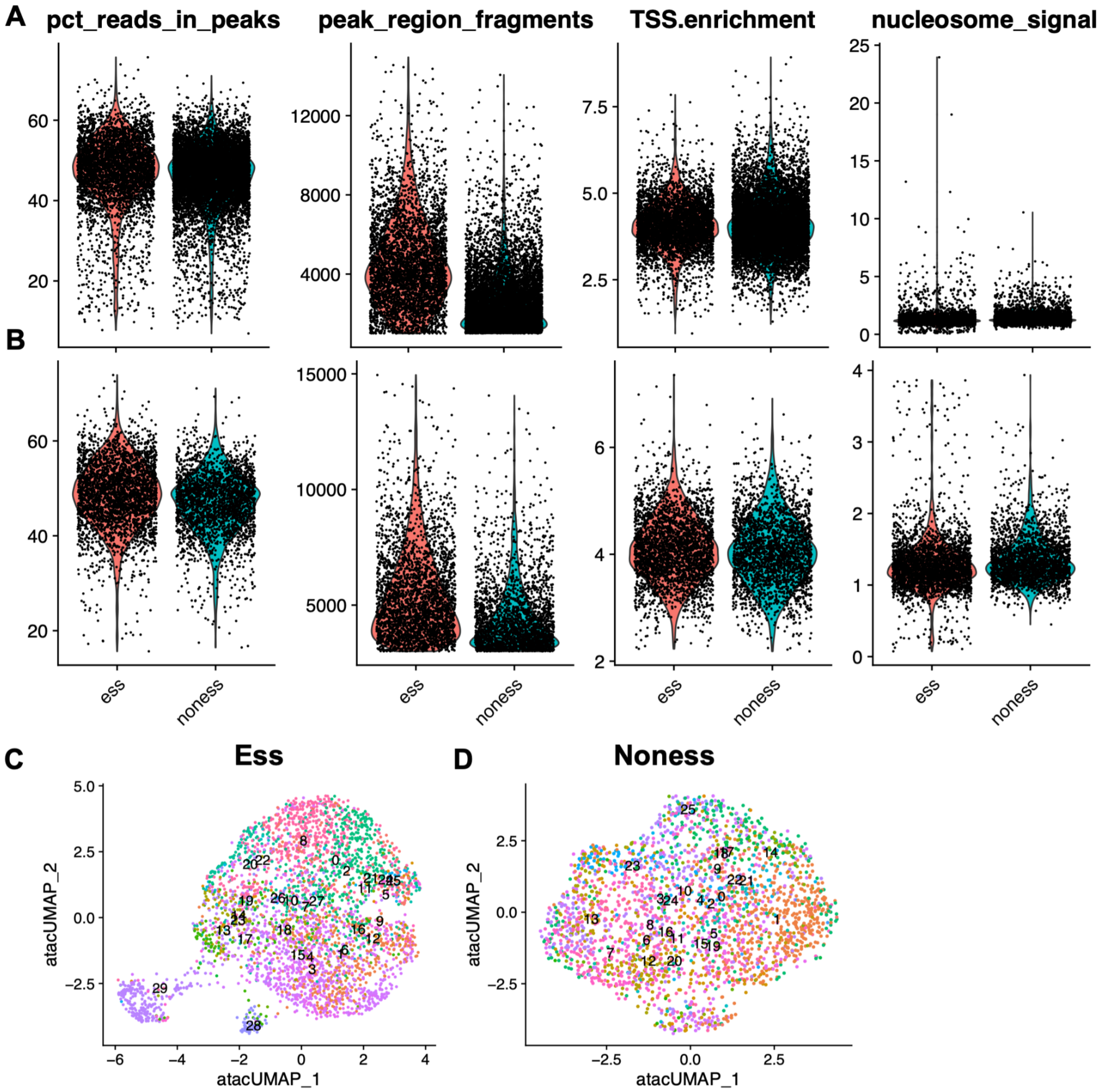


**Figure S14 scATAC-seq quality control results for the HUB-162 (Essential) and HUB-161 (Nonessential) deletion samples**

**(A, B)** Quality control (QC) plots for four metrics (i) percentage of reads in peaks, (ii) the number of peak region fragments, (iii) the transcription start site (TSS) enrichment fold, and (iv) nucleosome signals. Each black dot represents a single cell (barcode). The red violin plots represent the distributions of the HUB-162 deletion sample, and the blue plots represent HUB-161 deletion. **(A)** before and **(B)** after filtering out the low-quality single cells.

(**C, D**) UMAP visualization plots of single cells in the scATAC-seq, numbers indicate the clusters anchored to the corresponding clusters in the scRNA-seq analysis. (**C**) HUB-162 (Essential) and (**D**) HUB-161 (Nonessential).

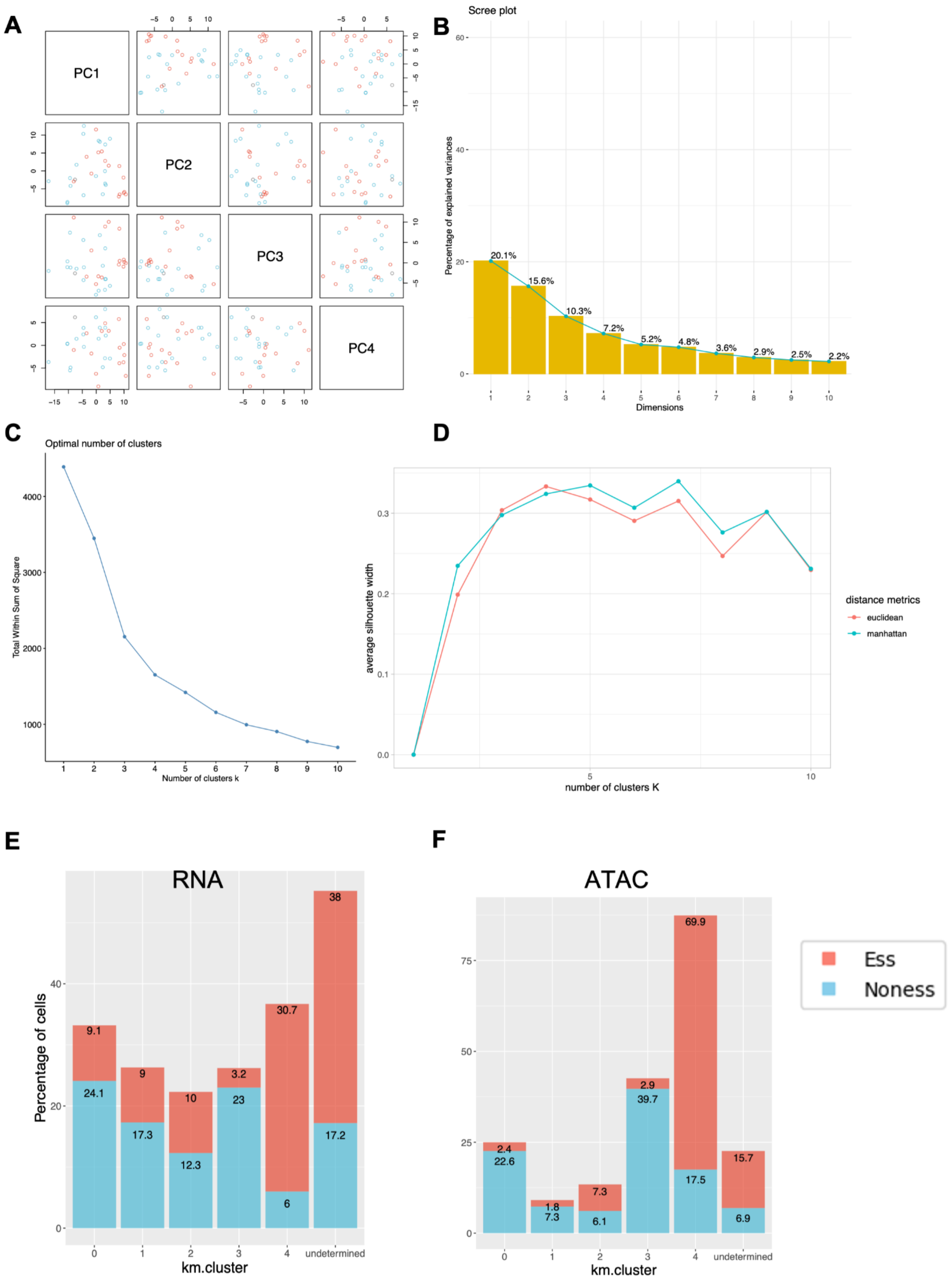
A
PC1
PC2
PC3
PC4
B
Scree plot
Percentage of explained variances
Dimensions
20.1%
15.6%
10.3%
7.2%
5.2%
4.8%
3.6%
2.9%
2.5%
2.2%
C
Optimal number of clusters
Total Within Sum of Square
Number of clusters k
D
average silhouette width
number of clusters K
distance metrics
euclidean
manhattan
E
RNA
Percentage of cells
km.cluster
0
1
2
3
4
undetermined
9.1
24.1
9
17.3
10
12.3
3.2
23
30.7
6
38
17.2
F
ATAC
km.cluster
2.4
22.6
1.8
7.3
7.3
6.1
2.9
39.7
69.9
17.5
15.7
6.9
Ess
Noness

**Figure S15 Data processing for the Taiji analysis**

**(A)** The top four principal components (PCs) of the TFs' PageRank scores generated by Taiji in the pseudobulk samples. Gray: WT, red: HUB-162 deletion (Essential), and blue: HUB-161 deletion (Nonessential).

**(B)** The percentage of explained variance with the top 10 PCs.

**(C)** The elbow method to determine the optimal *K* value in the K-Means clustering using the top four PCs, and the optimal *K* = 5.

**(D)** The Silhouette score method to determine the optimal *K* value in the K-Means clustering, with the optimal *K* = 4 when using Euclidean distance, and *K* = 5 when using Manhattan distance.

**(E, F)** Percentage of cells in each km.cluster in the **(E)** scRNA-seq and **(F)** scATAC-seq experiment. Numbers on the bars indicate number_in_cluster / number_total. The tag "Undetermined" means cells in the clusters excluded for Taiji inputs (refer to Methods).

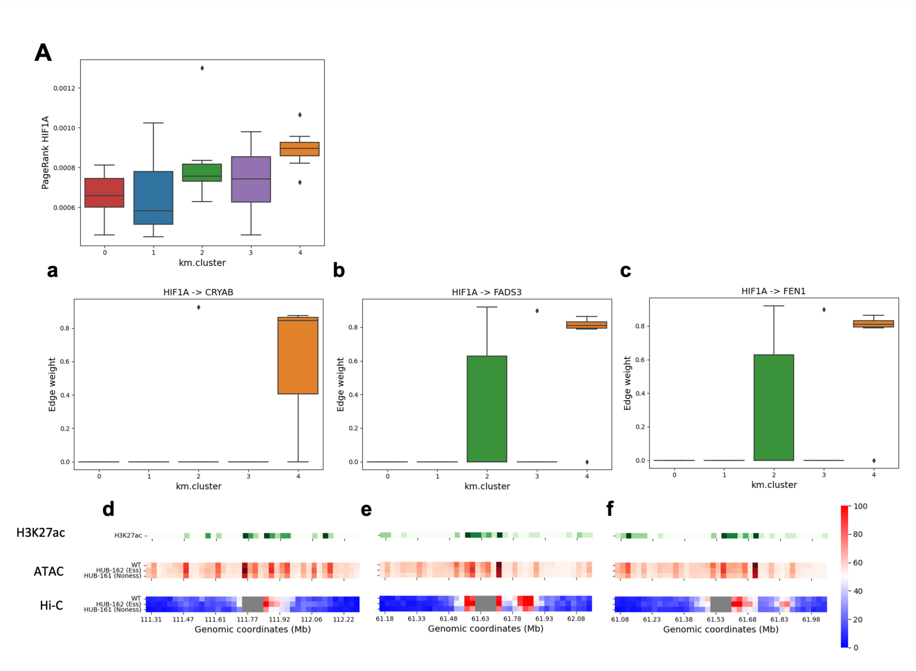
A
PageRank HIF1A
km.cluster
a
b
c
HIF1A -> CRYAB
HIF1A -> FADS3
HIF1A -> FEN1
Edge weight
km.cluster
d
e
f
H3K27ac
ATAC
Hi-C
WT
HUB-162 (Ess)
HUB-161 (Noness)
Genomic coordinates (Mb)

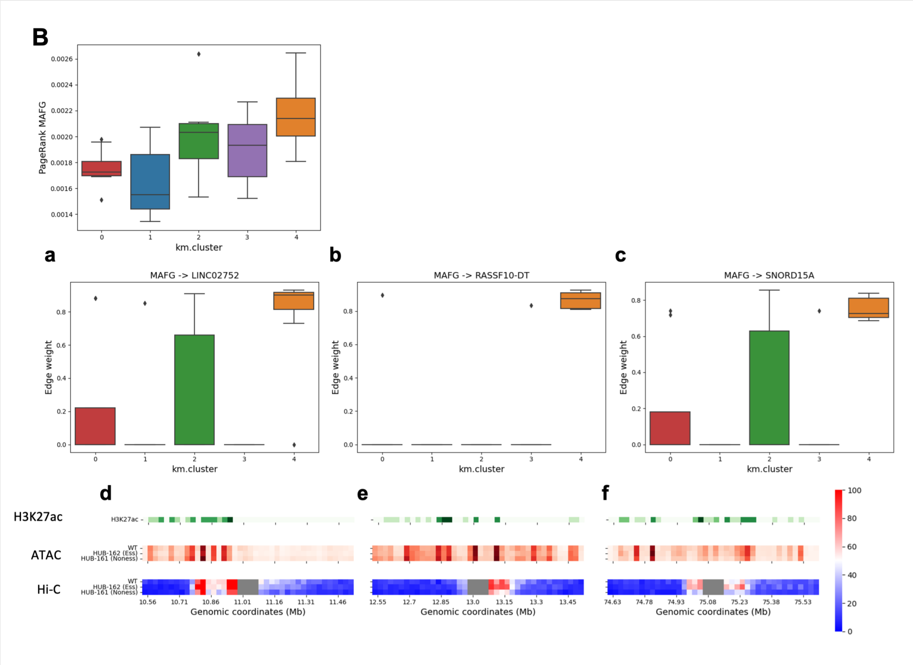
B
PageRank MAFG
km.cluster
a
b
c
MAFG -> LINC02752
MAFG -> RASSF10-DT
MAFG -> SNORD15A
Edge weight
km.cluster
d
e
f
H3K27ac
ATAC
Hi-C
WT
HUB-162 (Ess)
HUB-161 (Noness)
Genomic coordinates (Mb)

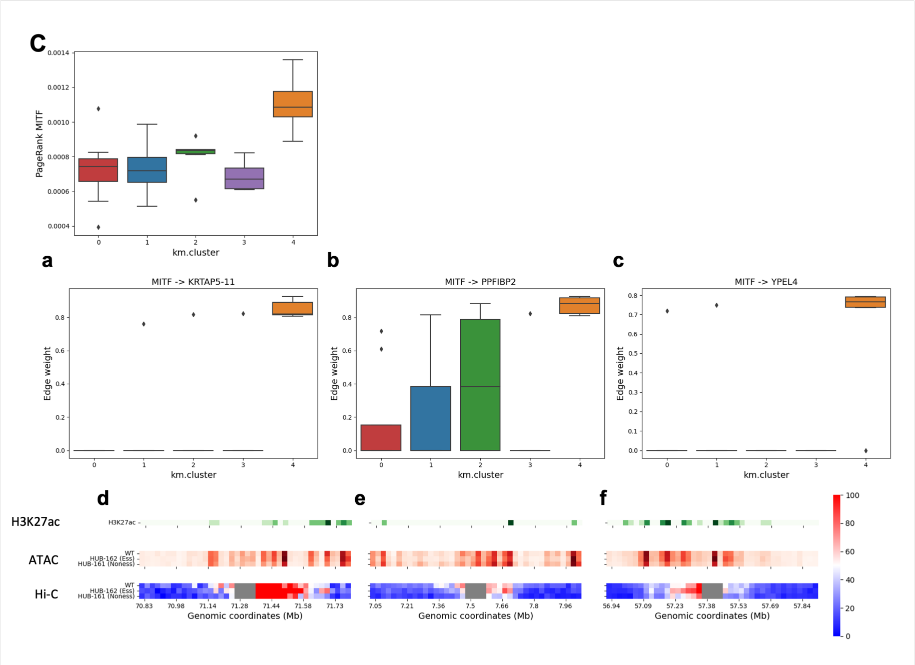


**Figure S16 PageRank scores and the associated edges of the cluster 4-specific (Essential) TFs**

**(A)** HIF1A, **(B)** MAFG, and **(C)** MITF.

**(a ~ c)** Three example edges (TF -> regulatee genes on chr11) with significantly high weights in cluster 4 (Essential) and low weights in both cluster 0 (WT) and 3 (Nonessential).

**(d ~ f)** The genome browser tracks show the corresponding H3K27ac peaks in the K562 WT; and ATAC peaks and Hi-C normalized reads in WT, HUB-162 deletion (Essential), and HUB-161 deletion (Nonessential) samples, in the 1-Mb proximity of the regulatee genes. The resolution is 25 kb, and coordinates are in hg19.

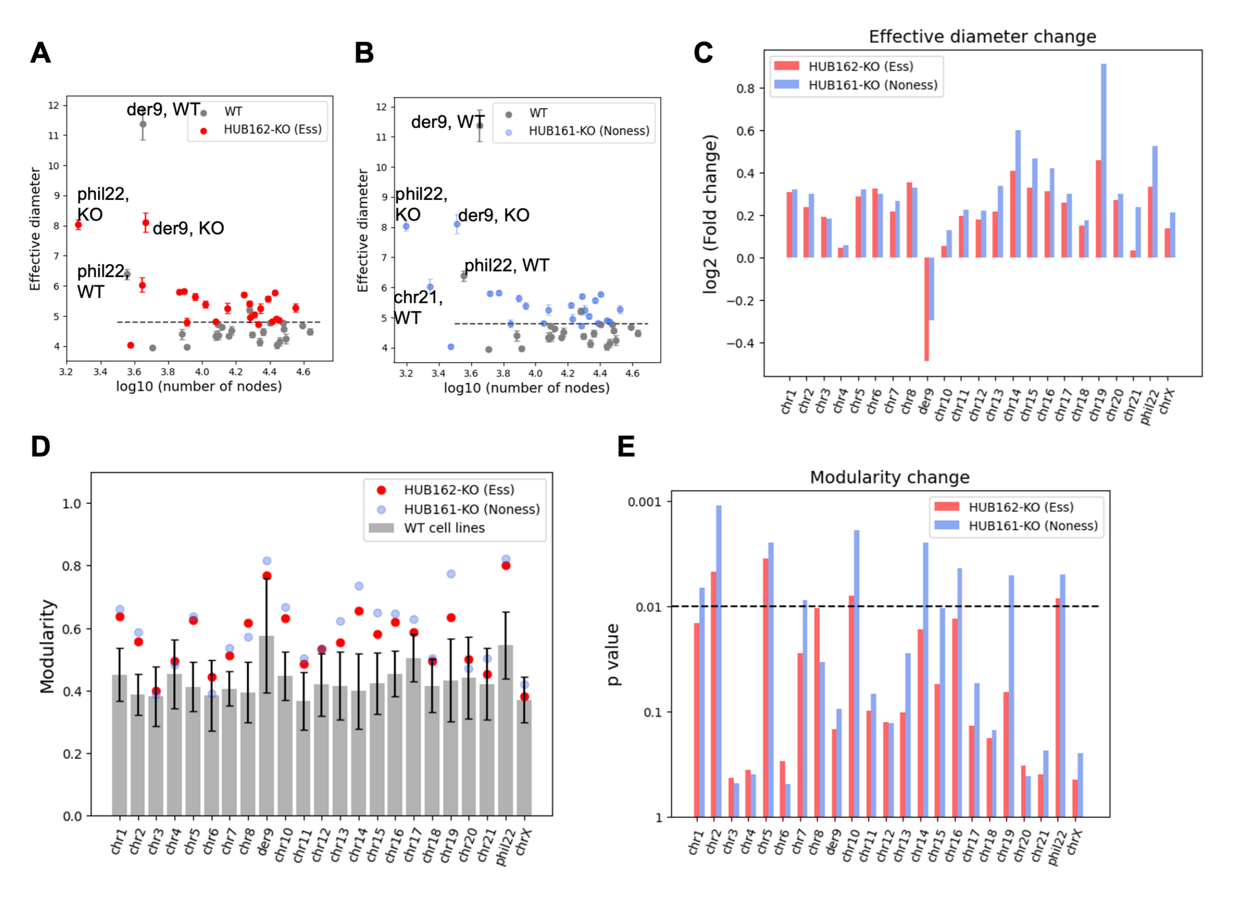


**Figure S17 FCN analysis on the hub-KO Hi-C experiments**

**(A, B)** Effective diameters vs log10 (number of nodes). Diameter is represented as mean ± standard deviation (with 10 different initializations) for each chromosome. Effective diameters universally increase (except der9) with either HUB-162 deletion (Essential, red) or HUB-161 deletion (Nonessential, blue).

**(C)** log2 (Fold change) of effective diameter on each chromosome.

**(D)** Modularities of each chromosome in 7 WT cell lines (gray bar), HUB-162 deletion (Essential, red), and HUB-161 deletion (Nonessential, blue).

**(E)** Barplot shows the *p*-values of the modularity gain on each chromosome upon HUB-162 deletion (Essential, red) and HUB-161 deletion (Nonessential, blue)

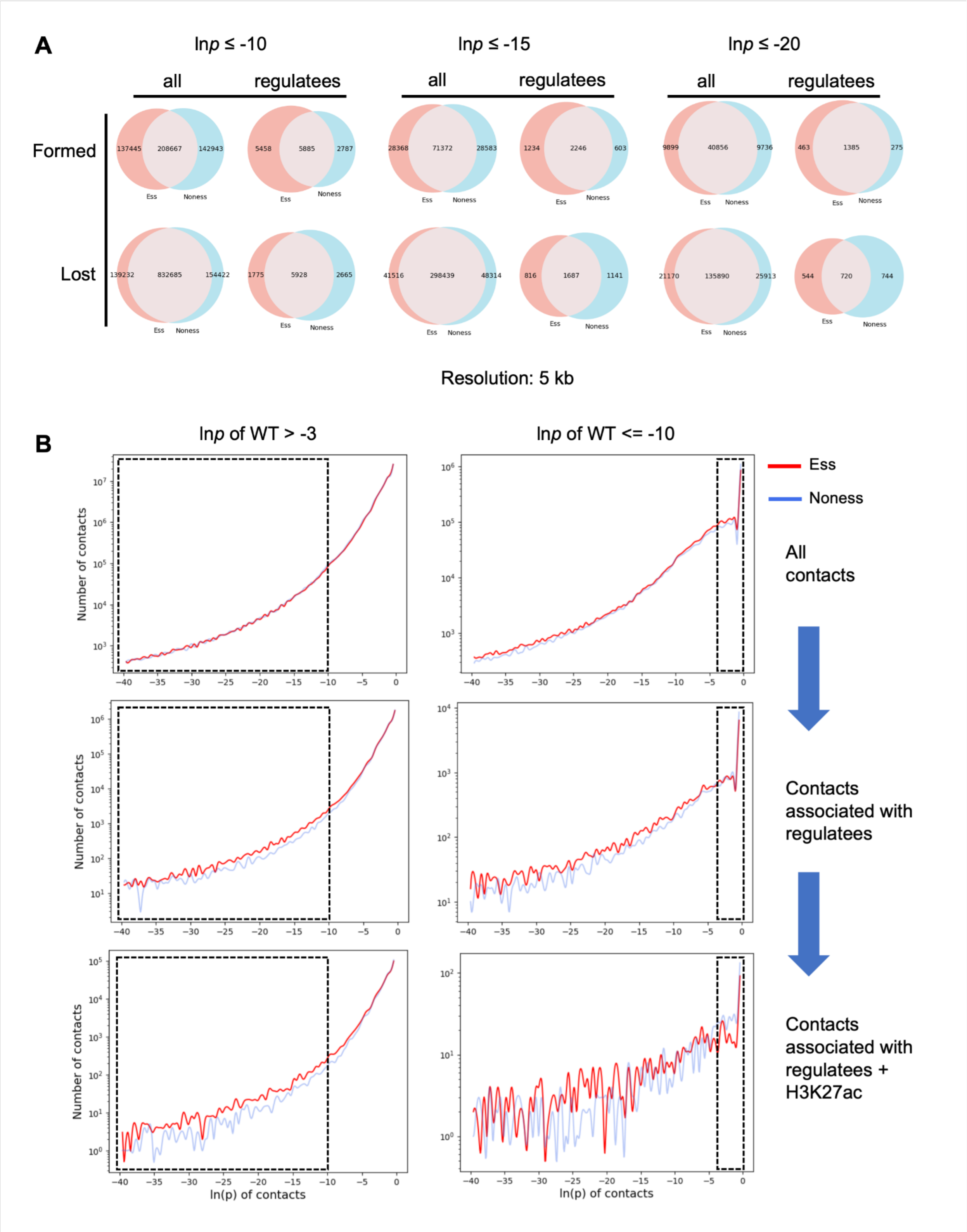


**Figure S18 Contact analysis on the bulk Hi-C experiments**

(**A**) The numbers of the newly formed and lost Hi-C contacts at 5-kb resolution upon hub deletion compared to the WT, at ln*p* cutoffs of -10, -15, and -20, respectively. Red: HUB-162 deletion (Essential). Blue: HUB-161 deletion (Nonessential). All, all Hi-C contacts in the genome. Regulatees, Hi-C contacts associated with the weight-altered edges in the Taiji analysis. Results refer to Figure 6A.
(**B**) Number of all Hi-C contacts at different ln(*p*-values). The Y-axis is in log scale. Red: HUB-162 deletion (Essential). Blue: HUB-161 deletion (Nonessential). ln*p* of WT > -3 indicates non-significant contacts in WT, and contacts in the black dashed boxes indicate newly-formed contacts. ln*p* of WT <= -10 indicates strong Hi-C contacts in WT, and contacts in the black dashed boxes indicate lost contacts. Results refer to Figure 7A ~ C.

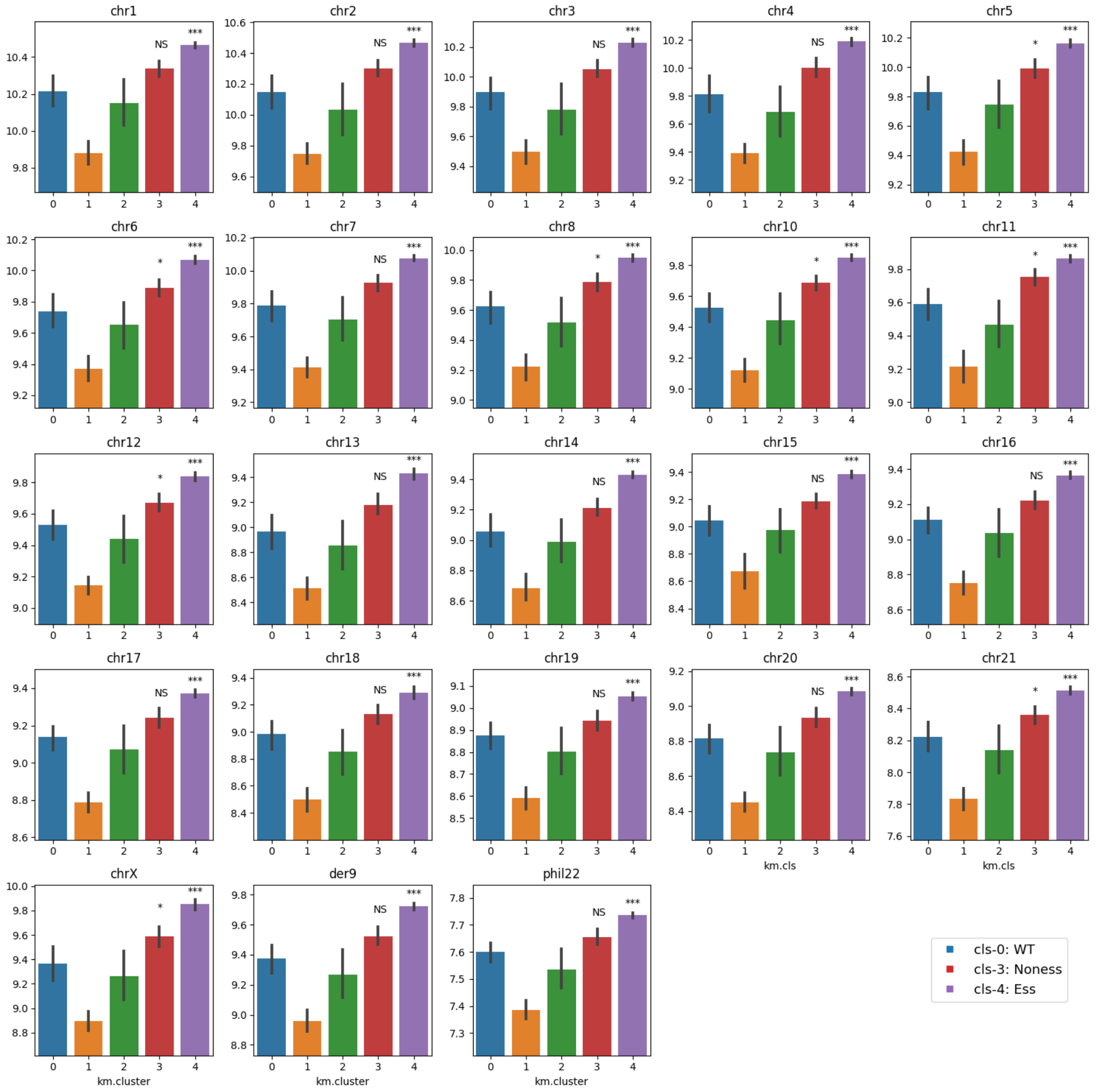


**Figure S19 Entropy of chromatin accessibility based on the scATAC-seq experiments**

Barplots show the ATAC entropy value of each K-Means cluster identified by Taiji (refer to Figure 5B) on different chromosomes. Asterisks on top of bars represent the *p*-values obtained by performing Mann-Whitney U test, comparing the entropy values of all pseudobulk samples in the K-Means cluster with those in cluster 0 (WT). * $p < 0.05$, ** $p < 0.01$, *** $p < 0.001$, and NS means not significant.

**Tables**

| Experiment | Sequencing file | Unique mapping rate |
|---|---|---|
| re1-day0 | hs0_S1_L005_R1_001.fastq | 0.84485067 |
| re1-day5 | hs5_S2_L005_R1_001.fastq | 0.84842045 |
| re1-day10 | hs10_S3_L005_R1_001.fastq | 0.85590525 |
| re1-day15 | hs15_S4_L005_R1_001.fastq | 0.7306458 |
| re1-day20 | hs20_S5_L005_R1_001.fastq | 0.7331175 |
| re1-day25 | hs25_S6_L005_R1_001.fastq | 0.65389167 |
| re2-day0 | hs0_S1_L008_R1_001.fastq | 0.9142271 |
| re2-day5 | hs5_S2_L008_R1_001.fastq | 0.91991176 |
| re2-day10 | hs10_S3_L008_R1_001.fastq | 0.92561651 |
| re2-day15 | hs15_S4_L008_R1_001.fastq | 0.77246838 |
| re2-day20 | hs20_S5_L008_R1_001.fastq | 0.78491704 |
| re2-day25 | hs25_S6_L008_R1_001.fastq | 0.66629062 |

**Table S5 Mapping rates of the CRISPR screening experiments**

| TF type | Edge type | Km.cluster type | |
|---|---|---|---|
| | | Essential | Nonessential |
| Increased pagerank scores | Stronger | 14,948 | 106 |
| | Weaker | 6 | 29 |
| Reduced pagerank scores | Stronger | 1,717 | 11 |
| | Weaker | 0 | 135 |

**Table S10 Number of significantly changed edges in Taiji analysis**

The one-way Kruskal–Wallis test was performed on the edge weight percentiles, adjusted $p$-values < 0.05. The edges were firstly grouped by “TF type” based on the TFs’ (starting nodes) *FC* of the PageRank scores, >= 2 (increased), or <= 0.5 (reduced). The edges were further grouped by “edge type” based on the weight percentile difference, >= 0.5 (stronger), or <= -0.5 (weaker), compared to both WT and the opposite-type K-means clusters. Lastly, they were grouped by “km.cluster type” based on their association with essential or nonessential hub deletion.
